# Transcriptomic profiling of reward and sensory brain areas in perinatal fentanyl exposed juvenile mice

**DOI:** 10.1101/2022.11.18.517129

**Authors:** Jimmy Olusakin, Gautam Kumar, Mahashweta Basu, Cali A. Calarco, Megan E. Fox, Jason B. Alipio, Catherine Haga, Makeda D. Turner, Asaf Keller, Seth A. Ament, Mary Kay Lobo

## Abstract

Use of the synthetic opioid fentanyl increased ∼300% in the last decade, including among women of reproductive ages. Adverse neonatal outcomes and long-term behavioral disruptions are associated with perinatal opioid exposure. Our previous work demonstrated that perinatal fentanyl exposed mice displayed enhanced negative affect and somatosensory circuit and behavioral disruptions during adolescence. However, little is known about molecular adaptations across brain regions that underlie these outcomes. We performed RNA-sequencing across three reward and two sensory brain areas to study transcriptional programs in perinatal fentanyl exposed juvenile mice. Pregnant dams received 10μg/ml fentanyl in the drinking water from embryonic day 0 (E0) through gestational periods until weaning at postnatal day 21 (P21). RNA was extracted from nucleus accumbens (NAc), prelimbic cortex (PrL), ventral tegmental area (VTA), somatosensory cortex (S1) and ventrobasal thalamus (VBT) from perinatal fentanyl exposed mice of both sexes at P35. RNA-sequencing was performed, followed by analysis of differentially expressed genes (DEGs) and gene co-expression networks. Transcriptome analysis revealed DEGs and gene modules significantly associated with exposure to perinatal fentanyl in a sex-wise manner. The VTA had the most DEGs, while robust gene enrichment occurred in NAc. Genes enriched in mitochondrial respiration were pronounced in NAc and VTA of perinatal fentanyl exposed males, extracellular matrix (ECM) and neuronal migration enrichment were pronounced in NAc and VTA of perinatal fentanyl exposed males, while genes associated with vesicular cycling and synaptic signaling were markedly altered in NAc of perinatal fentanyl exposed female mice. In sensory areas from perinatal fentanyl exposed females, we found alterations in mitochondrial respiration, synaptic and ciliary organization processes. Our findings demonstrate distinct transcriptomes across reward and sensory brain regions, with some showing discordance between sexes. These transcriptome adaptations may underlie structural, functional, and behavioral changes observed in perinatal fentanyl exposed mice.

## INTRODUCTION

The opioid epidemic is one of the most severe and intractable public health crises in US history (1). Fentanyl and other potent synthetic opioid analogs, which are 50-100 times more potent than morphine, are commonly added to illicit drugs, increasing fentanyl exposure, and driving opioid-related overdose deaths (1). In 2016, about 11.5 million people reported opioid misuse, out of which 18.2% met criteria for an opioid use disorder (OUD) (https://www.samhsa.gov/data/). Furthermore, prescription opioids among women of reproductive ages have increased over the years posing a public health concern (2). Opioid use during pregnancy predisposes babies to withdrawal symptoms, premature births, decreased birth weight and body size, and, in extreme cases, stillbirth and infant death (3,4). There is also the risk of opioid dependent babies born with neonatal abstinence syndrome. Long term consequences of prenatal and perinatal opioid exposure include dysregulated stress reactivity, altered glucocorticoid levels, hyperactivity, impulsivity, and aggression (5). Perinatal opioid exposure can also produce delayed sensory milestones in preclinical models and human offspring (6,7), as well as abnormal pain sensitivity (8). Opioids cross the placental barrier, entering fetal circulation and the central nervous system where they may affect fetal brain development. Understanding the neuro-molecular signatures and transcriptional substrates resulting from developmental opioid exposure that may mediate long term behavioral phenotypes has been challenging. Preclinical studies describing the molecular and cellular changes in endogenous opioid and neurotransmitter signaling often focus on isolated brain regions and behaviors canonically associated with addiction including the reward and sensory areas (9–11) with little or no emphasis on sex as a variable (12).

Previous models on in utero opioid exposure have focused on prenatal exposure (13) which do not completely encompass human embryonic developmental timeline. We recently developed a mouse model of perinatal fentanyl exposure to mimic and capture the human embryonic developmental exposure period (11,14). At juvenile ages of postnatal day 35 (P35), we showed that perinatal fentanyl exposed mice displayed signs of withdrawal, increased negative affect and an imbalance in synaptic excitation-inhibition within layer 5 pyramidal neurons of the somatosensory cortex (11,14) suggesting a direct action of fentanyl during early development. To understand molecular mechanisms underlying the behavioral and physiological deficits in these juvenile mice, we investigated the transcriptional programs altered in reward-related and sensory brain areas of perinatal fentanyl exposed mice of both sexes.

## METHODS AND MATERIALS

### Experimental subjects

All experiments were performed in accordance with the Institutional Animal Care and Use Committee guidelines at the University of Maryland School of Medicine (UMSOM). C57BL/6 control mice were given food and water *ad libitum* and housed in UMSOM animal facilities on a 12:12 h light:dark cycle.

### Fentanyl administration

Upon determination of pregnancy by presence of vaginal plugs, C57BL/6 mice were singly housed. Each dam was provided with a single 50 mL falcon tube with a rubber stopper and ball-point sipper tube (Ancare) containing 10µg/mL fentanyl citrate (Cayman #22659) in water (15) as their sole liquid source for embryonic day 0 (E0) through gestation till weaning of pups at postnatal day 21 (P21). Control mice received plain water in identical 50 mL tubes. Fentanyl or water consumption was determined by weighing the tubes daily and normalized to individual mouse body weight. After weaning pups at P21, pups were grouped housed with same-sex littermates and given food and water *ad libitum* until P35.

### Tissue collection and RNA extraction

At P35, juvenile mice were weighed prior to tissue collection and four mice/group/sex from mixed population (one animal from one litter) were used. All tissue punches were collected between 10 am and 12 pm as follows, prelimbic cortex (PrL) with a single midline 12-gauge, ventral tegmental area (VTA) with a single midline 14-gauge, nucleus accumbens (NAc) and ventrobasal thalamic nuclei (VBT) with bilateral 14-gauge and primary somatosensory cortex (S1) with bilateral 15-gauge and stored at −80 °C until ready for RNA processing. Total RNA was isolated by using Trizol (Invitrogen) homogenization and chloroform layer separation. RNA was then purified using an RNeasy Mini (#74104, Qiagen) kit manual with a DNase step (#79254, Qiagen). RNA concentration and purity were measured using a Nanodrop (ThermoScientific).

### RNA sequencing and RNAseq data analysis

For RNA sequencing, only samples with RNA integrity numbers >8 were used. Samples were submitted in biological quadruplicates for RNA sequencing at the UMSOM Institute for Genome Sciences (IGS) and processed as described previously (16). Libraries were prepared from 10 ng of RNA from each sample using the SMART-Seq v4 kit (Takara). One VTA male sample (VTA_MW_3 *– see page 9 and 10 of supplementary data 1*) was an outlier and removed from subsequent clustering analysis. Samples were sequenced on an Illumina HiSeq 4000 with a 75 bp paired-end read. 64–100 million reads were obtained for each sample. Reads were aligned to the mouse genome (*Mus musculus*. GRCm38) using TopHat2 (version 2.0.8; maximum number of mismatchesD=DD2; segment lengthD=DD30; maximum multi-hits per readD=DD25; maximum intron lengthD=DD50,000). The number of reads that aligned to the predicted coding regions was determined using HTSeq. We applied two strategies to characterize gene expression changes in these data. First, we identified individual genes with significant gene expression changes following perinatal fentanyl exposure. We used limma-trend to fit log2-normalized read counts per million to a linear model and tested for significant effects of fentanyl in each brain area, separately in males and in females, as well as treating sex as a covariate (*See Supplementary data 1 for methodology of gene analysis*). In our primary analysis, we set a False Discovery Rate (FDR) < 0.05 to define differentially expressed genes. We used a threshold-free approach, Rank-Rank Hypergeometric Overlap (RRHO), to compare effects between connected brain areas. For functional enrichment analysis, we analyzed sets of fentanyl-associated genes with FDR < 0.05 and fold change (FC) +/− 0.3. We also considered a broader set of fentanyl-associated genes with (uncorrected) p-values < 0.05. Analysis of over-represented Gene Ontology (GO) terms were performed with clusterProfiler 4.0 (17) and gprofiler2 for R (18), and Metascape (using default enrichment parameters, http://www.metascape.org) (19) for functional enrichment analysis. iRegulon plugin on Cytoscape 3.0 (20) was used to predict transcription factors, targets and binding motifs. In addition, we characterized the effects of fentanyl on gene co-expression networks. As a starting point for this analysis, we used limma-trend to select the set of genes that exhibited a nominally-significant effect (p-value < 0.05) of fentanyl or sex. We then used weighted gene co-expression network analysis (WGCNA) to characterize co-expressed modules among these genes, separately across all 5 brain areas (*See Supplementary data 4 for methodology of WGCNA analysis*). Module detection was performed with the blockwiseModules() function with power = 10, corType = ‘bicor’, networkType = ‘signed’, minModuleSize = 25, reassignThreshold = 0, mergeCutHeight = 0.25, minMEtoStay = 0, and otherwise default parameters. We tested for differential expression of module eigengenes with limma, using posthoc contrasts to estimate effects of fentanyl while adjusting for sex differences, as above. Hub genes for each module were defined by calculating the Pearson correlation between the eigengene and each gene in the module. For functional enrichment analyses of network modules, we used gprofiler2, in which genes were ranked by their module membership scores, with hub genes ranked highest. We report the most re-occurring enriched terms to describe each module. Only genes and DEGs expressed in brain regions were used to calculate functional enrichment.

## RESULTS

### Perinatal fentanyl exposure induces gene expression changes by sex in reward and sensory brain regions of juvenile mice

Pregnant C57BL/6 dams received 10μg/ml fentanyl in their drinking water from embryonic day 0 (E0) through birth until weaning at postnatal day 21 (P21). Liquid consumption by dams all through gestation to weaning was comparable between water and fentanyl in water group. Post-weaning, at P35, we performed an unbiased transcriptomic analysis in perinatal fentanyl exposed mice using high-throughput RNAseq of bulk-tissue punches from VTA, NAc, PrL, S1 and VBT from four juvenile (P35) mice per sex/treatment (Fig. 1A), (https://www.ncbi.nlm.nih.gov/geo/query/acc.cgi?acc=GSE231981). Prior to tissue collection for RNAseq, weights of juvenile mice were recorded (Fig. S1). We confirmed expression of some tissue specific marker genes within the differentially expressed genes (DEGs) across all brain areas (Fig. S2). Next, we used two independent parallel streams of analyses to identify gene substrates altered by perinatal fentanyl exposed. First analyzing for DEGs using an FDR < 0.05 to identify DEGs across each brain region in a sex-wise manner, and testing for functional enrichments. Second, using two threshold-free methods to (a) evaluate the degree of overlap of gene expression between sexes within brain regions using a Rank Rank Hypergeometric Overlap analysis (RRHO) and (b), a Weighted Gene Correlation Network Analysis (WGCNA) approach which allowed us to cluster genes dysregulated by perinatal fentanyl exposure into modules.

**Figure 1.**
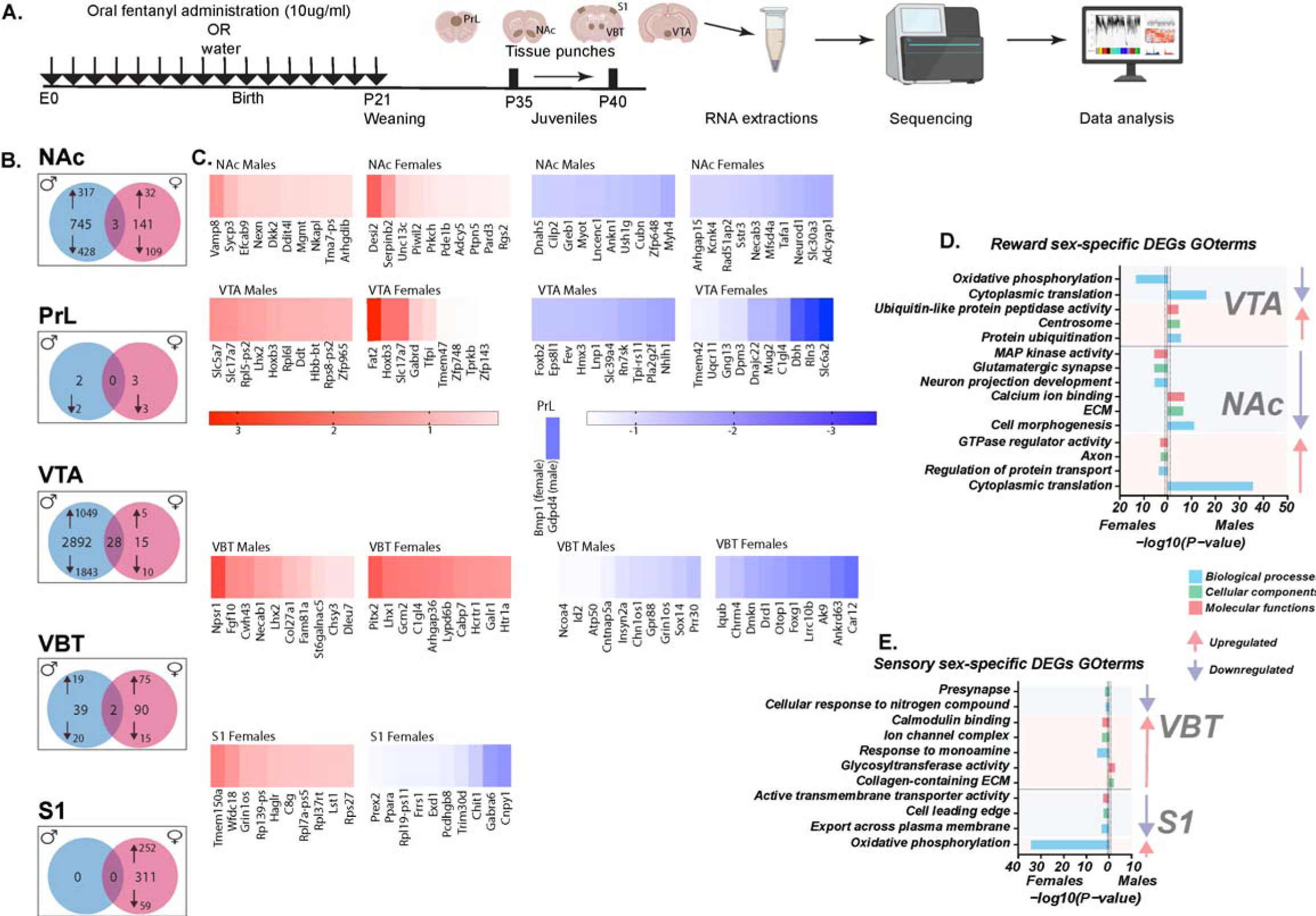
Transcriptomic profiling of reward and sensory brain areas of perinatal fentanyl exposed juvenile mice. (A) Schematic of experimental design and pipeline. Pregnant C57/BL6 mice were administered 10µg/ml fentanyl in drinking water from E0 through birth till weaning of pups at P21. Between P35 and P40, 4 mice/treatment/sex were sacrificed, and tissue punches were collected from PrL with a single midline 12-gauge, NAc and VBT with bilateral 14-gauge, VTA with a single midline 14-gauge and S1 with bilateral 15-gauge. Thereafter, RNAs were extracted from tissue punches and sequenced. Following quality control, subsequent transcriptomic analyses was done with significance set at FDR<0.05. (B) Venn diagram of DE transcripts across brain areas using an FDR of < 0.05 and Log Fold Change (LogFC) cut-off of +/− 0.3. (C) Heatmaps showing top DEGs (upregulated in Red and downregulated in blue) by sex across all brain areas. (D) and (E) Gene Ontology Enrichment analyses on significantly altered perinatal fentanyl exposed DEGs by sex across reward (D) and sensory (E) areas. Blue bars (BP), green bars (CC) and red bars (MF). Y-axis indicates top enrichment terms and X-axis indicates significance level of enrichment. Blue and red arrows indicate expression pattern of enriched terms by sex. E0, embryonic day 0; P21, P35 and P40, postnatal days 21, 35 and 40; PrL, prelimbic cortex; NAc, nucleus accumbens; VBT, ventrobasal thalamus; VTA, ventral tegmental area; S1, somatosensory area 1; DE, differentially expressed; DEGs, differentially expressed genes, BP, biological processes; CC, cellular components; MF, molecular functions.

A total of 2,920 genes were differentially expressed between the perinatal fentanyl exposed and control groups in male and/or female mice in at least one of the five tissues (henceforth, “perinatal fentanyl exposed DEGs”), with FDR < 0.05 (*Supplementary data 2 for full list of perinatal fentanyl exposed DEGs*). We identified more DEGs in the VTA (2,935 DEGs) and NAc (889 DEGs) compared to S1 (311 DEGs), VBT (131 DEGs), or PrL (5 DEGs) (Fig. 1B). In the NAc, we observed *Vamp8(FDR<0.04;LogFC=1.77) and Sycp3(FDR<0.03;LogFC=1.21)* among top upregulated DEGs in males and *Desi2(FDR<0.03;LogFC=2.42), Serpinb2(FDR<0.04;LogFC=1.66)* in females (Fig. 1C). In the VTA, male perinatal fentanyl exposed mice had *Slc5a7(FDR<0.01;LogFC=1.8)* and *Slc17a7(FDR<0.01;LogFC=1.76)* as top upregulated and females had *Fat2(FDR<0.03;LogFC=3.24)* and *Hoxb3(FDR<0.01;LogFC=2.21)* (Fig. 1C). Among top downregulated perinatal fentanyl exposed DEGs in the NAc were *Myh4(FDR<0.02;LogFC=-1.77), Zfp648(FDR<0.003;LogFC=-1.53)* (Fig. 1C) in males and *Adcyap1(FDR<0.04;LogFC=-1.77)*, and *Slc30a3(FDR<0.04;LogFC=-1.73)* (Fig. 1C) in females. In the VTA, downregulated male perinatal fentanyl exposed DEGs included *Nhlh1(FDR<0.002;LogFC=-1.93)* and *Pla2g2f(FDR<0.004;LogFC=-1.88)* (Fig. 1C) while females had *Slc6a2(FDR<0.0006;LogFC=-3.92)* and *Rln3(FDR<0.0003;LogFC=-3.44)* (Fig. 1C). In the PrL, we observed no upregulated perinatal fentanyl exposed DEGs and only two known downregulated DEGs – Gdpd4*(FDR<0.03;LogFC=-2.61)* in males and Bmp1*(FDR<0.02;LogFC=-0.44)* in females (Fig. 1C). In the sensory areas, top upregulated VBT perinatal fentanyl exposed DEGs in males included *Npsr1(FDR<0.003;LogFC=2.74)* and *Fgf10(FDR<0.001;LogFC=1.9)*; while females had *Pitx2(FDR<0.02;LogFC=2.48)* and *Lhx1(FDR<0.01;LogFC=2.17)* (Fig. 1C). Top downregulated perinatal fentanyl exposed DEGs in male VBT included *Prr30(FDR<0.04;LogFC=-1.48)* and *Sox14(FDR<0.03;LogFC=-1.3)* while females had *Car12(FDR<0.01;LogFC=-2.74)* and *Ankrd63(FDR<0.007;LogFC=-2.53)* (Fig. 1C). No perinatal fentanyl exposed DEGs were observed in male S1, however female upregulated DEGs included *Tmem150a(FDR<0.04;LogFC=2.05)* and *Wfdc18(FDR<0.005;LogFC=1.71)* and downregulated DEGs included *Cnpy1(FDR<0.01;LogFC=-2.21)* and *Gabra6(FDR<0.04;LogFC=-2.04)* (Fig. 1C). We observed 28, 3, and 2 perinatal fentanyl exposed DEGs common between both sexes in VTA, NAc, and VBT, respectively, and none in S1 and PrL (Fig. 1B). We also considered genes with trend effects (uncorrected p-values < 0.05 in each sex/tissue) across all of the brain areas by sex. This was done to accommodate for analysis of the PrL since we observed very few DEGs in this brain area with FDR<0.05. In addition, we attempted using uncorrected p values (p<0.05) with the rationale that this would show major differences in functional gene enrichment as compared to using FDR<0.05. However, this was not the case for all analysis in Fig. 1 and S2, FDR<0.05 was used. We present the analysis with the uncorrected p values in Fig. S3.

Heatmaps of common genes with uncorrected P values (*P<0.05*) between male and female juvenile mice across all brain regions showed similar expression profile in PrL, VTA and S1 (Fig. S3). However, in NAc and VBT, many of these common genes were expressed in opposite directions in males vs. females (Fig. S3). In summary, perinatal fentanyl exposure had dramatic effects on gene expression in VTA and NAc by sex.

We performed functional enrichment analyses of perinatal fentanyl exposed DEGs in a sex-wise manner to gain insight into the biological consequences of perinatal fentanyl. Functional enrichment analyses of DEGs suggested that the strongest effects of perinatal fentanyl exposed were on the expression of genes related to mitochondrial respirasome and structural ribosomal constituents (Fig. 1D,E; Fig. S4; *Supplementary Data 3*). In particular, genes related to complex 1 NADH:ubiquinone oxidoreductase (*Nduf*) were differentially expressed in a sex-dependent manner across most of the brain regions assessed, with the strongest effects in the NAc and VTA of males and S1 of female mice. Interestingly, these *Nduf* genes were upregulated in male NAc and female S1 but were downregulated in the VTA of both sexes (Table S1A). This could indicate an increase in active protein synthesis and cellular energy production in S1 and NAc of females and males respectively while a decrease in cellular energy production in the VTA.

Perinatal fentanyl exposed was also associated with gene expression changes in pathways with functions more specific to neurons. We observed downregulation of collagen type (*Col*) genes specifically in males in both NAc and VTA (Fig. 1D, Table S1B). These collagens are components of the extracellular matrix (ECM) and play a critical role in regulation of myriad cellular and structural functions ranging from cell migration, angiogenesis to inflammation (21,22). In NAc, perinatal fentanyl exposed was also associated with down-regulation of synaptic and vesicular signaling genes, primarily in females (Table S1C).

### Sex effects of perinatal fentanyl exposure are most visible in the NAc and VBT of juvenile mice

Next, we sought to compare the effects of perinatal fentanyl exposed on transcriptional landscape using a threshold free analysis between sexes across brain regions. For this purpose, we performed a two-sided rank-rank hypergeometric overlap analysis (RRHO) to identify patterns and strength of genome-wide overlap (Fig. 2A). Heatmap expression profile from RRHO analyses showed concordance (lower left and upper right quadrants) overlap of perinatal fentanyl exposed genes in PrL (Fig. 2B,B’), VTA (Fig. 2D,D’) and S1 (Fig. 2E,E’). However, NAc (Fig. 2C,C’) and VBT (Fig. 2F,F’) had the majority of discordant (upper left and lower right) perinatal fentanyl exposed genes. We reasoned that the discordant gene clusters would show sex differences in gene substrates altered by perinatal fentanyl exposed. Functional enrichment analysis of discordant genes in the NAc revealed upregulated perinatal fentanyl exposed male genes and downregulated female genes were enriched in Mitochondrial protein complex (Biological processes), oxidative phosphorylation (Cellular components) and Proton-transmembrane transporter activity (Molecular function) (Fig. 2C’). However, downregulated perinatal fentanyl exposed male genes and upregulated female genes were enriched in Morphogenesis (Biological processes), Cilium (Cellular components) and ATP-dependent chromatin remodeler activity (Molecular function) (Fig. 2C’). In the VBT, upregulated perinatal fentanyl exposed male genes and downregulated female genes were enriched in Cilium movement (Biological processes), Cilium (Cellular components) and Extracellular matrix structural constituent (Molecular function) (Fig. 2F’). Downregulated perinatal fentanyl exposed male genes and upregulated female genes were enriched in Trans-synaptic signaling (Biological processes), Presynaptic membrane (Cellular components) and Neurotransmitter receptor activity (Molecular function) (Fig. 2F’).

**Figure 2.**
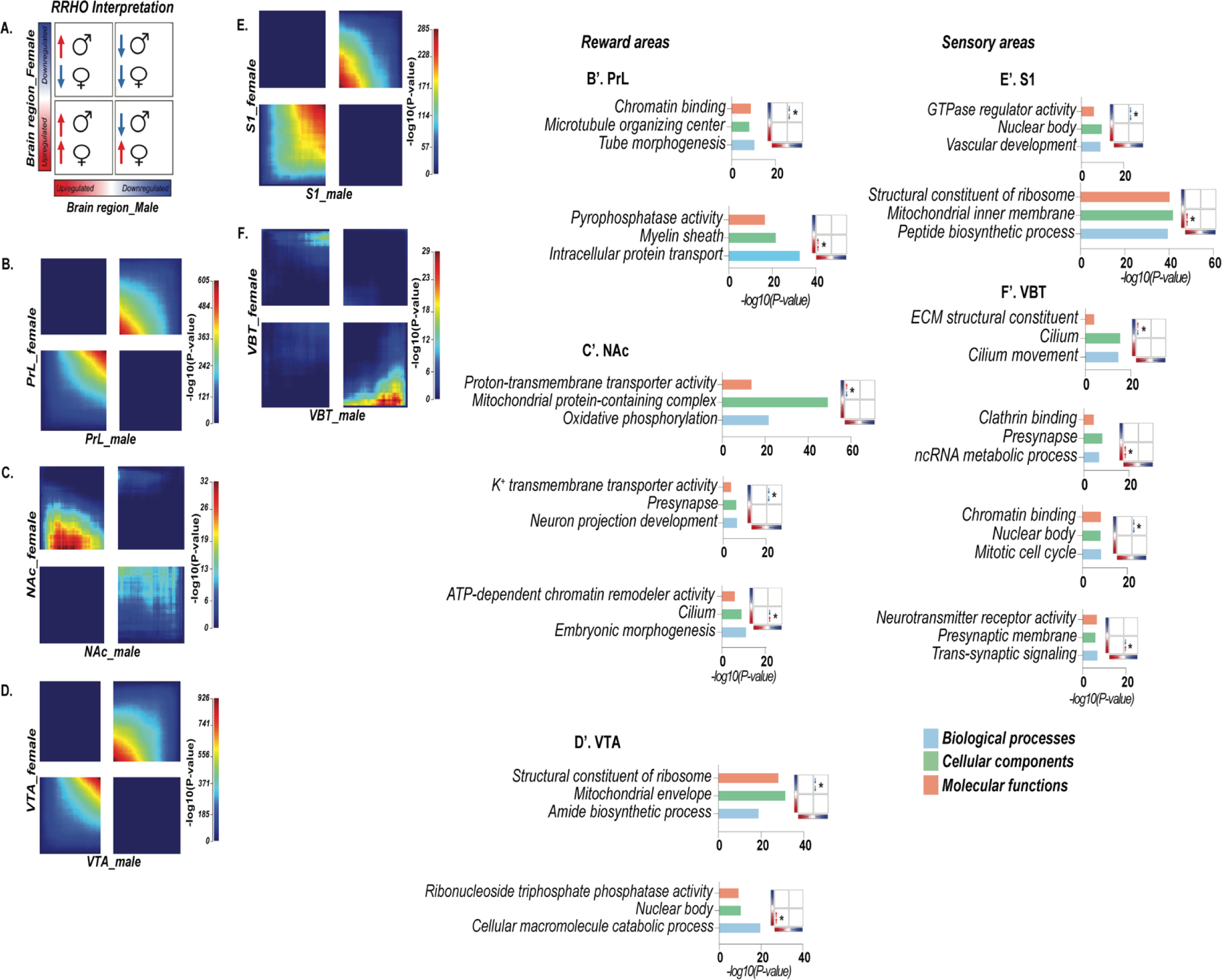
Threshold-free analysis of concordant and discordant gene overlap between sexes across brain areas in perinatal fentanyl exposed mice. (A) Schematic representation of heat maps on RRHO plots. (B) and (B’) Heat map of PrL genes and functional enrichment of concordant genes. (C) and (C’) Heat map of NAc genes and functional enrichment of concordant and discordant genes. (D) and (D’) Heat map of VTA genes and functional enrichment of concordant genes. (E) and (E’) Heat map of S1 genes and functional enrichment of concordant genes. (F) and (F’) Heat map of VBT genes and functional enrichment of concordant and discordant genes. Blue bars (BP), green bars (CC) and red bars (MF). Y-axis indicate top enrichment and X-axis indicate significance level of enrichment. RRHO small squares orient to quadrant enrichment using (A) as a guide. PrL, prelimbic cortex; NAc, nucleus accumbens; VBT, ventrobasal thalamus; VTA, ventral tegmental area; S1, somatosensory area 1; BP, biological processes; CC, cellular components; MF, molecular functions.

### Network analyses reveals sex-wise transcriptional signatures correlated with perinatal fentanyl exposed

To better resolve gene clusters associated with fentanyl in a sex-wise manner, and define functional enrichment by modules, we performed Weighted Gene Co-expression Network Analysis (WGCNA, *Supplementary data 4*) for each brain region separately, identifying 10 to 46 gene co-expression modules per brain region (*Supplementary data 4, page 6-8*). We used linear models to test the sex associations of each module with fentanyl treatment. We identified 7 fentanyl-associated modules in the NAc, 8 in the PrL and 6 in the VTA (Fig. 3A,B). S1 and VBT each had 7 fentanyl-associated modules (Fig. 4A,B). Gene set enrichment analyses identified functional categories enriched in many of these modules (Fig. 3C and Fig. 4C). In the NAc and VTA,

**Figure 3.**
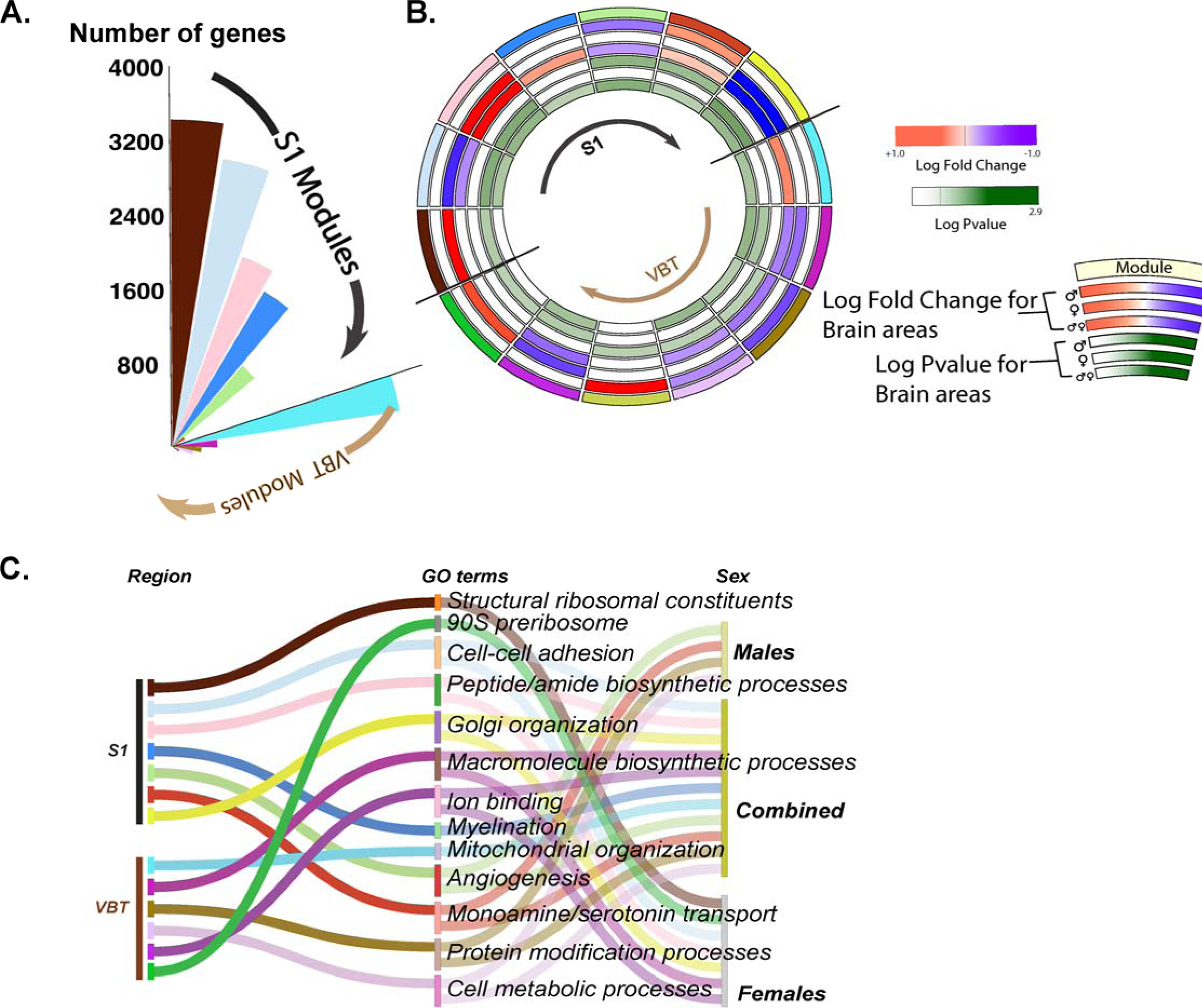
Weighted gene co-expression network analyses (WGCNA) showing significant modules associated with fentanyl in a sex-wise manner in reward brain areas. (A) Circular bar plot showing significant modules associated with fentanyl. Each bar on plot is arbitrarily color coded to denote a module. The size of each bar represents the number of genes within the module. (B) Circos plot from WGCNA of modules with significant differential expression (fentanyl ∼ water). Modules were built using the average expression of the eigen genes. On circus plot, modules are shown in a clockwise fashion from PrL, VTA and NAc. Each pie represents a module with outermost segment retaining module color, and inner 3 segments containing fold change in expression for male, female and common genes associated with fentanyl respectively, and innermost 3 segments representing −log_10_(*P* values) respectively. (C) Sankey plot showing description of top gene ontology enrichment for each significant module.

**Figure 4.**
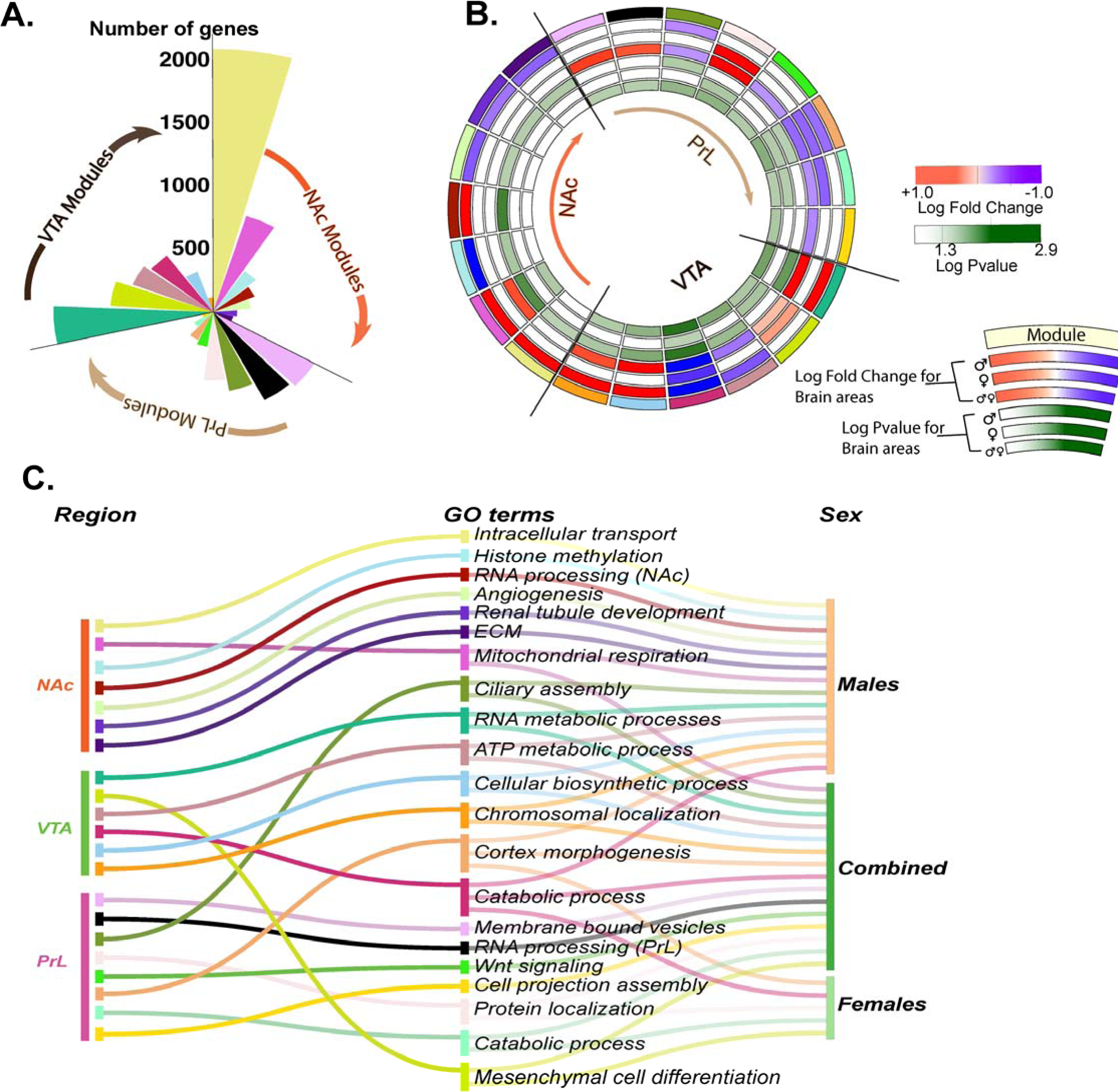
WGCNA showing significant modules associated with fentanyl in a sex-wise manner in sensory brain areas. (A) Circular bar plot with significant modules associated with fentanyl. Each bar on plot is arbitrarily color coded to denote a significant module. The size of each bar represents the number of genes within the module. (B) Circos plot showing modules in a clockwise fashion from S1 to VBT. Each pie represents a module with outermost segment retaining module color and inner 3 segments containing fold change in expression for male, female and common genes associated with fentanyl respectively, and innermost 3 segments representing −log_10_(*P* values) respectively. (C) Sankey plot showing description of top gene ontology enrichment for each significant module.

11 out of the 13 significant modules contained genes positively associated with perinatal fentanyl exposed only in males (Fig. 3B). All PrL significant modules showed positive association with fentanyl when sexes were combined. Five of 8 modules had downregulated genes – one module each, “turquoise” and “green” were specifically driven by gene expression changes in females and in males, respectively (Fig. 3B). Three modules in the PrL contained upregulated genes, with one association driven specifically by gene expression changes in females.

In reward brain regions, we identified histone methylation, angiogenesis, mitochondria (Fig. 3C) – processes described to be dysregulated in adult opioid use disorder (23,24) also dysregulated in perinatal fentanyl exposed juvenile mice. Interestingly, we found in the NAc “hot pink” module the highest enrichment for mitochondrial respiration and structural ribosomal terms specifically in upregulated male genes. Angiogenesis was enriched in downregulated male genes in NAc “light green” module (Fig. 3C). In the PrL, although no obvious enrichment was seen with regional analyses, modular analyses revealed significant enrichment in ciliary assembly and neuronal projection morphogenesis. In the VTA, the only module associated with perinatal fentanyl exposed female upregulated genes was enriched in mesenchymal cell differentiation (Fig. 3C). The remaining 5 modules were enriched in metabolic processes.

In the S1 cortex there was a greater association of fentanyl with genes differentially expressed in females. Specifically, 4 of 7 modules contained female genes – 2 contained upregulated and 2 contained downregulated genes (Fig 4B). The VBT, however, had equal number of sex specific modules associated with fentanyl. Structural ribosomal constituent was enriched in upregulated female modular “brown module” in S1 (Fig. 4C). Other terms of interest that were revealed by the enrichment analyses in S1 were myelination, monoamine/serotonin transport and Golgi organization (Fig. 4C).

Since our GO term analysis suggested common recurrent functional enrichment processes altered by perinatal fentanyl exposed, we then sought to find potential transcriptional regulators for these processes (Fig. 5). We used the transcriptional regulator analysis iRegulon on genes from enriched GO terms across reward (Fig. 5A) and sensory (Fig. 5B) regions to identify the top 5 predicted transcriptional regulators based off their normalized enrichment score (NES). We identified 2 zinc finger transcription factors predicted to regulate genes associated with metabolic processes and intracellular transport across the reward areas (Fig. 5A). Specifically, *Zfp143* (Zinc finger protein 143), a transcriptional activator, is predicted to regulate target genes involved in intracellular transport, mitochondrial respiration, ATP metabolic processes across all three reward areas (Fig. 5A). *Yy2* (Yin yang 2 or zinc finger protein 631), a transcription factor which possess both activation and repression domains regulates genes involved in RNA processing across all reward areas (Fig. 5A). Ebf1 (Early B-Cell factor 1) essential for lineage specification in early B cell development had the highest NES predicted to regulate genes enriched in angiogenesis in the NAc (Fig. 5A). We identified *Mef2c* (Myocyte enhancer factor 2C) and *Zmat2* (Zinc finger matrin-type 2) (Fig. 5A) transcription factors with highest NES in processes involved in ciliary organization andIn sensory modules, we identified Elk4 (Serum response factor accessory protein 1) and Mapk1 (Mitogen-activated protein kinase 1) as top predicted transcription factors to regulate mitochondrial organization and blood vessel development genes respectively in S1 (Fig. 5B). In the VBT, we identified Atf7 (Activating transcriptor factor 7) and Zfp110 (Zinc finger protein 110 – an ortholog to human zinc finger protein 274) as top regulators of protein modification and cellular metabolic process genes respectively (Fig. 5B).

**Figure 5.**
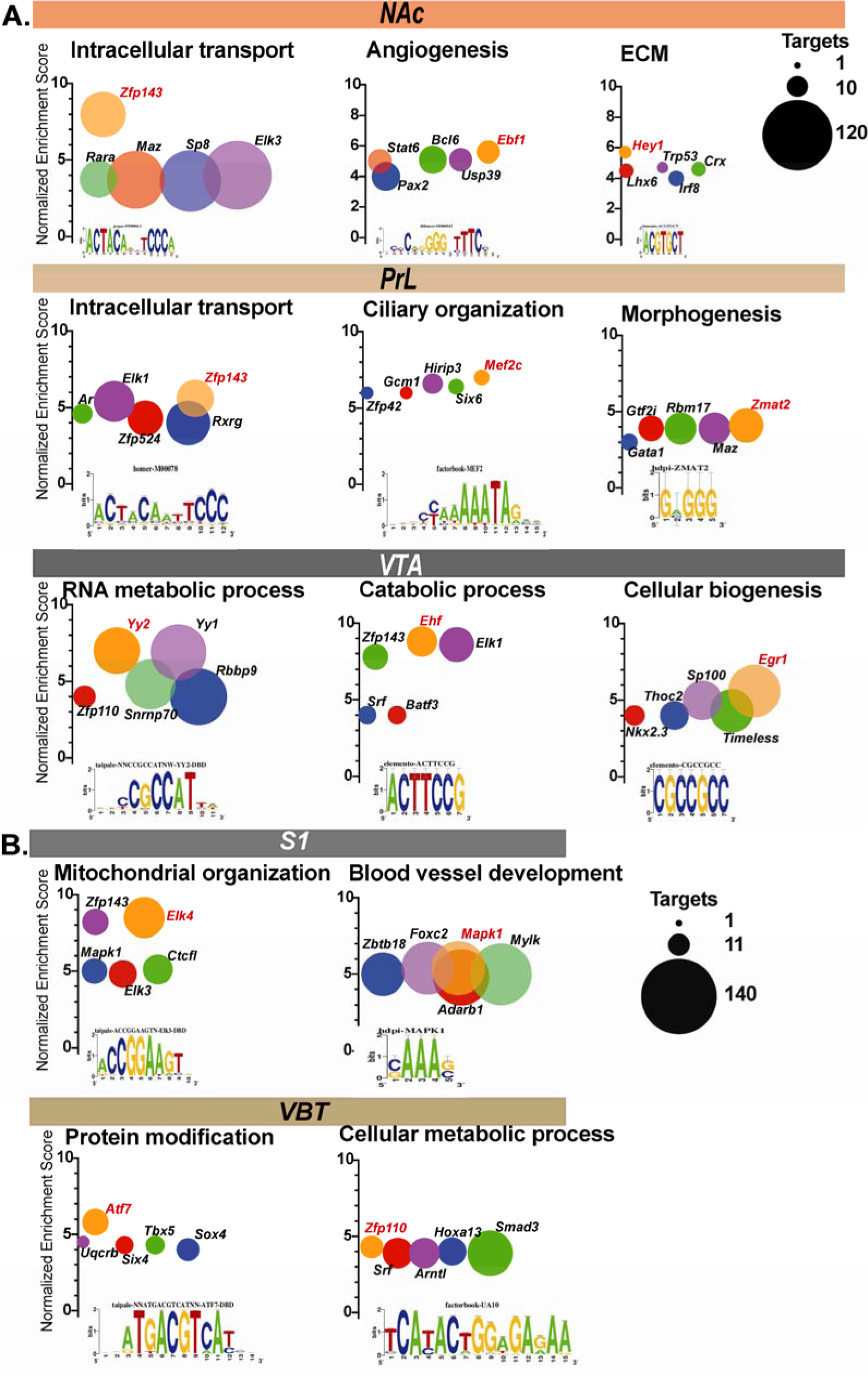
Transcription factors (TFs) predicted with iRegulon for selected biological processes altered in reward (A) and sensory (B) brain areas of perinatal fentanyl exposed mice. Bubble plots show top five TFs predicted to regulate enriched biological processes within modules across all five brain areas. On the Y-axis are the normalized enrichment scores (NES) for each TF. Size of each bubble indict number of downstream target genes regulated by TF. The insert show predicted binding motifs of the most enriched TF for a particular biological process highlighted in red.

## DISCUSSION

In this study, we present a comprehensive transcriptional landscape of genes dysregulated by sex in juvenile mice exposed perinatally to fentanyl. Pregnant mice were administered 10µg/ml fentanyl in the drinking water throughout gestation until weaning. While daily forced administration through drinking could potentially be a stressor, this route minimizes stress from daily handling and injections. In addition, mice will readily self-administer 10µg/ml fentanyl in drinking water without any motor deficits (11,14,25). Further, this dose is consistent with our previous studies demonstrating brain and behavioral adaptations which led us to focus on reward and sensory brain regions (11,14). The novelty in our study describes distinct transcriptional programs and altered biological processes in three reward areas – NAc, PrL and VTA and two sensory areas - S1 and VBT. Further, we demonstrate unique sex selective patterns of gene expression including discordance of overlapping genes between the sexes in NAc and VBT regions. The perinatal fentanyl exposed procedure we used was based on an established model (11,14) that displays negative affective behaviors, such as heightened anxiety-like behaviors and reduced grooming behavior, or sensory deficits, including defective sensory adaptation (11) and reduced pain sensitivity behaviors (8) at juvenile ages. We previously reported that despite having lower birth weights, at juvenile ages, perinatal fentanyl exposed mice show higher body weights compared to their control counterparts (14). Indeed, we noted that juvenile mice used for RNAseq in this study had higher body weight, though not significant compared to their control littermates.

### Perinatal fentanyl exposure alters synaptic, cell adhesion and vesicular transporter genes in a sex-wise manner

We used two parallel analysis streams and show convergence between these analyses. Our primary DEG analyses showed that the VTA had the greatest number of DEGs regulated by perinatal fentanyl exposure. The VTA has a fundamental role in opioid dependence and withdrawal, including neuronal structural and functional plasticity particularly elicited by opioid exposure, which could explain the high transcriptional substrates we observed in our perinatal fentanyl data. The prefrontal cortex including its PrL region undergoes a protracted development and substantial remodeling throughout adolescence (26). Previous studies have observed attenuated neuronal activity in the PrL of adolescent rats following heroin self-administration (27). In this study, we did not observe substantial DEGs regulated by perinatal fentanyl exposure in the PrL of both sexes probably due to a number of factors – (1) due to the protracted development and dynamic changes occurring in this brain area, there might not be quantifiable significant effect of perinatal fentanyl exposure, (2) variability in gene expression among biological samples. However, we observed *Gdpd4* – Glycerophosphodiester phosphodiesterase domain containing 4 to be downregulated in PrL of perinatal fentanyl exposed male mice. *Gdpd4* is one of five family of *Gdpd* and strongly expressed in testes of male mice. It has been reported that no overt developmental or fertility abnormalities in Gdpd4 mutant mice (28), however, inherited or de novo copy number variants encompassing Gdpd4 were found in human patients with autism (29). In perinatal fentanyl exposed females, downregulation of bone morphogenetic protein 1 (*Bmp1*) could indicate abnormal maturation of diverse ECM proteins as well as precocious activation of TGF-ß related growth factors (30). Synaptic adaptations within the mesolimbic reward areas and associated neural circuits persists long after drug cessation which may influence behavior. Previous studies have reported altered synaptic communication involving components of the SNARE complex remained altered well into the abstinence period and thus could contribute to relapse liability (31–33). In line with these reports, we observed that in perinatal fentanyl exposure, male juvenile mice top upregulated DEGs were *Vamp8*, *Sycp3* and *Slc5a7* and *Slc17a7* in the NAc and VTA respectively. Several studies support the role of cell adhesion molecules as well as the involvement of proteases in opioid reward. Serpins are inhibitors of neuronal proteases involved in diverse functions including synaptic plasticity, inflammation and axonal development (34). A previous study found *Serpind1* downregulation in the brainstem of mice withdrawing from chronic postnatal morphine exposure (35). While we did not observe any changes in *Serpind1* expression from perinatal fentanyl exposure, across the reward areas, we observed *Serpinb2* as a top upregulated DEG in the female NAc. In the female VTA, top upregulated DEGs were *Fat2*, *Hoxb3* involved in cell adhesion and developmental processes.

We observed transcription factors *Zfp648* and *Nhlh1* as top downregulated DEGs in the NAc and VTA of juvenile perinatal fentanyl exposed male mice respectively. While not much is documented on the role of both transcription factors in the context of opioid use disorder, some studies have reported their implication in neuronal differentiation and stress regulation (36,37). While Nhlh1 expression appears to be restricted to the nervous system, Cogliati et al showed that Nhlh1-null mice exhibited altered parasympathetic tone and stress-induced arrhythmia (36). Dysregulation of these transcription factors could contribute to the heightened anxiety-like behaviors we previously observed in perinatal fentanyl exposed male juvenile mice (14). In females, we observed *Adcyap1*, *Slc30a3* and *Slc6a2*, *Rln3* as top downregulated DEGs in the NAc and VTA of juvenile perinatal fentanyl exposed female mice respectively. Adcyap1 otherwise known as *Pituitary Adenylate Cyclase-Activating Polypeptide* or *PACAP* has been reported to modulate anxiety-like behaviors in mice (38–39) and more recently *PACAP* gene polymorphisms was described to increase the risk of developing methamphetamine addiction (40). Dysregulation of zinc homeostasis within the CNS (41) as well as zinc deficiency has been implicated in mood and other negative affective states (42–44). In line with this, our observation of decreased expression of the zinc transporter – *Slc30a3* in the NAc of female perinatal fentanyl exposed mice could account for the deficit in grooming behaviors we previously observed (14). We also found the norepinephrine transporter – *Slc6a2* significantly downregulated in the VTA of female perinatal fentanyl exposed mice.

The VBT appears to be the more impacted by perinatal fentanyl exposure as compared to the S1 cortex. We observed transcription factors *Pitx2* and *Lhx1* as top upregulated DEGs in the VBT of female perinatal fentanyl exposed mice while in the males, we observed *Sox14*, transcription factor downregulated in VBT. All three transcription factors have been implicated in neuronal migration and regulation of embryonic development (45–47) and specifically in differentiation, migration and maintenance of GABAergic interneurons (48). This suggest that perinatal fentanyl exposure could alter the developmental trajectory of GABAergic interneurons which could persist at least until juvenile ages. In accordance with this, a recent study by Nieto-Estévez et al., reported that prenatal buprenorphine exposure altered interneuron distribution and these alterations persisted into adulthood (49).

### High degree of discordant genes between sexes in NAc and VBT in perinatal fentanyl exposed juvenile mice

We reasoned that discordant gene overlap between sexes across brain areas will give a clearer pattern of sex differences in transcriptional programs dysregulated by perinatal fentanyl exposure. Our threshold-free ranking revealed that the NAc and VBT were the two brain areas with strongest impact of perinatal fentanyl exposure by sex. Specifically, we found a high degree of discordance in the expression profile of genes between sexes in these two areas. In the NAc, driving this discordance were genes enriched in mitochondrial respiration, transmembrane transporter activity, morphogenesis and blood vessel development. When we cross-referenced the genes from NAc RRHO enrichment with NAc DEGs (*FDR < 0.05*) we observed upregulation of complex 1 (*Nduf*) mitochondrial genes in perinatal fentanyl exposed males. This is suggestive of higher energy demands in the NAc of perinatal fentanyl exposed juvenile male mice. Mitochondrial dynamics is key to drug seeking and abstinence. Our lab previously showed increase in mitochondria fission related to excitatory synaptic function in NAc D1-MSNs and drug seeking behavior of mice that self-administered cocaine (50). While we cannot directly determine if these changes in mitochondrial respiratory genes reflects mitochondrial dysfunction or a higher metabolic demand, we observed dual specificity phosphatase 14 (*Dusp14)*, an immediate early gene (IEG) that is regulated in response to oxidative damage, to be upregulated in NAc of perinatal fentanyl exposed males. In line with this, Wojcieszak and colleagues found *Dusp14* among IEGs upregulated in mice striatum in response to acute synthetic cathinone administration (51). In addition, we found the cytosolic antioxidant, Glutathione S-transferase mu 5 (*Gstm5*), to be upregulated in NAc of males, which could reflect a possible response to the increase in oxidative stress. Similar changes in these mitochondrial oxidative stress genes were observed in the striatum of mouse model of Parkinson’s disease (52).

Genes enriched in morphogenesis and blood vessel development were also discordantly regulated in the NAc. Specifically, these genes were found to be downregulated in the males. Morphogenesis enrichment included genes like *Notch 1* and *2, Adgrg1, Lama5*, known to influence, the extracellular matrix, and cell-to-cell signaling. The ECM provides proper structural stability in synaptic function, blood-brain barrier integrity and cell-to-cell communication (22,53). ECM macromolecules are essential for crafting the cellular domains required during development and morphogenesis (19). Proper ECM development and signaling is necessary in setting up brain architecture, opioids impact ECM and sexual dimorphism in ECM has been observed across species (22), thus consideration of ECM in brain development during opioid exposure of both sexes is paramount. Furthermore, we observed downregulation of growth factor receptors – *Fgfr3, Lrp1, Egfr* and collagen type IV genes – *Col4a1 and Col4a2* in NAc of male perinatal fentanyl exposed mice. *Lrp1* and the receptor tyrosine kinases, *Fgfr3* and *Egfr* have been described to have important roles in proliferation, migration and differentiation of vascular endothelia cells (54,55). Downregulation of these receptors could suggest a compensatory mechanism from chronic exposure to receptor ligands or fentanyl exposure. The latter may be the case since mRNA expression levels of receptor ligands (*Egf* and *Fgfs*) are not significantly changed. A recent postmortem study by Mendez et al., found these angiogenic genes including vascular endothelia basement membrane genes *Col4a1* and *Col4a2* upregulated in the PFC of human polysubstance users (56). While we did not find any of these genes significantly affected in the PrL, we cannot directly compare our findings to theirs, we found vascular genes constituently downregulated in the NAc and VTA of male perinatal fentanyl exposed mice.

In the VBT, sex discordant genes were mostly enriched in structural constituent of ECM and trans-synaptic signaling. Most synaptic and presynaptic membrane genes with FDR<0.05 were particularly upregulated in the VBT of perinatal fentanyl exposed females and with a trend towards downregulation (uncorrected p<0.05) in male juvenile mice. Among upregulated female perinatal fentanyl exposed genes are *Gabra5*, *Hcn1*, *Doc2a* and the *G_i_*-coupled 5-HT receptors – *Htr1a* and *Htr1f*. We observe that most of the upregulated genes in the VBT of females encode membrane proteins involved in neuronal hyperpolarization or signaling casade resulting in inhibition of adenylate cyclase suggesting perinatal fentanyl exposure modulates the excitability (probably towards an inhibition) of thalamo-cortico-thalamic network.

### Network analysis reveals a strong association with perinatal fentanyl exposure in the nucleus accumbens of male juvenile mice

WGCNA which takes advantage of some of the pitfalls of DEG analysis to highlight higher-order relationships among genes revealed significant modules associated with perinatal fentanyl exposure. In the reward brain areas, we observed all seven significant modules in the NAc were altered in the male group while no changes were observed in the female group. The lime green module in the NAc had the greatest number of genes (2070) with *Rabac1*, *Clta* and *Atp5d* as top hub genes. *Rabac1* and *Clta* are involved in vesicle formation and intracellular trafficking while *Atp5d* is a subunit of mitochondrial ATP synthase. It was therefore not surprising to have Intracellular transport as the highest gene enrichment for this module. This is in line with the upregulation of mitochondrial respiratory DEGs we observed in the NAc of male perinatal fentanyl exposed mice. In the sensory brains, the dark brown module in S1 had the greatest number of genes (3502) and this module was biased in favor of female perinatal fentanyl exposed group. This module had *Dtx3*, *Enkd1* and *Epn1* as top hub genes. All three hub genes have are described to be involved in proper cytoskeletal reorganization as well as regulating proper intracellular transport (57–59).

Our iRegulon analysis was an in-silico approach to predict transcriptional factors regulating functional categories over-represented within module genes. We found in the reward brain areas, iRegulon predicted transcription factors seem to play vast regulatory roles during brain development (60,61) and gene regulatory roles for functional categories within our analysis. Specifically, we found *Zfp143* (Staf, selenorysteine RNA gene transcription activating factor) consistently enriched for mitochondrial and intracellular transport across brain regions. Although *Zfp143* expression was present in our data across all brain regions, it did not meet our DEG filtering criteria. Investigations on Zfp143 has mainly focused on its transcriptional changes following ethanol consumption (62,63). An understanding of how changes in its expression level and activity modulates developmental opioid consumption warrants further investigation.

In conclusion, our findings elucidate that perinatal fentanyl exposure induces sex-wise transcriptional alterations in reward and sensory brain areas with marked changes in the NAc, VTA and S1 areas. Perinatal fentanyl exposed male juvenile mice showed the most transcriptional perturbations specifically in processes relating to mitochondrial respiration, ECM, and neuronal guidance. Perinatal fentanyl exposed female mice showed transcriptional alterations in genes enriched for vesicular and synaptic signaling processes. The altered functional enrichment we observe are consistent with opioid induced gene enrichment described in the literature. We highlight that these altered processes have a developmental root and as such early intervention beginning precisely at the early stages of abstinence may prevent or reduce some of these neuro-molecular changes and possibly behavioral adaptations.

## Acknowledgements

This work was supported by NIH R01DA054905 (to MKL, SAA, and AK), R01DA038613 (to MKL), University of Maryland Strategic Partnership Empowering the State, and University of Maryland School of Medicine (UMSOM) Center for Substance Use in Pregnancy. The authors also thank the UMSOM Institute for Genome Sciences for RNA-seq services.

## Author contributions

CAC, MEF, JAB, CH and MDT assisted with experiments and tissue collection. SAA, MB, JO and GK analyzed data. JO, MEF, MKL, AK and SAA oversaw experimental design. JO and MKL oversaw data interpretation and writing the manuscript. MKL conceived and directed the project. All authors reviewed and edited the manuscript.

