## Supplementary data 1 for "Transcriptomic profiling of reward and sensory brain areas in perinatal fentanyl exposed juvenile mice"

### Report Developmental Fentanyl Exposure

#### Analysis of Differential gene expression analysis for Developmental Fentanyl Exposure

```
library(knitr)
opts_chunk$set(tidy.opts=list(width.cutoff=50),tidy=TRUE)
```

```
workdir = "/local/projects/idea/mbasu/XLOBO/dev_fentanyl/plot1"
options(stringsAsFactors = F)
library(data.table)
library(edgeR)
library(limma)
library(GO.db)
library(RColorBrewer)
library(gplots)
library(knitr)
library(kableExtra)
library(dplyr)
# library(png)
library(VennDiagram)
library(gridExtra)
library(gplots)
library("devtools")
source("/local/projects/idea/mbasu/software/heatmap.3.R")
library("xlsx")
library(r2excel)
library(biomaRt)
library("org.Mm.eg.db")
options(knitr.table.format = "markdown")
```

##### Count matrix

```
pipeline_id = 13999837338
folder_counts = paste0("/local/projects/RNASEQ/XLOBO/rnaseq/ergatis/output_repository/htseq/",
    pipeline_id, "_exon_counts/i1")
setwd(folder_counts)

htseq.counts.files = Sys.glob("g*/*counts")
file.names = unlist(lapply(1:length(htseq.counts.files),
    function(i) {
        unlist(strsplit(htseq.counts.files[i], "\\/"))[2]
    }))
sample.names.full = gsub(".accepted_hits.sorted_by_name.exon.counts",
```

```r
    "", file.names)

nc = length(sample.names.full)

sample.names = unlist(lapply(1:nc, function(i) {
    x = unlist(strsplit(sample.names.full[i], "_"))
    l = length(x)
    paste(x[1], paste(substr(x[l - 2], 1, 1), substr(x[l -
        1], 1, 1), sep = ""), x[l], sep = "_")
}))

tmp = unique(unlist(lapply(1:length(htseq.counts.files),
    function(i) {
        length(readLines(htseq.counts.files[i]))
    })))
if (length(tmp) == 1) {
    nr = tmp - 5
    print("Number of transcripts across the sample match")
} else {
    print("Mismatch of transcripts across the sample")
}
```

```
## [1] "Number of transcripts across the sample match"
```

```r
nr = nrow(read.table(htseq.counts.files[1])) - 5

counts = matrix(NA, nr, nc)
for (i in 1:nc) {
    tmp = read.table(htseq.counts.files[i], nrow = nr)
    if (i == 1)
        rownames(counts) = tmp[, 1]
    counts[, i] = tmp[, 2]
}
colnames(counts) = sample.names

# Count matrix dimention `r dim(counts)`
# row:transcripts X column:samples
```

##### Meta file

```r
setwd(workdir)
tmp = strsplit(sample.names.full, split = "_")
celltype = sex = trt = biorep = rep(NA)
for (i in 1:nc) {
    l = length(tmp[[i]])
    celltype[i] = tmp[[i]][1]
    sex[i] = ifelse(any(grep("Male", tmp[[i]])), "M",
        "F")
    trt[i] = ifelse(any(grep("Water", tmp[[i]])), "W",
        "F")
    biorep[i] = tmp[[i]][l]
}
```

```
meta = data.frame(sample.names.full, celltype, sex,
    trt, biorep)
meta$Group = paste(meta$celltype, paste(meta$sex, meta$trt,
    sep = ""), sep = "_")
counts.allsampl = counts
meta.allsampl = meta
```

Table 1: Meta file for Fentanyl

| sample.names.full | celltype | sex | trt | biorep | Group |
|---|---|---|---|---|---|
| VBT_Male_Water_3 | VBT | M | W | 3 | VBT_MW |
| NAc_Female_Water_5 | NAc | F | W | 5 | NAc_FW |
| S1_Female_Fentanyl_4 | S1 | F | F | 4 | S1_FF |
| NAc_Female_Water_3 | NAc | F | W | 3 | NAc_FW |
| S1_Male_Fentanyl_2 | S1 | M | F | 2 | S1_MF |
| S1_Female_Water_3 | S1 | F | W | 3 | S1_FW |
| S1_Male_Fentanyl_5 | S1 | M | F | 5 | S1_MF |
| PrL_Male_Fentanyl_3 | PrL | M | F | 3 | PrL_MF |
| VBT_Male_Fentanyl_2 | VBT | M | F | 2 | VBT_MF |
| VTA_Male_Water_1 | VTA | M | W | 1 | VTA_MW |
| S1_Female_Water_5 | S1 | F | W | 5 | S1_FW |
| VBT_Male_Water_5 | VBT | M | W | 5 | VBT_MW |
| VTA_Male_Water_2 | VTA | M | W | 2 | VTA_MW |
| PrL_Male_Fentanyl_4 | PrL | M | F | 4 | PrL_MF |
| NAc_Female_Fentanyl_4 | NAc | F | F | 4 | NAc_FF |
| VBT_Male_Water_1 | VBT | M | W | 1 | VBT_MW |
| PrL_Female_Fentanyl_5 | PrL | F | F | 5 | PrL_FF |
| S1_Male_Water_4 | S1 | M | W | 4 | S1_MW |
| S1_Female_Fentanyl_5 | S1 | F | F | 5 | S1_FF |
| NAc_Male_Fentanyl_4 | NAc | M | F | 4 | NAc_MF |
| VTA_Male_Fentanyl_1 | VTA | M | F | 1 | VTA_MF |
| PrL_Female_Fentanyl_2 | PrL | F | F | 2 | PrL_FF |
| VTA_Female_Water_4 | VTA | F | W | 4 | VTA_FW |
| PrL_Female_Fentanyl_3 | PrL | F | F | 3 | PrL_FF |
| VBT_Male_Fentanyl_5 | VBT | M | F | 5 | VBT_MF |
| VTA_Female_Fentanyl_1 | VTA | F | F | 1 | VTA_FF |
| S1_Female_Water_2 | S1 | F | W | 2 | S1_FW |
| NAc_Female_Water_1 | NAc | F | W | 1 | NAc_FW |
| VBT_Male_Water_2 | VBT | M | W | 2 | VBT_MW |
| NAc_Female_Fentanyl_2 | NAc | F | F | 2 | NAc_FF |
| PrL_Female_Water_2 | PrL | F | W | 2 | PrL_FW |
| NAc_Male_Fentanyl_1 | NAc | M | F | 1 | NAc_MF |
| VTA_Male_Fentanyl_2 | VTA | M | F | 2 | VTA_MF |
| S1_Male_Water_3 | S1 | M | W | 3 | S1_MW |
| VBT_Female_Water_5 | VBT | F | W | 5 | VBT_FW |
| PrL_Female_Water_5 | PrL | F | W | 5 | PrL_FW |
| VBT_Female_Water_1 | VBT | F | W | 1 | VBT_FW |
| PrL_Female_Fentanyl_1 | PrL | F | F | 1 | PrL_FF |
| VBT_Female_Fentanyl_5 | VBT | F | F | 5 | VBT_FF |
| PrL_Male_Fentanyl_2 | PrL | M | F | 2 | PrL_MF |
| NAc_Female_Fentanyl_3 | NAc | F | F | 3 | NAc_FF |
| VTA_Female_Water_2 | VTA | F | W | 2 | VTA_FW |
| S1_Female_Fentanyl_2 | S1 | F | F | 2 | S1_FF |

| sample.names.full | celltype | sex | trt | biorep | Group |
|---|---|---|---|---|---|
| PrL_Female_Water_4 | PrL | F | W | 4 | PrL_FW |
| PrL_Female_Water_3 | PrL | F | W | 3 | PrL_FW |
| VTA_Male_Fentanyl_5 | VTA | M | F | 5 | VTA_MF |
| VBT_Female_Water_2 | VBT | F | W | 2 | VBT_FW |
| S1_Male_Water_1 | S1 | M | W | 1 | S1_MW |
| VBT_Male_Fentanyl_1 | VBT | M | F | 1 | VBT_MF |
| NAc_Male_Water_1 | NAc | M | W | 1 | NAc_MW |
| NAc_Male_Water_5 | NAc | M | W | 5 | NAc_MW |
| VBT_Male_Fentanyl_4 | VBT | M | F | 4 | VBT_MF |
| S1_Male_Fentanyl_1 | S1 | M | F | 1 | S1_MF |
| VTA_Female_Fentanyl_3 | VTA | F | F | 3 | VTA_FF |
| NAc_Male_Water_4 | NAc | M | W | 4 | NAc_MW |
| PrL_Male_Water_1 | PrL | M | W | 1 | PrL_MW |
| NAc_Female_Fentanyl_1 | NAc | F | F | 1 | NAc_FF |
| S1_Female_Water_1 | S1 | F | W | 1 | S1_FW |
| VTA_Female_Water_3 | VTA | F | W | 3 | VTA_FW |
| VTA_Male_Water_4 | VTA | M | W | 4 | VTA_MW |
| NAc_Male_Fentanyl_2 | NAc | M | F | 2 | NAc_MF |
| VTA_Male_Fentanyl_3 | VTA | M | F | 3 | VTA_MF |
| NAc_Male_Fentanyl_3 | NAc | M | F | 3 | NAc_MF |
| VTA_Male_Fentanyl_4 | VTA | M | F | 4 | VTA_MF |
| S1_Male_Fentanyl_3 | S1 | M | F | 3 | S1_MF |
| VTA_Female_Fentanyl_4 | VTA | F | F | 4 | VTA_FF |
| PrL_Male_Water_5 | PrL | M | W | 5 | PrL_MW |
| VBT_Female_Fentanyl_4 | VBT | F | F | 4 | VBT_FF |
| S1_Female_Fentanyl_1 | S1 | F | F | 1 | S1_FF |
| NAc_Male_Water_2 | NAc | M | W | 2 | NAc_MW |
| PrL_Male_Water_4 | PrL | M | W | 4 | PrL_MW |
| NAc_Female_Water_2 | NAc | F | W | 2 | NAc_FW |
| PrL_Male_Fentanyl_5 | PrL | M | F | 5 | PrL_MF |
| VBT_Male_Fentanyl_3 | VBT | M | F | 3 | VBT_MF |
| S1_Male_Fentanyl_4 | S1 | M | F | 4 | S1_MF |
| NAc_Male_Water_3 | NAc | M | W | 3 | NAc_MW |
| VTA_Male_Water_5 | VTA | M | W | 5 | VTA_MW |
| NAc_Male_Fentanyl_5 | NAc | M | F | 5 | NAc_MF |
| VTA_Male_Water_3 | VTA | M | W | 3 | VTA_MW |
| S1_Male_Water_5 | S1 | M | W | 5 | S1_MW |
| VBT_Female_Fentanyl_2 | VBT | F | F | 2 | VBT_FF |
| VBT_Female_Fentanyl_1 | VBT | F | F | 1 | VBT_FF |
| PrL_Female_Water_1 | PrL | F | W | 1 | PrL_FW |
| VTA_Female_Fentanyl_2 | VTA | F | F | 2 | VTA_FF |
| VTA_Female_Water_5 | VTA | F | W | 5 | VTA_FW |
| NAc_Female_Fentanyl_5 | NAc | F | F | 5 | NAc_FF |
| VBT_Female_Water_3 | VBT | F | W | 3 | VBT_FW |
| VBT_Female_Fentanyl_3 | VBT | F | F | 3 | VBT_FF |
| PrL_Female_Fentanyl_4 | PrL | F | F | 4 | PrL_FF |
| VBT_Male_Water_4 | VBT | M | W | 4 | VBT_MW |
| NAc_Female_Water_4 | NAc | F | W | 4 | NAc_FW |
| VTA_Female_Fentanyl_5 | VTA | F | F | 5 | VTA_FF |
| PrL_Male_Water_3 | PrL | M | W | 3 | PrL_MW |
| PrL_Male_Fentanyl_1 | PrL | M | F | 1 | PrL_MF |
| PrL_Male_Water_2 | PrL | M | W | 2 | PrL_MW |

| sample.names.full | celltype | sex | trt | biorep | Group |
|---|---|---|---|---|---|
| S1_Female_Fentanyl_3 | S1 | F | F | 3 | S1_FF |
| S1_Female_Water_4 | S1 | F | W | 4 | S1_FW |
| VTA_Female_Water_1 | VTA | F | W | 1 | VTA_FW |
| VBT_Female_Water_4 | VBT | F | W | 4 | VBT_FW |

#### Create DGEList object

```
y <- DGEList(counts = counts, group = meta$Group)
keep <- rowSums(cpm(y) > 1) >= 2 & grepl("ENSMUSG",
    rownames(cpm(y)))

# In this case we remove genes that do NOT have at
# least 1 CPM in at least 2 samples
y = y[keep, , keep.lib.sizes = T]
y <- calcNormFactors(y)
logcpm = cpm(y, log = T, prior.count = 3)
```

##### MDS plot to detect sample outliers

```
col = vector(mode = "character", length = length(colnames(y$counts)))
pch = vector(mode = "character", length = length(colnames(y$counts)))

pch[grep("MF", colnames(y$counts))] = 16
pch[grep("MW", colnames(y$counts))] = 1
pch[grep("FF", colnames(y$counts))] = 15
pch[grep("FW", colnames(y$counts))] = 0

col[grep("PrL", colnames(y$counts))] = "springgreen3"
col[grep("NAc", colnames(y$counts))] = "orange"
col[grep("VTA", colnames(y$counts))] = "red2"
col[grep("VBT", colnames(y$counts))] = "blue"
col[grep("S1", colnames(y$counts))] = "hotpink"

# pdf(paste0(workdir,'/mds.pdf'),width=10,height=5)
par(mfrow = c(1, 2))
plotMDS(y, dim.plot = c(1, 2), cex = 1.2, top = 500,
    col = col, pch = as.numeric(pch))
legend("topright", col = c("springgreen3", "orange",
    "red2", "blue", "hotpink"), legend = c("Prl", "NAc",
    "VTA", "VBT", "S1"), pch = 20, cex = 1)
legend("bottomright", col = c("grey77"), legend = c("MW",
    "FW", "MF", "FF"), pch = c(1, 0, 16, 15), cex = 1)

plotMDS(y, dim.plot = c(2, 3), cex = 1.2, top = 500,
    col = col, pch = as.numeric(pch))
legend("topright", col = c("springgreen3", "orange",
    "red2", "blue", "hotpink"), legend = c("Prl", "NAc",
    "VTA", "VBT", "S1"), pch = 20, cex = 1)
```

```
legend("topleft", col = c("grey77"), legend = c("MW",
    "FW", "MF", "FF"), pch = c(1, 0, 16, 15), cex = 1)
```

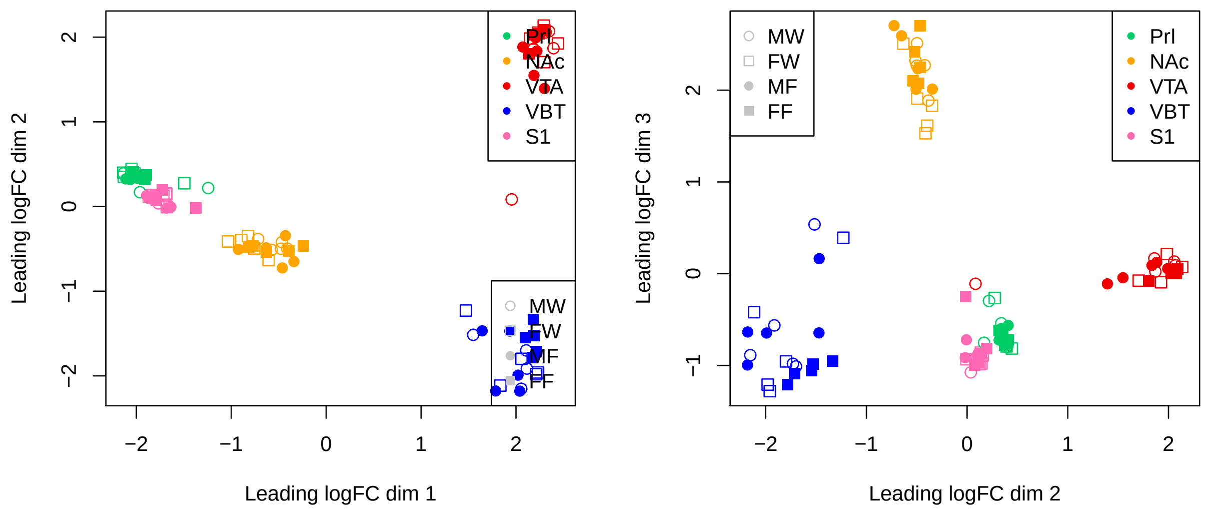


```
plotMDS(y[, meta$celltype == "PrL"], dim.plot = c(1,
    2), col = "springgreen3", cex = 1.2, pch = as.numeric(pch[meta$celltype ==
    "PrL"]), main = "Celltype: PrL")
legend("topleft", col = c("springgreen3"), legend = c("MW",
    "FW", "MF", "FF"), pch = c(1, 0, 16, 15), cex = 1)

plotMDS(y[, meta$celltype == "NAc"], dim.plot = c(1,
    2), col = "orange", cex = 1.2, pch = as.numeric(pch[meta$celltype ==
    "NAc"]), main = "Celltype: NAc")
legend("topleft", col = c("orange"), legend = c("MW",
    "FW", "MF", "FF"), pch = c(1, 0, 16, 15), cex = 1)
```

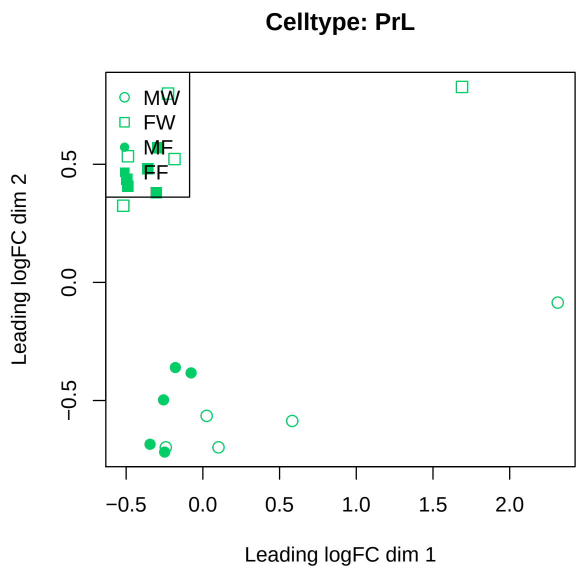


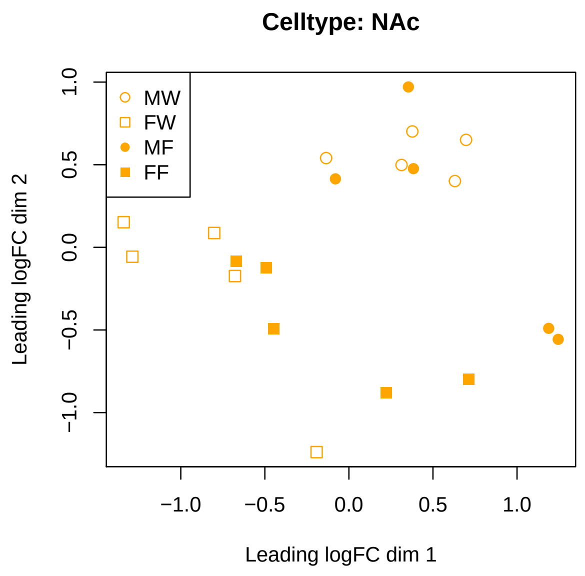


```
plotMDS(y[, meta$celltype == "VBT"], dim.plot = c(1,
    2), col = "blue", cex = 1.2, pch = as.numeric(pch[meta$celltype ==
    "VBT"]), main = "Celltype: VBT")
legend("bottomleft", col = c("blue"), legend = c("MW",
    "FW", "MF", "FF"), pch = c(1, 0, 16, 15), cex = 1)

plotMDS(y[, meta$celltype == "S1"], dim.plot = c(1,
    2), col = "hotpink", cex = 1.2, pch = as.numeric(pch[meta$celltype ==
    "S1"]), main = "Celltype: S1")
legend("topleft", col = c("hotpink"), legend = c("MW",
    "FW", "MF", "FF"), pch = c(1, 0, 16, 15), cex = 1)
```

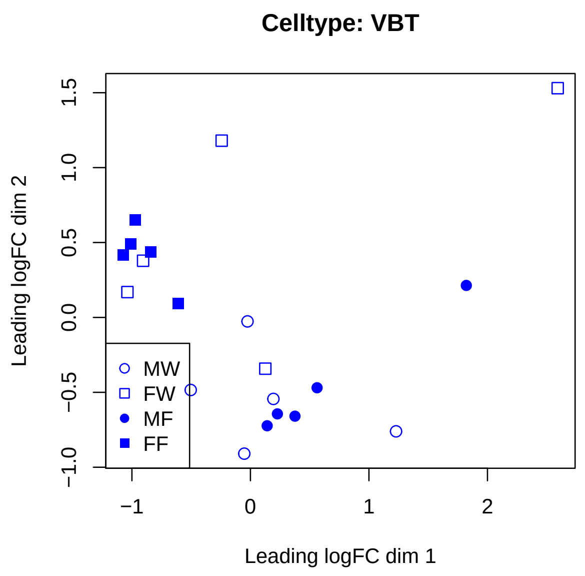


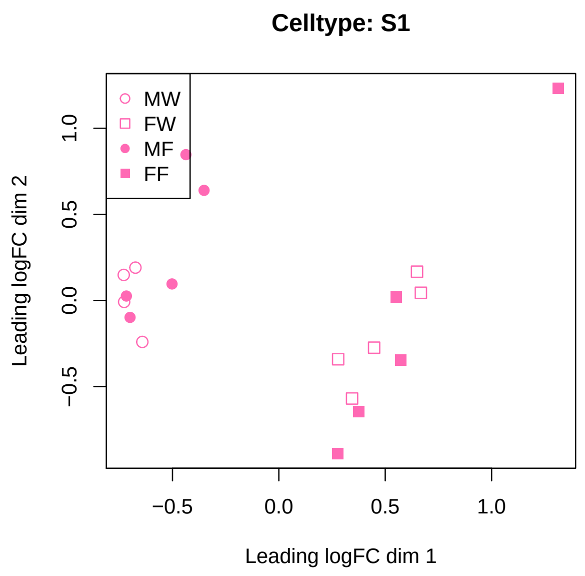

```r
plotMDS(y[, meta$celltype == "VTA"], dim.plot = c(1,
    2), col = "red2", cex = 1.2, pch = as.numeric(pch[meta$celltype ==
    "VTA"]), main = "Celltype: VTA")
legend("topleft", col = c("red2"), legend = c("MW",
    "FW", "MF", "FF"), pch = c(1, 0, 16, 15), cex = 1)
plotMDS(y[, meta$celltype == "VTA"], dim.plot = c(2,
    3), col = "red2", cex = 1.2, pch = as.numeric(pch[meta$celltype ==
    "VTA"]), main = "Celltype: VTA")
legend("topleft", col = c("red2"), legend = c("MW",
    "FW", "MF", "FF"), pch = c(1, 0, 16, 15), cex = 1)
```

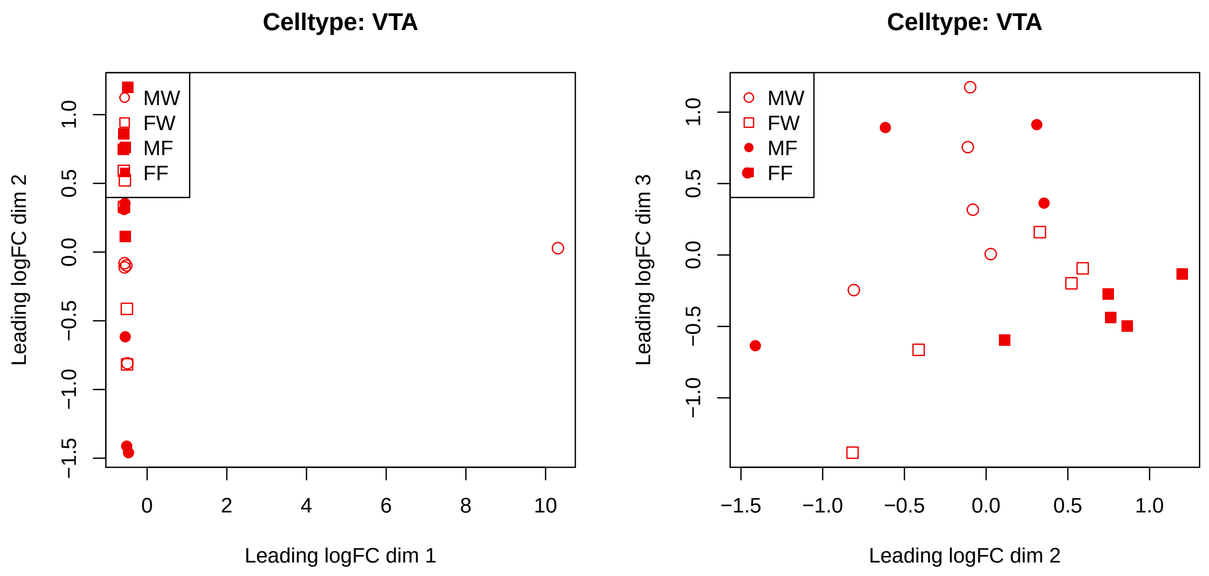


```r
plotMDS(y[, meta$celltype == "VTA"], dim.plot = c(1,
    2), col = "red2", cex = 1, main = "Celltype: VTA",
    xlim = c(-4, 14))
```

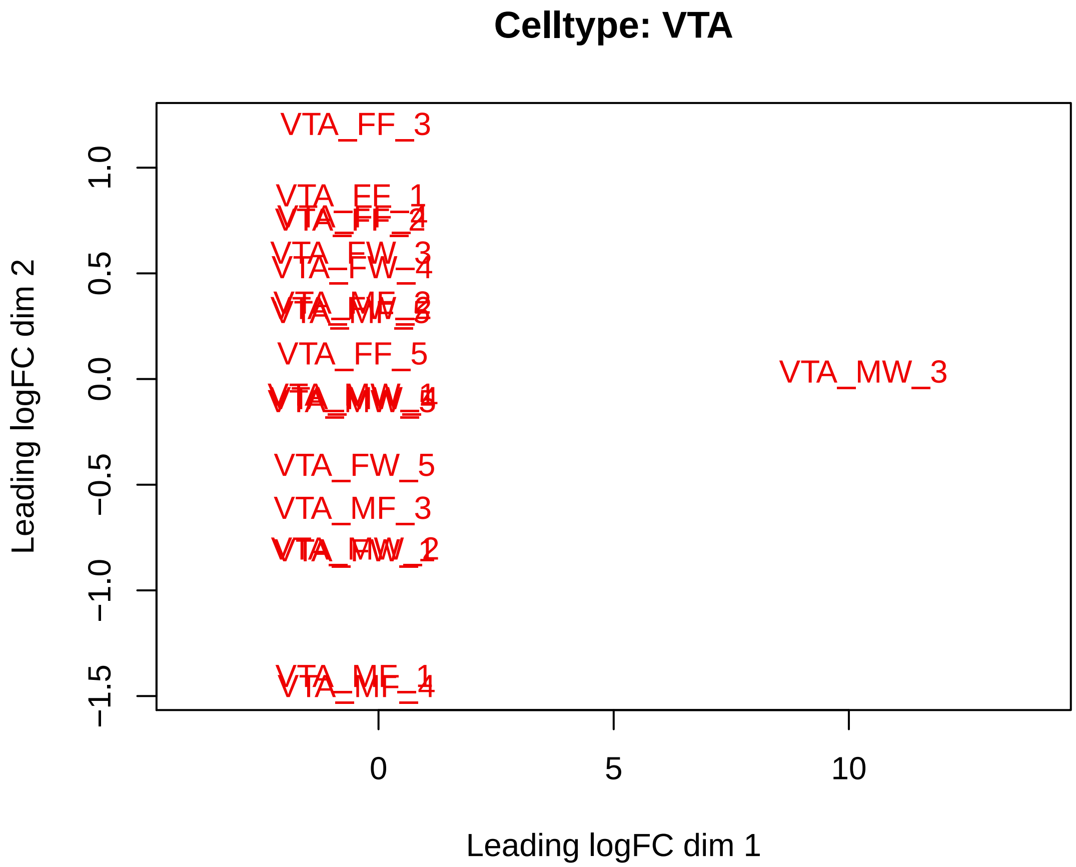


One VTA sample Male-water is outlier.

###### Hierarchical clustering to detect sample outliers

###### Clustering of samples using top 500 variable genes

```
var_genes <- apply(logcpm, 1, var)
select_var <- names(sort(var_genes, decreasing = TRUE))[1:500]
highly_variable_lcpm <- logcpm[select_var, ]
dim(highly_variable_lcpm)
```

```
## [1] 500  99
```

###### Plot the heatmap for the variable genes

```
mypalette <- brewer.pal(11, "RdYlBu")
morecols <- colorRampPalette(mypalette)
# Set up colour vector for celltype variable
col.cell <- c("orange", "springgreen3", "hotpink",
    "blue", "red2")[as.numeric(factor(meta$celltype))]
```

```r
heatmap.2(highly_variable_lcpm, col = rev(morecols(50)),
    trace = "none", main = "Top 500 most variable genes across samples",
    ColSideColors = col.cell, scale = "row")
```

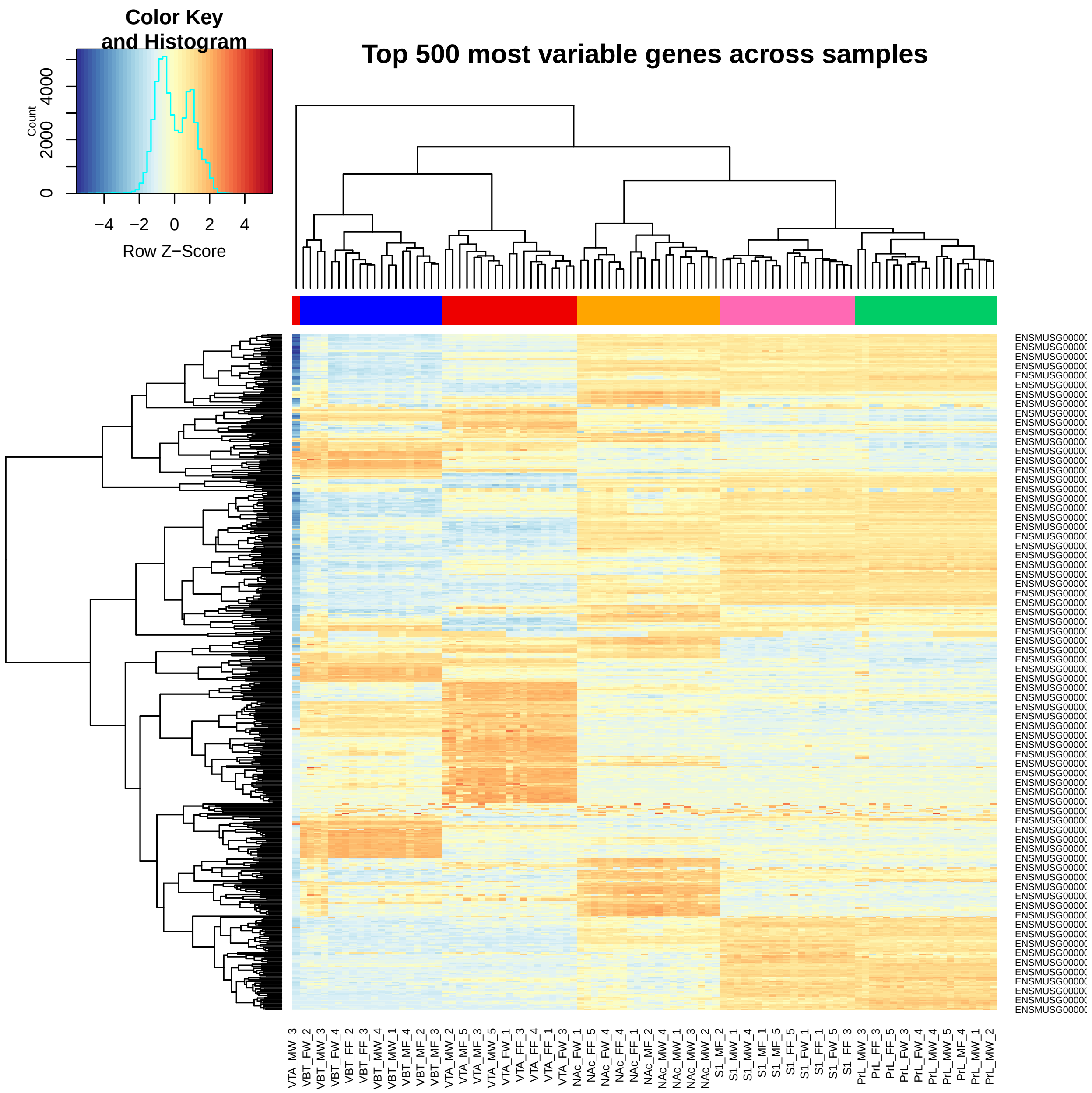


One sample is outlier: VTA_MW_3

#### Remove the outlier sample : VTA_MW_3

```r
countsall = counts
counts = counts[, -c(which(colnames(counts) == "VTA_MW_3"))]
meta.all = meta
meta = meta[-c(which(meta$sample.names.full == "VTA_Male_Water_3")),
```

```
    ]
y <- DGEList(counts = counts, group = meta$Group)
keep <- rowSums(cpm(y) > 1) >= 2 & grepl("ENSMUSG",
    rownames(cpm(y)))
y = y[keep, , keep.lib.sizes = T]
y <- calcNormFactors(y)
logcpm = cpm(y, log = T, prior.count = 3)
```

###### Clustering of samples using top 500 variable genes after removing the outlier sample

```
var_genes <- apply(logcpm, 1, var)
select_var <- names(sort(var_genes, decreasing = TRUE))[1:500]
highly_variable_lcpm <- logcpm[select_var, ]

mypalette <- brewer.pal(11, "RdYlBu")
morecols <- colorRampPalette(mypalette)
# Set up colour vector for celltype variable
col.cell <- c("orange", "springgreen3", "hotpink",
    "blue", "red2")[as.numeric(factor(meta$celltype))]

heatmap.2(highly_variable_lcpm, col = rev(morecols(50)),
    trace = "none", main = "Top 500 most variable genes across samples",
    ColSideColors = col.cell, scale = "row")
```

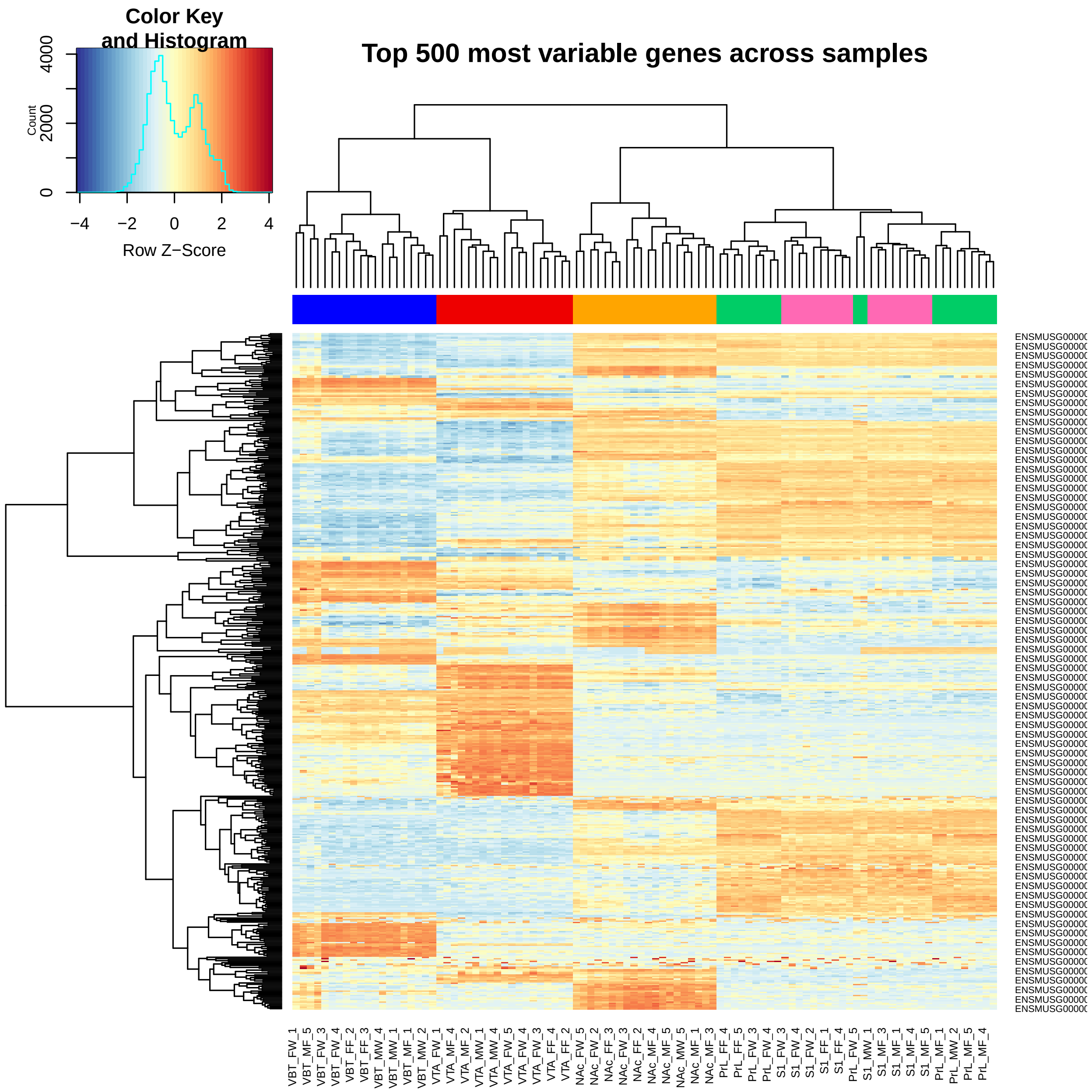


#### Differenial expression using lmFit

```
design = model.matrix(~0 + meta$Group)
colnames(design) = unlist(strsplit(colnames(design),
    "meta\\$Group"))[seq(2, 40, 2)]

fit1 = lmFit(logcpm, design = design)
fit1 <- eBayes(fit1)

contr = makeContrasts(NAcM = (NAc_MF - NAc_MW), NAcF = (NAc_FF -
    NAc_FW), NAc = (NAc_MF + NAc_FF)/2 - (NAc_MW +
    NAc_FW)/2, PrLM = (PrL_MF - PrL_MW), PrLF = (PrL_FF -
```

```
    PrL_FW), PrL = (PrL_MF + PrL_FF)/2 - (PrL_MW +
    PrL_FW)/2, S1M = (S1_MF - S1_MW), S1F = (S1_FF -
    S1_FW), S1 = (S1_MF + S1_FF)/2 - (S1_MW + S1_FW)/2,
    VBTM = (VBT_MF - VBT_MW), VBTF = (VBT_FF - VBT_FW),
    VBT = (VBT_MF + VBT_FF)/2 - (VBT_MW + VBT_FW)/2,
    VTAM = (VTA_MF - VTA_MW), VTAF = (VTA_FF - VTA_FW),
    VTA = (VTA_MF + VTA_FF)/2 - (VTA_MW + VTA_FW)/2,
    levels = design)

fit1a = contrasts.fit(fit1, contr)
fit1a = eBayes(fit1a, trend = T)

res1 = topTable(fit1a, coef = 1, number = Inf)
res1 = merge(res1, topTable(fit1a, coef = 2, number = Inf),
    by = 0)
res1 = merge(res1, topTable(fit1a, coef = 3, number = Inf),
    by.x = 1, by.y = 0)
res1 = merge(res1, topTable(fit1a, coef = 4, number = Inf),
    by.x = 1, by.y = 0)
res1 = merge(res1, topTable(fit1a, coef = 5, number = Inf),
    by.x = 1, by.y = 0)
res1 = merge(res1, topTable(fit1a, coef = 6, number = Inf),
    by.x = 1, by.y = 0)
res1 = merge(res1, topTable(fit1a, coef = 7, number = Inf),
    by.x = 1, by.y = 0)
res1 = merge(res1, topTable(fit1a, coef = 8, number = Inf),
    by.x = 1, by.y = 0)
res1 = merge(res1, topTable(fit1a, coef = 9, number = Inf),
    by.x = 1, by.y = 0)
res1 = merge(res1, topTable(fit1a, coef = 10, number = Inf),
    by.x = 1, by.y = 0)
res1 = merge(res1, topTable(fit1a, coef = 11, number = Inf),
    by.x = 1, by.y = 0)
res1 = merge(res1, topTable(fit1a, coef = 12, number = Inf),
    by.x = 1, by.y = 0)
res1 = merge(res1, topTable(fit1a, coef = 13, number = Inf),
    by.x = 1, by.y = 0)
res1 = merge(res1, topTable(fit1a, coef = 14, number = Inf),
    by.x = 1, by.y = 0)
res1 = merge(res1, topTable(fit1a, coef = 15, number = Inf),
    by.x = 1, by.y = 0)

n = ncol(contr)
limma.pvals = res1[, c(1:n * 6 - 1)]
limma.logFC = res1[, c(1:n * 6 - 4)]
limma.AveExpr = res1[, c(1:n * 6 - 3)]
colnames(limma.pvals) = colnames(limma.logFC) = colnames(limma.AveExpr) = colnames(contr)
rownames(limma.pvals) = rownames(limma.logFC) = rownames(limma.AveExpr) = res1[,
    1]

limma.fdr = apply(limma.pvals, 2, p.adjust)
colnames(limma.fdr) = paste("FDR", colnames(limma.fdr),
    sep = "_")
```

```r
colnames(limma.pvals) = paste("P.Value", colnames(limma.pvals),
    sep = "_")
colnames(limma.logFC) = paste("logFC", colnames(limma.logFC),
    sep = "_")
colnames(limma.AveExpr) = paste("AveExpr", colnames(limma.AveExpr),
    sep = "_")

summa.fit1a <- decideTests(fit1a, adjust.method = "none",
    p.value = 0.001)
```

Table 2: DE for pval=0.001

| | NAcM | NAcF | NAc | PrLM | PrLF | PrL | S1M | S1F | S1 | VBTM | VBTF | VBT | VTAM | VTAF | VTA |
|---|---|---|---|---|---|---|---|---|---|---|---|---|---|---|---|
| Down | 353 | 203 | 259 | 93 | 56 | 243 | 20 | 93 | 126 | 57 | 40 | 19 | 1465 | 105 | 1567 |
| NotSig | 17403 | 17955 | 17801 | 18106 | 18161 | 17875 | 18174 | 17852 | 17763 | 18124 | 18056 | 18176 | 15708 | 18066 | 15573 |
| Up | 478 | 76 | 174 | 35 | 17 | 116 | 40 | 289 | 345 | 53 | 138 | 39 | 1061 | 63 | 1094 |

Table 3: DE for fdr=0.05

| | NAcM | NAcF | NAc | PrLM | PrLF | PrL | S1M | S1F | S1 | VBTM | VBTF | VBT | VTAM | VTAF | VTA |
|---|---|---|---|---|---|---|---|---|---|---|---|---|---|---|---|
| Down | 617 | 124 | 315 | 2 | 3 | 241 | 0 | 98 | 160 | 21 | 18 | 2 | 2544 | 35 | 2634 |
| NotSig | 16818 | 18071 | 17722 | 18232 | 18231 | 17881 | 18234 | 17835 | 17665 | 18193 | 18137 | 18232 | 13691 | 18187 | 13553 |
| Up | 799 | 39 | 197 | 0 | 0 | 112 | 0 | 301 | 409 | 20 | 79 | 0 | 1999 | 12 | 2047 |

Table 4: DE for fdr=0.1

| | NAcM | NAcF | NAc | PrLM | PrLF | PrL | S1M | S1F | S1 | VBTM | VBTF | VBT | VTAM | VTAF | VTA |
|---|---|---|---|---|---|---|---|---|---|---|---|---|---|---|---|
| Down | 1001 | 343 | 565 | 51 | 5 | 492 | 0 | 228 | 349 | 37 | 40 | 6 | 3153 | 96 | 3216 |
| NotSig | 15967 | 17742 | 17258 | 18172 | 18229 | 17397 | 18234 | 17562 | 17249 | 18163 | 18060 | 18222 | 12532 | 18080 | 12393 |
| Up | 1266 | 149 | 411 | 11 | 0 | 345 | 0 | 444 | 636 | 34 | 134 | 6 | 2549 | 58 | 2625 |

##### Venn-diagram ( DEG lmFit : p-value<= 0.001 )

```r
celltype = unique(meta$celltype)
# print(a)
mfrow.old <- par()$mfrow
par(mfrow = c(2, 3))
for (i in celltype) {
    xx = summa.fit1a[, grep(i, colnames(summa.fit1a))]
    # a <- vennCounts(xx) vennDiagram(a)
    vennDiagram(xx, include = c("up", "down"), counts.col = c("red",
        "blue"), circle.col = c("red", "blue", "green3"))
}
par(mfrow = mfrow.old)
```

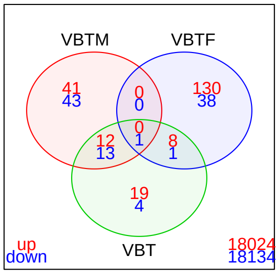


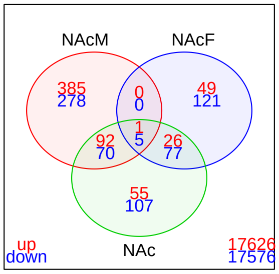


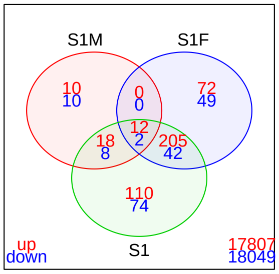


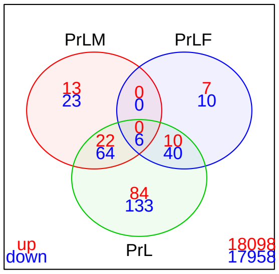


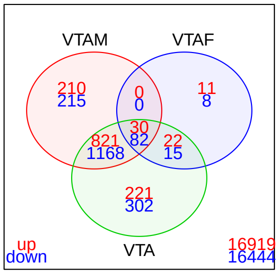


##### DEGs: overall for each celltype : lmFit

```
celltype = unique(meta$celltype)
sig.genes.pval = list()
pcut = 0.001
for (i in celltype) {
    sig.genes.pval[[i]] = rownames(limma.pvals)[rowSums(limma.pvals[,
        grep(i, colnames(limma.pvals))] <= pcut) >
        0]
}

sig.genes.fdr = list()
for (i in celltype) {
    sig.genes.fdr[[i]] = rownames(limma.fdr)[rowSums(limma.fdr[,
        grep(i, colnames(limma.fdr))] <= 0.1) > 0]
}

cbind(genepval = lapply(sig.genes.pval, length), genefdr = lapply(sig.genes.fdr,
    length))
```

```
##     genepval genefdr
## VBT 304      27
## NAc 1264     114
## S1  612      60
## PrL 412      23
## VTA 3105     981
```

```r
myCol <- brewer.pal(5, "Pastel1")
z1 = venn.diagram(sig.genes.pval, lwd = 2, fill = myCol,
    filename = NULL)
z2 = venn.diagram(sig.genes.fdr, lwd = 2, fill = myCol,
    filename = NULL)
```

**Overlap for DE pval<=0.001**

```r
grid.draw(z1)
```

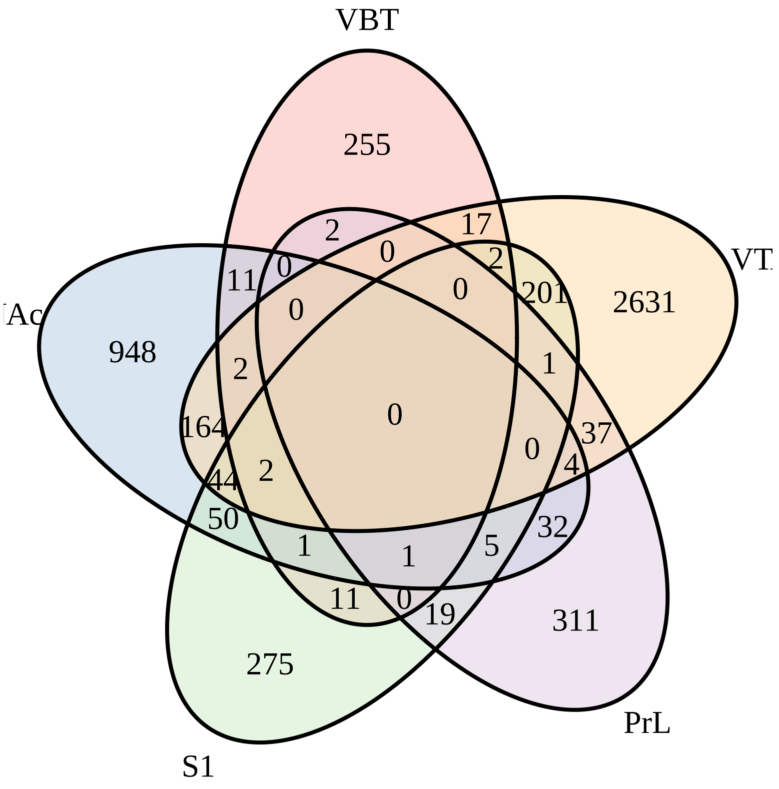


**Overlap for DE FDR<=0.001**

```r
grid.draw(z2)
```

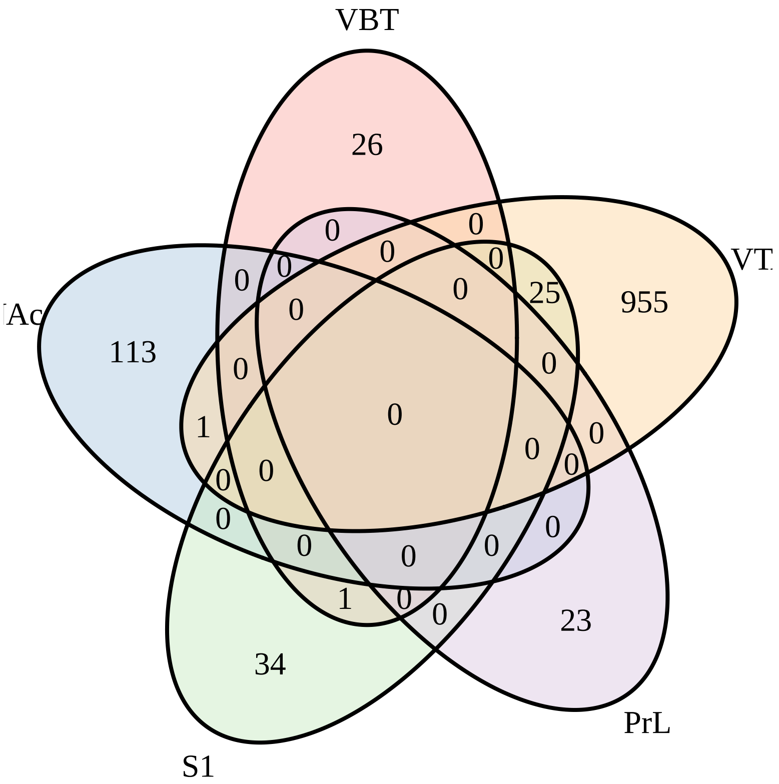


**Write the DEGs (lmFit )**

```
library(biomaRt)
mart = useMart("ensembl")
mart = useDataset("mmusculus_gene_ensembl", mart)
ann = getBM(mart = mart, attributes = c("ensembl_gene_id",
    "mgi_symbol", "gene_biotype", "description"), filters = "ensembl_gene_id",
    values = rownames(logcpm))
ann = ann[duplicated(ann$ensembl_gene_id) == F, ]

sigGenes.tab = lapply(1:length(celltype), function(i) {
    data.frame(ensembl_gene_id = sig.genes.pval[[celltype[i]]],
        limma.pvals[sig.genes.pval[[celltype[i]]],
            grep(celltype[i], colnames(limma.pvals))],
        limma.logFC[sig.genes.pval[[celltype[i]]],
            grep(celltype[i], colnames(limma.pvals))])
})
names(sigGenes.tab) = celltype

DEoutp = lapply(1:length(celltype), function(i) {
    merge(ann, sigGenes.tab[[celltype[i]]], by = 1)
})
names(DEoutp) = celltype

# Write all gene with DEGs pavls (lmFit )
sig.genes.pval = list()
pcut = 1
for (i in celltype) {
```

```r
    sig.genes.pval[[i]] = rownames(limma.pvals)[rowSums(limma.pvals[,
        grep(i, colnames(limma.pvals))] <= pcut) >
        0]
}
sigGenes.tab = lapply(1:length(celltype), function(i) {
    data.frame(ensembl_gene_id = sig.genes.pval[[celltype[i]]],
        limma.pvals[sig.genes.pval[[celltype[i]]],
            grep(celltype[i], colnames(limma.pvals))],
        limma.logFC[sig.genes.pval[[celltype[i]]],
            grep(celltype[i], colnames(limma.pvals))])
})
names(sigGenes.tab) = celltype

outp = lapply(1:length(celltype), function(i) {
    merge(ann, sigGenes.tab[[celltype[i]]], by = 1)
})
names(outp) = celltype

# save(counts,y,logcpm,meta,res1,limma.fdr,limma.pvals,limma.logFC,limma.AveExpr,outp,DEoutp,file=paste[illegible]
# save(countsall,meta.all,counts,y,logcpm,meta,res1,limma.fdr,limma.pvals,limma.logFC,limma.AveExpr,out[illegible]

for (i in 1:length(celltype)) {
    flname = paste0(workdir, "/DEgene_summary_", celltype[i],
        ".txt")
    write.table(outp[[celltype[i]]], file = flname,
        sep = "\t")
}
```

###### Heatmap of DEGs (lmFit ) : pval<=0.001

```r
celltype = unique(meta$celltype)
for (cell in celltype) {
    index = grep(cell, colnames(limma.pvals))
    flname = cell
    sig.genes = which(rowSums(limma.pvals[, index] <=
        0.001) > 0)
    z = (logcpm[sig.genes, grep(cell, colnames(logcpm))])
    z = cbind(rownames(z), z)
    outpx = merge(ann, z, by = 1)
    datExpr = outpx[, 5:ncol(outpx)]
    rownames(datExpr) = outpx[, "mgi_symbol"]
    x1 = rep(1, length(colnames(datExpr)))
    x2 = x1
    x1[grep("MW", colnames(datExpr))] = "darkolivegreen"
    x1[grep("FW", colnames(datExpr))] = "darkolivegreen3"
    x1[grep("MF", colnames(datExpr))] = "hotpink2"
    x1[grep("FF", colnames(datExpr))] = "lightpink"

    x = t(x1)
    rownames(x) = c("Expt")
    # if(cell=='D2'){pdf(file=paste0(workdir,'/DE_',flname,'_pval0.001_scaledRow.pdf'),height=20)
    # } else {
    # pdf(file=paste0(workdir,'/DE_',flname,'_pval0.001_scaledRow.pdf'))
```

```r
    heatmap.3(data.matrix(datExpr), scale = "row",
        margins = c(7, 12), ColSideColors = t(x), col = rev(brewer.pal(11,
            "RdBu")), ColSideColorsSize = 2, KeyValueName = "logcpm",
        main = cell)
    legend("topright", legend = c("MW", "FW", "MF",
        "FF"), fill = c("darkolivegreen", "darkolivegreen3",
        "hotpink2", "lightpink"), border = FALSE, bty = "n",
        cex = 0.8)
    # dev.off()
}
```

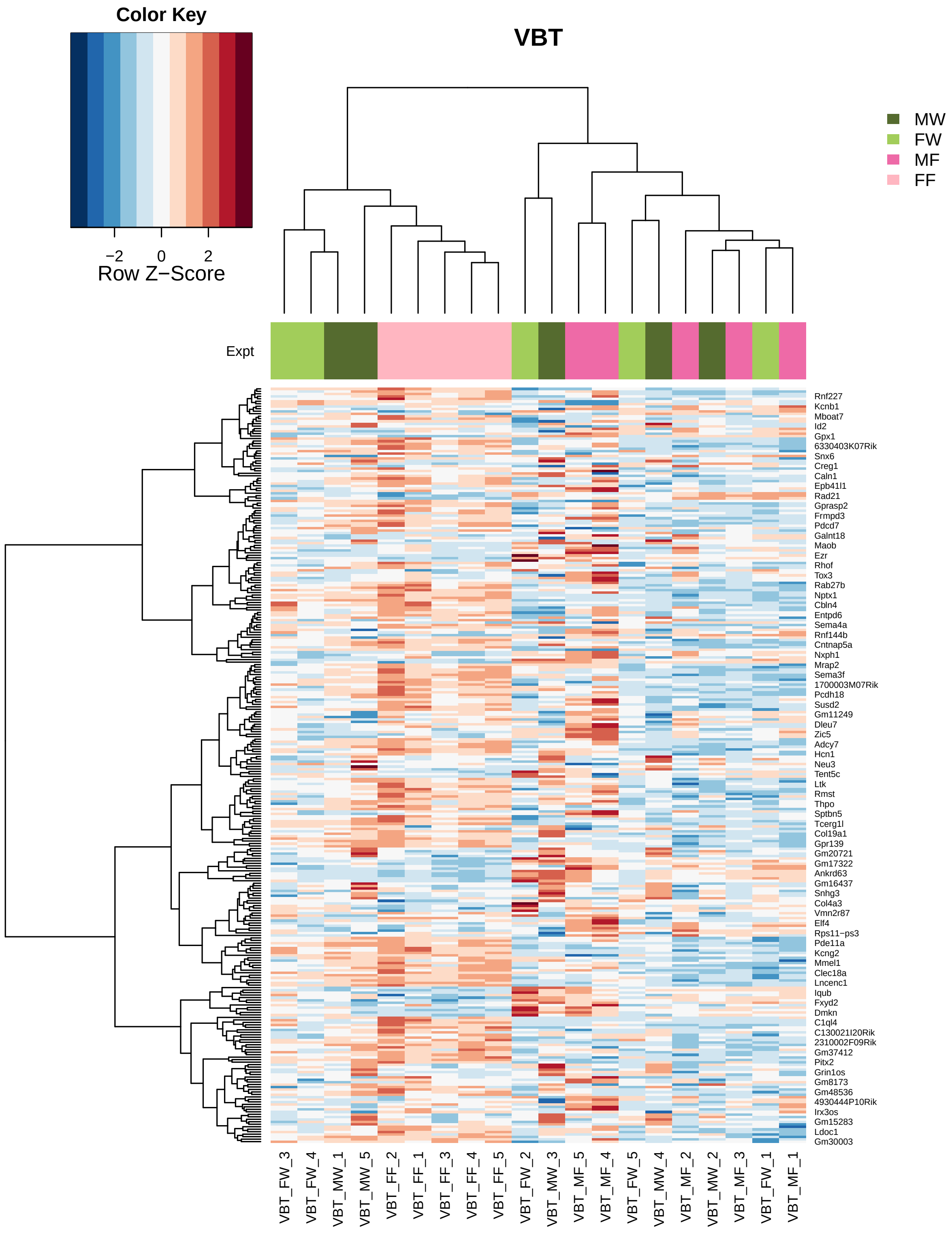
Color Key
−2
0
2
Row Z−Score
VBT
MW
FW
MF
FF
Expt
Rnf227
Kcnb1
Mboat7
Id2
Gpx1
6330403K07Rik
Snx6
Creg1
Caln1
Epb41l1
Rad21
Gprasp2
Frmpd3
Pdcd7
Galnt18
Maob
Ezr
Rhof
Tox3
Rab27b
Nptx1
Cbln4
Entpd6
Sema4a
Rnf144b
Cntnap5a
Nxph1
Mrap2
Sema3f
1700003M07Rik
Pcdh18
Susd2
Gm11249
Dleu7
Zic5
Adcy7
Hcn1
Neu3
Tent5c
Ltk
Rmst
Thpo
Sptbn5
Tcerg1l
Col19a1
Gpr139
Gm20721
Gm17322
Ankrd63
Gm16437
Snhg3
Col4a3
Vmn2r87
Elf4
Rps11−ps3
Pde11a
Kcng2
Mmel1
Clec18a
Lncenc1
Iqub
Fxyd2
Dmkn
C1ql4
C130021I20Rik
2310002F09Rik
Gm37412
Pitx2
Grin1os
Gm8173
Gm48536
4930444P10Rik
Irx3os
Gm15283
Ldoc1
Gm30003
VBT_FW_3
VBT_FW_4
VBT_MW_1
VBT_MW_5
VBT_FF_2
VBT_FF_1
VBT_FF_3
VBT_FF_4
VBT_FF_5
VBT_FW_2
VBT_MW_3
VBT_MF_5
VBT_MF_4
VBT_FW_5
VBT_MW_4
VBT_MF_2
VBT_MW_2
VBT_MF_3
VBT_FW_1
VBT_MF_1

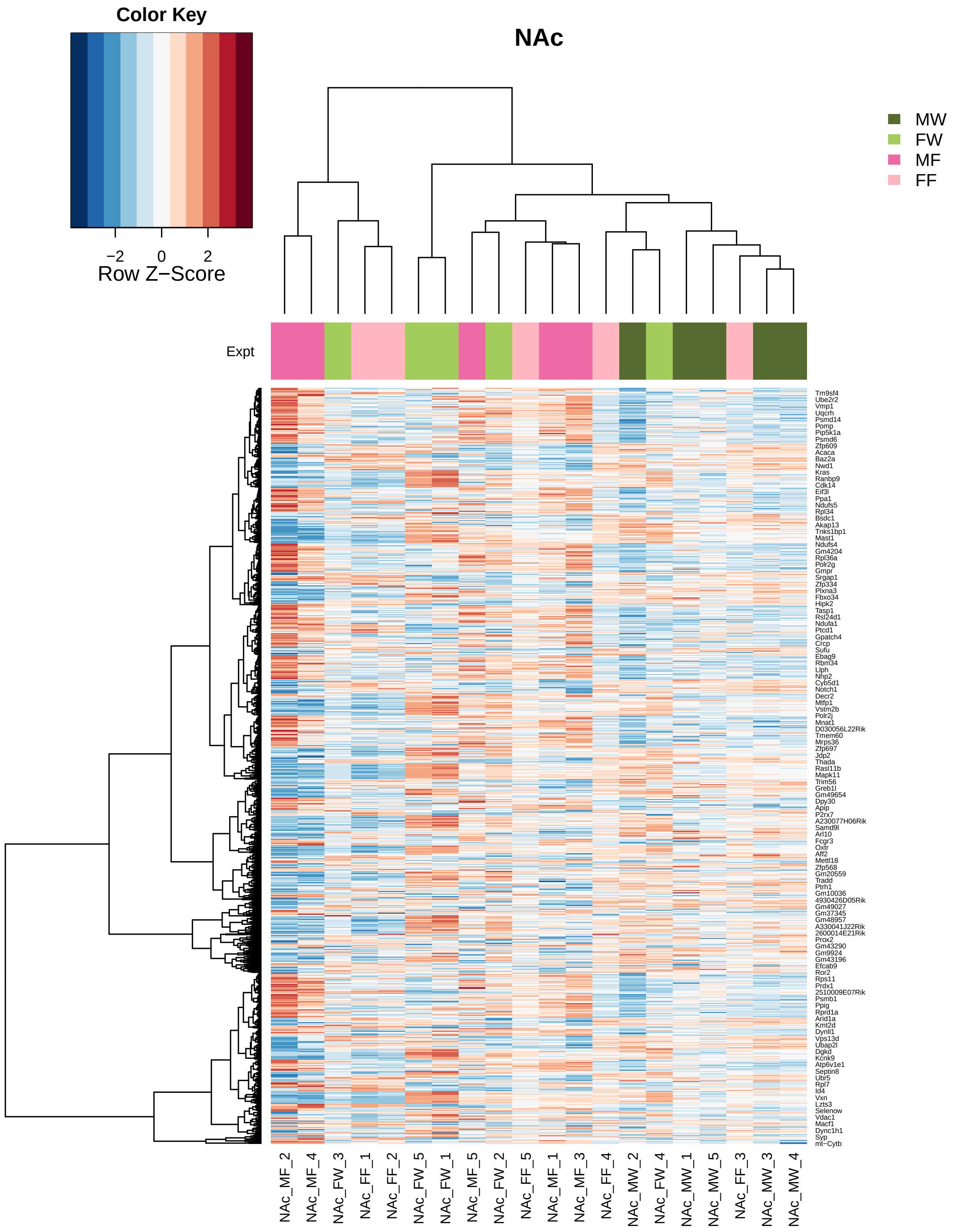

Color Key
−2
0
2
Row Z−Score
NAc
MW
FW
MF
FF
Expt
Tm9sf4
Ube2r2
Vmp1
Uqcrh
Psmd14
Pomp
Pip5k1a
Psmd6
Zfp609
Acaca
Baz2a
Nwd1
Kras
Ranbp9
Cdk14
Eif3l
Ppa1
Ndufs5
Rpl34
Bsdc1
Akap13
Tnks1bp1
Mast1
Ndufs4
Gm4204
Rpl36a
Polr2g
Gmpr
Srgap1
Zfp334
Plxna3
Fbxo34
Hipk2
Tasp1
Rsl24d1
Ndufa1
Ptcd1
Gpatch4
Crcp
Sufu
Ebag9
Rbm34
Llph
Nhp2
Cyb5d1
Notch1
Decr2
Mtfp1
Vstm2b
Polr2j
Mnat1
D030056L22Rik
Tmem60
Mrps36
Zfp697
Jdp2
Thada
Rasl11b
Mapk11
Trim56
Greb1l
Gm49654
Dpy30
Apip
P2rx7
A230077H06Rik
Samd9l
Arl10
Fcgr3
Oxtr
Aff2
Mettl18
Zfp568
Gm20559
Tradd
Ptrh1
Gm10036
4930426D05Rik
Gm49027
Gm37345
Gm48957
A330041J22Rik
2600014E21Rik
Prox2
Gm43290
Gm9924
Gm43196
Efcab9
Ror2
Rps11
Prdx1
2510009E07Rik
Psmb1
Ppig
Rprd1a
Arid1a
Kmt2d
Dynll1
Vps13d
Ubap2l
Dgkd
Kcnk9
Atp6v1e1
Septin8
Ubr5
Rpl7
Id4
Vxn
Lzts3
Selenow
Vdac1
Macf1
Dync1h1
Syp
mt−Cytb
NAc_MF_2
NAc_MF_4
NAc_FW_3
NAc_FF_1
NAc_FF_2
NAc_FW_5
NAc_FW_1
NAc_MF_5
NAc_FW_2
NAc_FF_5
NAc_MF_1
NAc_MF_3
NAc_FF_4
NAc_MW_2
NAc_FW_4
NAc_MW_1
NAc_MW_5
NAc_FF_3
NAc_MW_3
NAc_MW_4

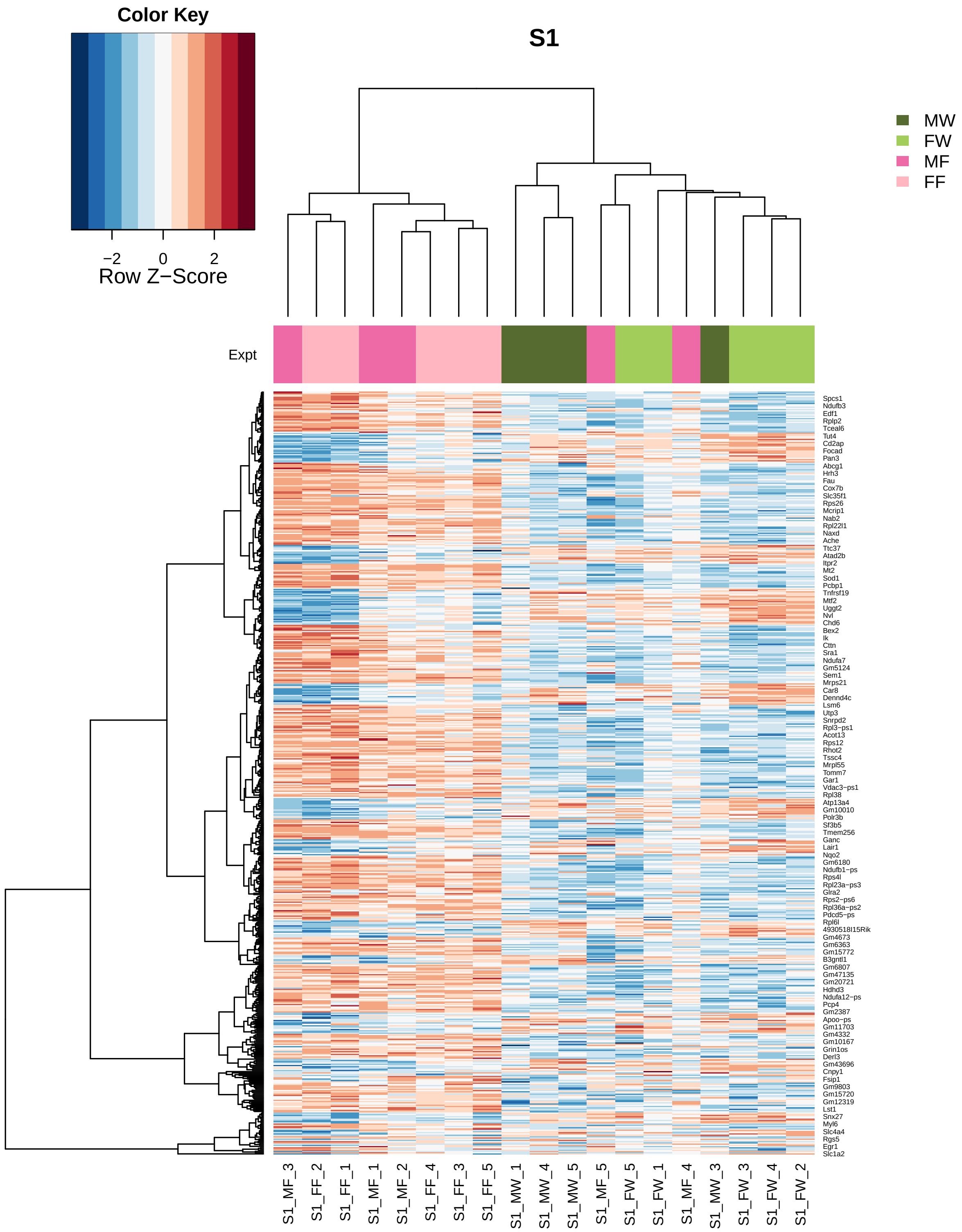

Color Key
−2
0
2
Row Z−Score
S1
MW
FW
MF
FF
Expt
Spcs1
Ndufb3
Edf1
Rplp2
Tceal6
Tut4
Cd2ap
Focad
Pan3
Abcg1
Hrh3
Fau
Cox7b
Slc35f1
Rps26
Mcrip1
Nab2
Rpl22l1
Naxd
Ache
Ttc37
Atad2b
Itpr2
Mt2
Sod1
Pcbp1
Tnfrsf19
Mtf2
Uggt2
Nvl
Chd6
Bex2
Ik
Cttn
Sra1
Ndufa7
Gm5124
Sem1
Mrps21
Car8
Dennd4c
Lsm6
Utp3
Snrpd2
Rpl3−ps1
Acot13
Rps12
Rhot2
Tssc4
Mrpl55
Tomm7
Gar1
Vdac3−ps1
Rpl38
Atp13a4
Gm10010
Polr3b
Sf3b5
Tmem256
Ganc
Lair1
Nqo2
Gm6180
Ndufb1−ps
Rps4l
Rpl23a−ps3
Glra2
Rps2−ps6
Rpl36a−ps2
Pdcd5−ps
Rpl6l
4930518I15Rik
Gm4673
Gm6363
Gm15772
B3gntl1
Gm6807
Gm47135
Gm20721
Hdhd3
Ndufa12−ps
Pcp4
Gm2387
Apoo−ps
Gm11703
Gm4332
Gm10167
Grin1os
Derl3
Gm43696
Cnpy1
Fsip1
Gm9803
Gm15720
Gm12319
Lst1
Snx27
Myl6
Slc4a4
Rgs5
Egr1
Slc1a2
S1_MF_3
S1_FF_2
S1_FF_1
S1_MF_1
S1_MF_2
S1_FF_4
S1_FF_3
S1_FF_5
S1_MW_1
S1_MW_4
S1_MW_5
S1_MF_5
S1_FW_5
S1_FW_1
S1_MF_4
S1_MW_3
S1_FW_3
S1_FW_4
S1_FW_2

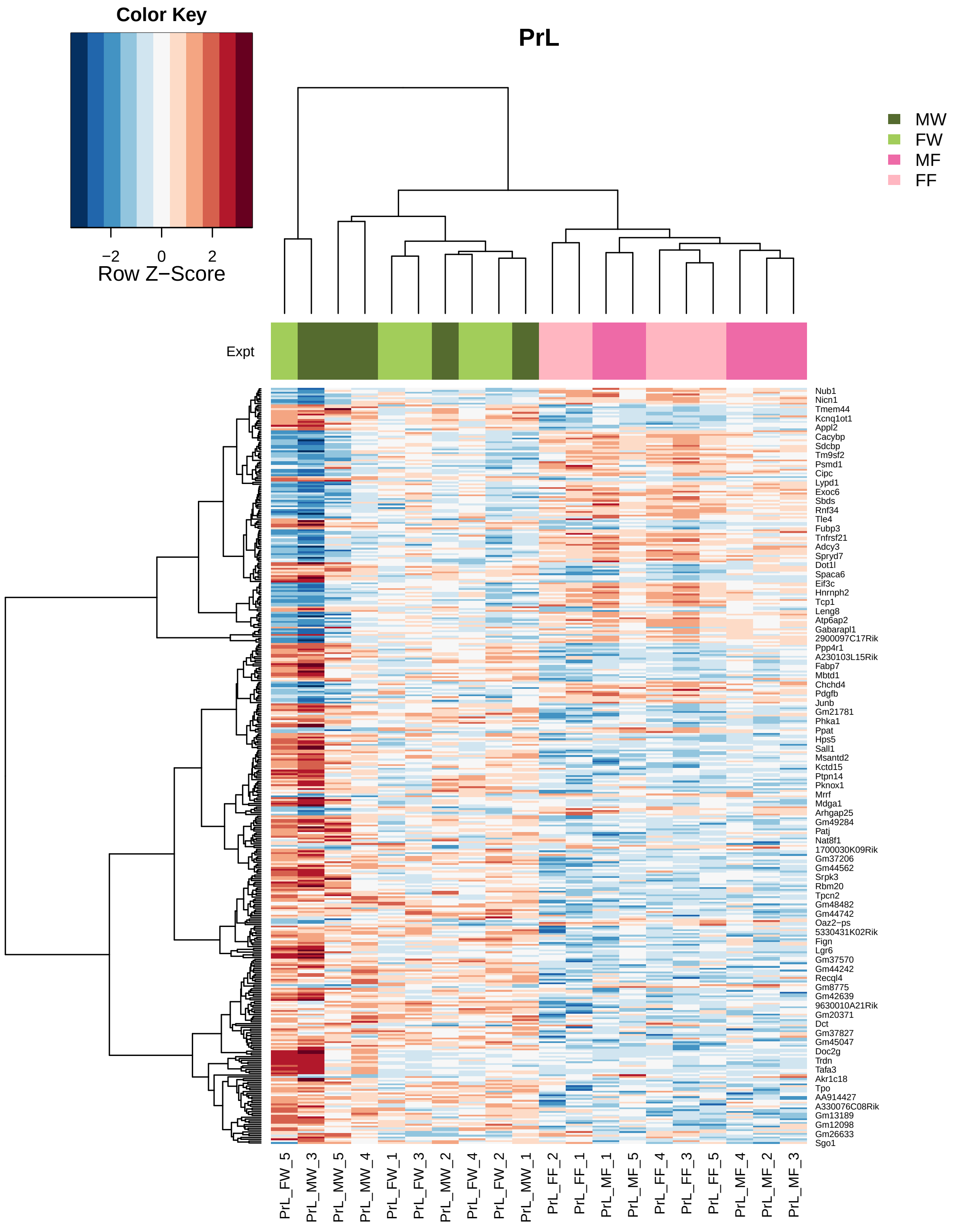
Color Key
−2
0
2
Row Z−Score
PrL
MW
FW
MF
FF
Expt
Nub1
Nicn1
Tmem44
Kcnq1ot1
Appl2
Cacybp
Sdcbp
Tm9sf2
Psmd1
Cipc
Lypd1
Exoc6
Sbds
Rnf34
Tle4
Fubp3
Tnfrsf21
Adcy3
Spryd7
Dot1l
Spaca6
Eif3c
Hnrnph2
Tcp1
Leng8
Atp6ap2
Gabarapl1
2900097C17Rik
Ppp4r1
A230103L15Rik
Fabp7
Mbtd1
Chchd4
Pdgfb
Junb
Gm21781
Phka1
Ppat
Hps5
Sall1
Msantd2
Kctd15
Ptpn14
Pknox1
Mrrf
Mdga1
Arhgap25
Gm49284
Patj
Nat8f1
1700030K09Rik
Gm37206
Gm44562
Srpk3
Rbm20
Tpcn2
Gm48482
Gm44742
Oaz2−ps
5330431K02Rik
Fign
Lgr6
Gm37570
Gm44242
Recql4
Gm8775
Gm42639
9630010A21Rik
Gm20371
Dct
Gm37827
Gm45047
Doc2g
Trdn
Tafa3
Akr1c18
Tpo
AA914427
A330076C08Rik
Gm13189
Gm12098
Gm26633
Sgo1
PrL_FW_5
PrL_MW_3
PrL_MW_5
PrL_MW_4
PrL_FW_1
PrL_FW_3
PrL_MW_2
PrL_FW_4
PrL_FW_2
PrL_MW_1
PrL_FF_2
PrL_FF_1
PrL_MF_1
PrL_MF_5
PrL_FF_4
PrL_FF_3
PrL_FF_5
PrL_MF_4
PrL_MF_2
PrL_MF_3

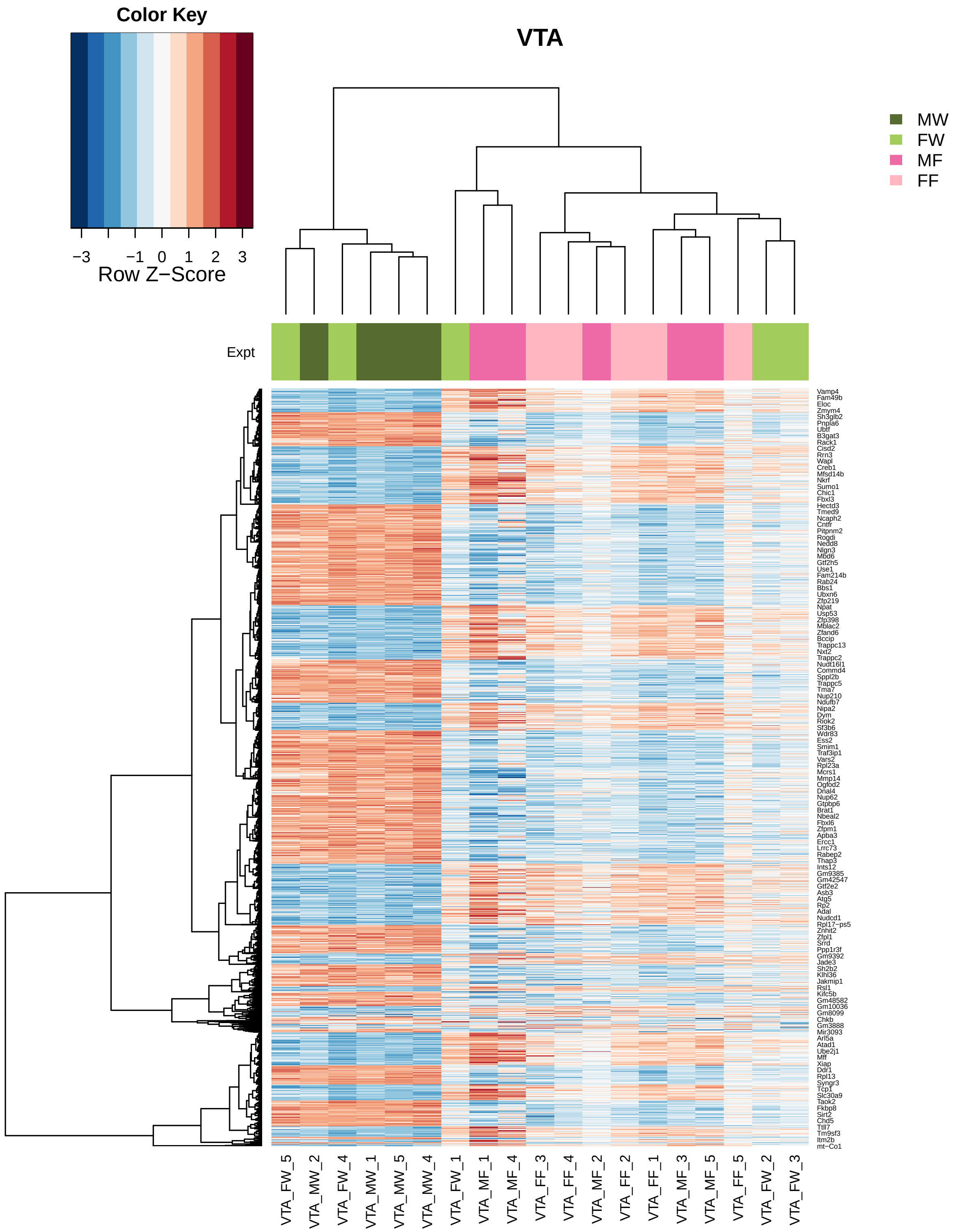
Color Key
−3
−1
0
1
2
3
Row Z−Score
VTA
MW
FW
MF
FF
Expt
VTA_FW_5
VTA_MW_2
VTA_FW_4
VTA_MW_1
VTA_MW_5
VTA_MW_4
VTA_FW_1
VTA_MF_1
VTA_MF_4
VTA_FF_3
VTA_FF_4
VTA_MF_2
VTA_FF_2
VTA_FF_1
VTA_MF_3
VTA_MF_5
VTA_FF_5
VTA_FW_2
VTA_FW_3

#### Differenial expression using limma-Voom

```
design = model.matrix(~0 + meta$Group)
colnames(design) = unlist(strsplit(colnames(design),
    "meta\\$Group"))[seq(2, 40, 2)]
contr = makeContrasts(NAcM = (NAc_MF - NAc_MW), NAcF = (NAc_FF -
    NAc_FW), NAc = (NAc_MF + NAc_FF)/2 - (NAc_MW +
    NAc_FW)/2, PrLM = (PrL_MF - PrL_MW), PrLF = (PrL_FF -
    PrL_FW), PrL = (PrL_MF + PrL_FF)/2 - (PrL_MW +
    PrL_FW)/2, S1M = (S1_MF - S1_MW), S1F = (S1_FF -
    S1_FW), S1 = (S1_MF + S1_FF)/2 - (S1_MW + S1_FW)/2,
    VBTM = (VBT_MF - VBT_MW), VBTF = (VBT_FF - VBT_FW),
    VBT = (VBT_MF + VBT_FF)/2 - (VBT_MW + VBT_FW)/2,
    VTAM = (VTA_MF - VTA_MW), VTAF = (VTA_FF - VTA_FW),
    VTA = (VTA_MF + VTA_FF)/2 - (VTA_MW + VTA_FW)/2,
    levels = design)
v <- voom(y, design, plot = TRUE)
```

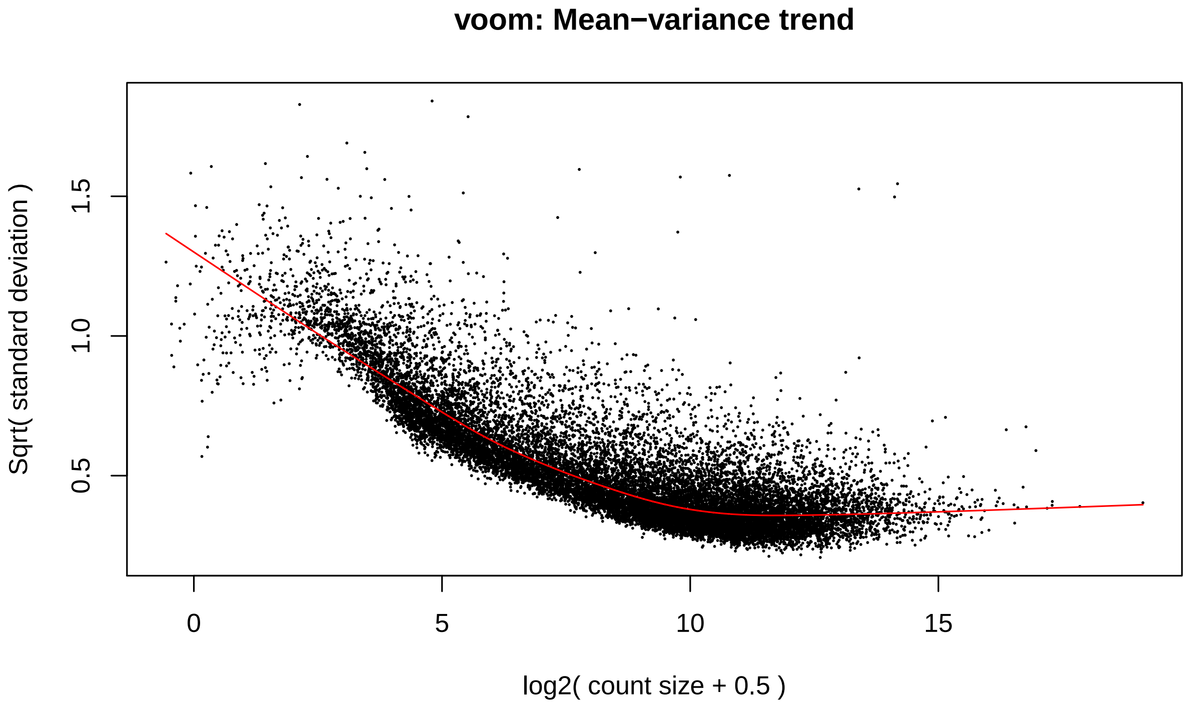


```
par(mfrow = c(2, 1))
boxplot(logcpm, xlab = "", ylab = "Log2 counts per million",
    las = 2, main = "Unnormalised logCPM")
abline(h = median(logcpm), col = "blue")
boxplot(v$E, xlab = "", ylab = "Log2 counts per million",
    las = 2, main = "Voom transformed logCPM")
abline(h = median(v$E), col = "blue")
```

**Unnormalised logCPM**

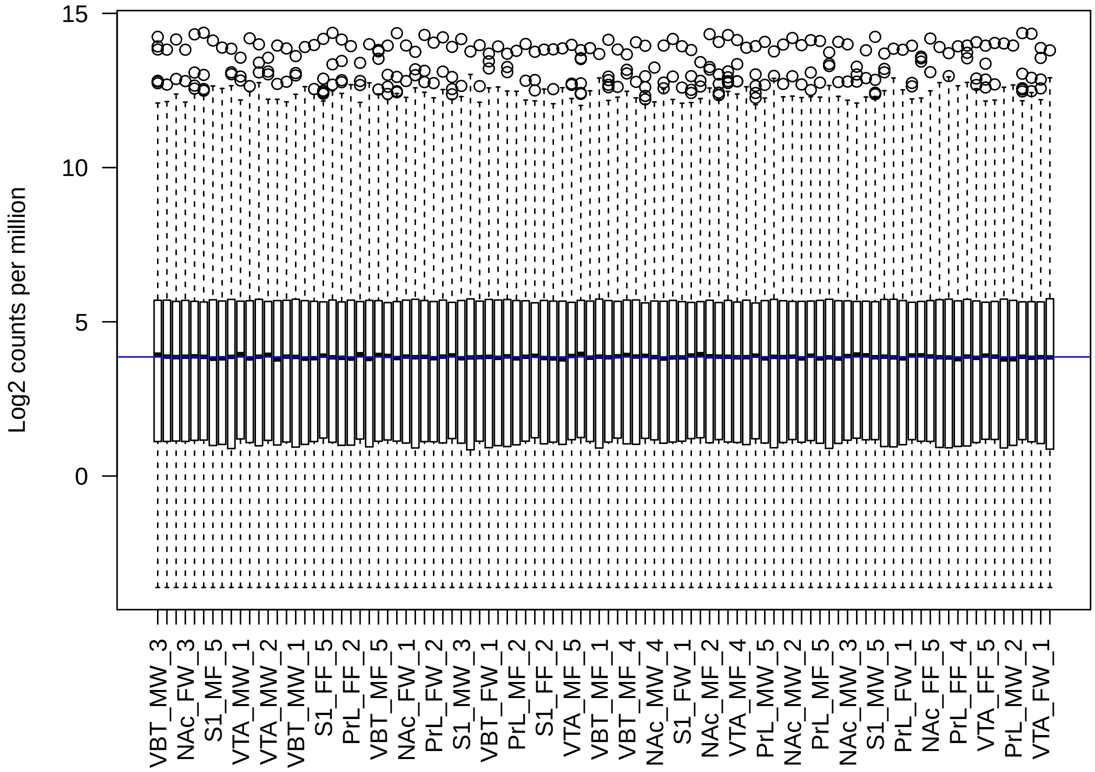


**Voom transformed logCPM**

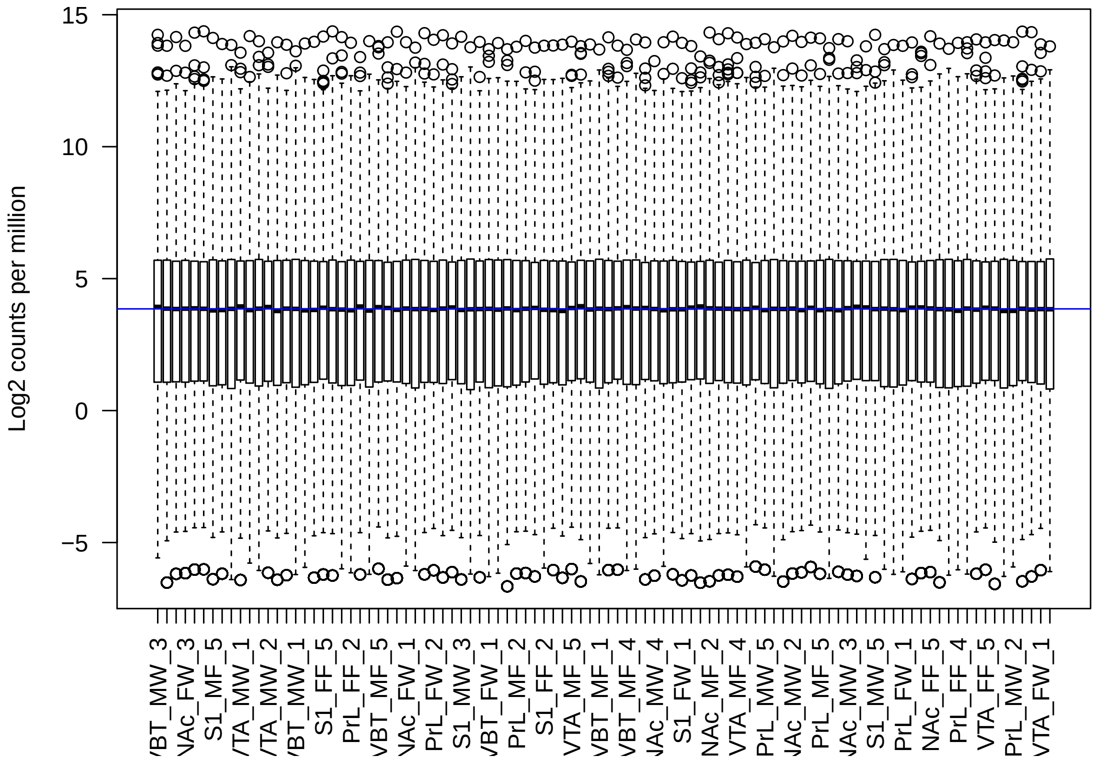

```
# Model fit: Voom
fit <- lmFit(v)
fit.cont = contrasts.fit(fit, contr)
fit.cont <- eBayes(fit.cont)
summa.fit <- decideTests(fit.cont, adjust.method = "none",
    p.value = 0.001)
```

Table 5: DE for pval=0.001

| | NAcM | NAcF | NAc | PrLM | PrLF | PrL | S1M | S1F | S1 | VBTM | VBTF | VBT | VTAM | VTAF | VTA |
|---|---|---|---|---|---|---|---|---|---|---|---|---|---|---|---|
| Down | 355 | 200 | 263 | 69 | 32 | 202 | 29 | 108 | 175 | 59 | 23 | 18 | 1567 | 94 | 1675 |
| NotSig | 17331 | 17915 | 17718 | 18119 | 18185 | 17892 | 18178 | 17867 | 17755 | 18132 | 18114 | 18194 | 15594 | 18070 | 15441 |
| Up | 548 | 119 | 253 | 46 | 17 | 140 | 27 | 259 | 304 | 43 | 97 | 22 | 1073 | 70 | 1118 |

```
for (i in 1:ncol(contr)) {
    if (i == 1) {
        res1 = topTable(fit.cont, coef = i, number = Inf)
    }
    if (i == 2) {
        res1 = merge(res1, topTable(fit.cont, coef = i,
            number = Inf), by = 0)
    }
    if (i > 2) {
        res1 = merge(res1, topTable(fit.cont, coef = i,
            number = Inf), by.x = 1, by.y = 0)
    }
}

n = ncol(contr)
voom.pvals = res1[, c(1:n * 6 - 1)]
voom.logFC = res1[, c(1:n * 6 - 4)]
voom.AveExpr = res1[, c(1:n * 6 - 3)]
colnames(voom.pvals) = colnames(voom.logFC) = colnames(voom.AveExpr) = colnames(contr)
rownames(voom.pvals) = rownames(voom.logFC) = rownames(voom.AveExpr) = res1[,
    1]

voom.fdr = apply(voom.pvals, 2, p.adjust)
colnames(voom.fdr) = paste("FDR", colnames(voom.fdr),
    sep = "_")
colnames(voom.pvals) = paste("P.Value", colnames(voom.pvals),
    sep = "_")
colnames(voom.logFC) = paste("logFC", colnames(voom.logFC),
    sep = "_")
colnames(voom.AveExpr) = paste("AveExpr", colnames(voom.AveExpr),
    sep = "_")

# save(counts,y,logcpm,meta,res1,limma.fdr,limma.pvals,limma.logFC,limma.AveExpr,voom.fdr,voom.pvals,
# voom.logFC, voom.AveExpr,
# outp,DEoutp,file=paste0(workdir,'/dev_fentanyl_DE.Rdata'))
# save(countsall,meta.all,counts,y,v,logcpm,meta,res1,limma.fdr,limma.pvals,limma.logFC,limma.AveExpr,v
# voom.logFC, voom.AveExpr,
```

```r
# outp,DEoutp,file=paste0(workdir,'/dev_fentanyl_DE_rmoutlier.Rdata'))
```

##### Venn-diagram DEG Voom

```r
celltype = unique(meta$celltype)
# print(a)
mfrow.old <- par()$mfrow
par(mfrow = c(2, 3))
for (i in celltype) {
    xx = summa.fit[, grep(i, colnames(summa.fit))]
    # a <- vennCounts(xx) vennDiagram(a)
    vennDiagram(xx, include = c("up", "down"), counts.col = c("red",
        "blue"), circle.col = c("red", "blue", "green3"))
}
par(mfrow = mfrow.old)
```

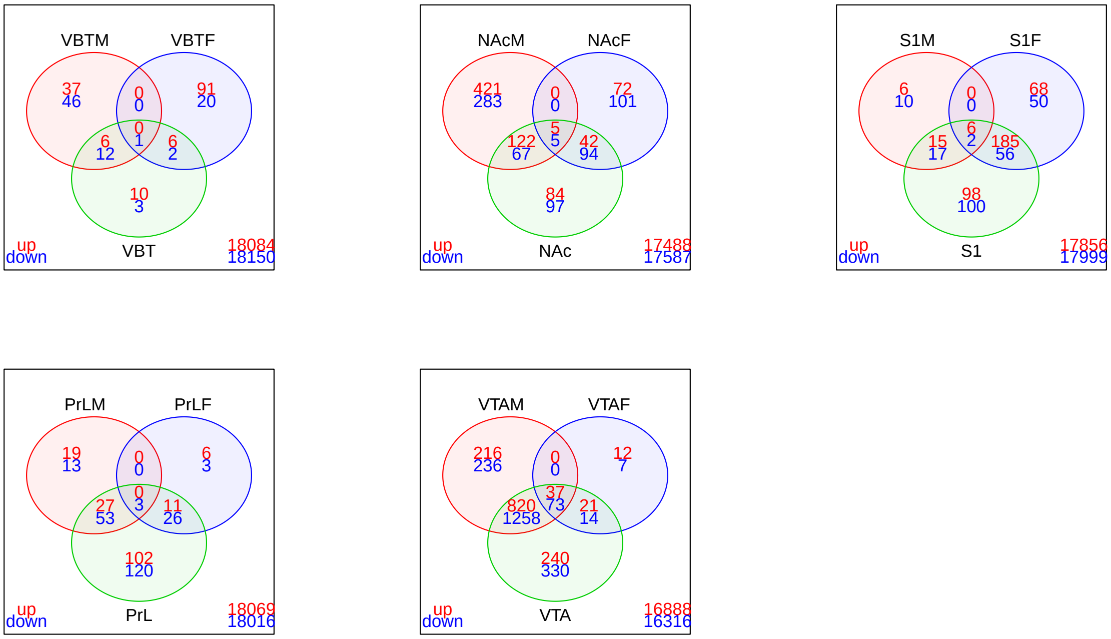


#### Compare DEG from Voom and limma

#### Gene Ontology enrichment of DEG : geneSetTest
