## Supplementary data 4 for "Transcriptomic profiling of reward and sensory brain areas in perinatal fentanyl exposed juvenile mice"

### Report WGCNA analysis for Developmental Fentanyl Exposure

```r
library(knitr)
opts_chunk$set(tidy.opts=list(width.cutoff=50),tidy=TRUE)
```

```r
workdir = "/local/projects/idea/mbasu/XLOBO/dev_fentanyl/plot1"
options(stringsAsFactors = F)
library(data.table)
library(WGCNA)
library(edgeR)
library(knitr)
library(dplyr)
library(kableExtra)
library(png)
library(VennDiagram)
library(gridExtra)
library(gplots)
library("devtools")
library("xlsx")
library(r2excel)
allowWGCNAThreads()
```

```
## Allowing multi-threading with up to 4 threads.
```

```r
options(knitr.table.format = "markdown")
```

#### Load DE data

```r
load(paste0(workdir, "/dev_fentanyl_DE_rmoutlier.Rdata"))
```

#### Load WGCNA data

*Evaluation code is below*

```r
load(paste0(workdir, "/powertab.Rdata"))
load(paste0(workdir, "/net_PrL.Rdata"))
load(paste0(workdir, "/net_NAc.Rdata"))
load(paste0(workdir, "/net_VBT.Rdata"))
load(paste0(workdir, "/net_S1.Rdata"))
load(paste0(workdir, "/net_VTA.Rdata"))
```

### 1. Building WGCNA network using all the genes, for each brain region

```
datExpr = t(logcpm)
celltype = unique(meta$celltype)
nSets = length(celltype)
shortLabels = setLabels = celltype
multiExpr = vector(mode = "list", length = nSets)
for (i in 1:length(celltype)) {
    multiExpr[[i]] = list(data = datExpr[grep(celltype[i],
        rownames(datExpr)), ])
}
exprSize = checkSets(multiExpr)
nGenes = exprSize$nGenes
nSamples = exprSize$nSamples
```

##### Power calculation

```
powers = c(c(1:10), seq(from = 12, to = 20, by = 2))
powerTables = vector(mode = "list", length = nSets)
for (set in 1:nSets) {
    powerTables[[set]] = list(data = pickSoftThreshold(multiExpr[[set]]$data,
        powerVector = powers, verbose = 2)[[2]])
}
# save(powerTables,file=paste0(workdir,'/powertab.Rdata'))
```

##### Plot:Determine Power

```
powers = c(c(1:10), seq(from = 12, to = 20, by = 2))
colors = c("red", "blue", "green", "orange", "darkorchid1")
# Will plot these columns of the returned scale
# free analysis tables
plotCols = c(2, 5, 6, 7)
colNames = c("Scale Free Topology Model Fit", "Mean connectivity",
    "Median connectivity", "Max connectivity")
# Get the minima and maxima of the plotted points
ylim = matrix(NA, nrow = 2, ncol = 4)
for (set in 1:nSets) {
    for (col in 1:length(plotCols)) {
        ylim[1, col] = min(ylim[1, col], powerTables[[set]]$data[,
            plotCols[col]], na.rm = TRUE)
        ylim[2, col] = max(ylim[2, col], powerTables[[set]]$data[,
            plotCols[col]], na.rm = TRUE)
    }
}

# sizeGrWindow(8, 6) pdf(file =
# paste0(workdir,'/power_celltype.pdf'), wi = 8, he
# = 6)
par(mfcol = c(2, 2))
```

```
par(mar = c(4.2, 4.2, 2.2, 0.5))
cex1 = 0.7
for (col in 1:length(plotCols)) for (set in 1:nSets) {
    if (set == 1) {
        plot(powerTables[[set]]$data[, 1], -sign(powerTables[[set]]$data[,
            3]) * powerTables[[set]]$data[, 2], xlab = "Soft Threshold (power)",
            ylab = colNames[col], type = "n", ylim = ylim[,
                col], main = colNames[col])
        addGrid()
    }
    if (col == 1) {
        text(powerTables[[set]]$data[, 1], -sign(powerTables[[set]]$data[,
            3]) * powerTables[[set]]$data[, 2], labels = powers,
            cex = cex1, col = colors[set])
    } else text(powerTables[[set]]$data[, 1], powerTables[[set]]$data[,
        plotCols[col]], labels = powers, cex = cex1,
        col = colors[set])
    if (col == 1) {
        legend("bottomright", legend = setLabels, col = colors,
            pch = 20)
    } else legend("topright", legend = setLabels, col = colors,
        pch = 20)
}
```

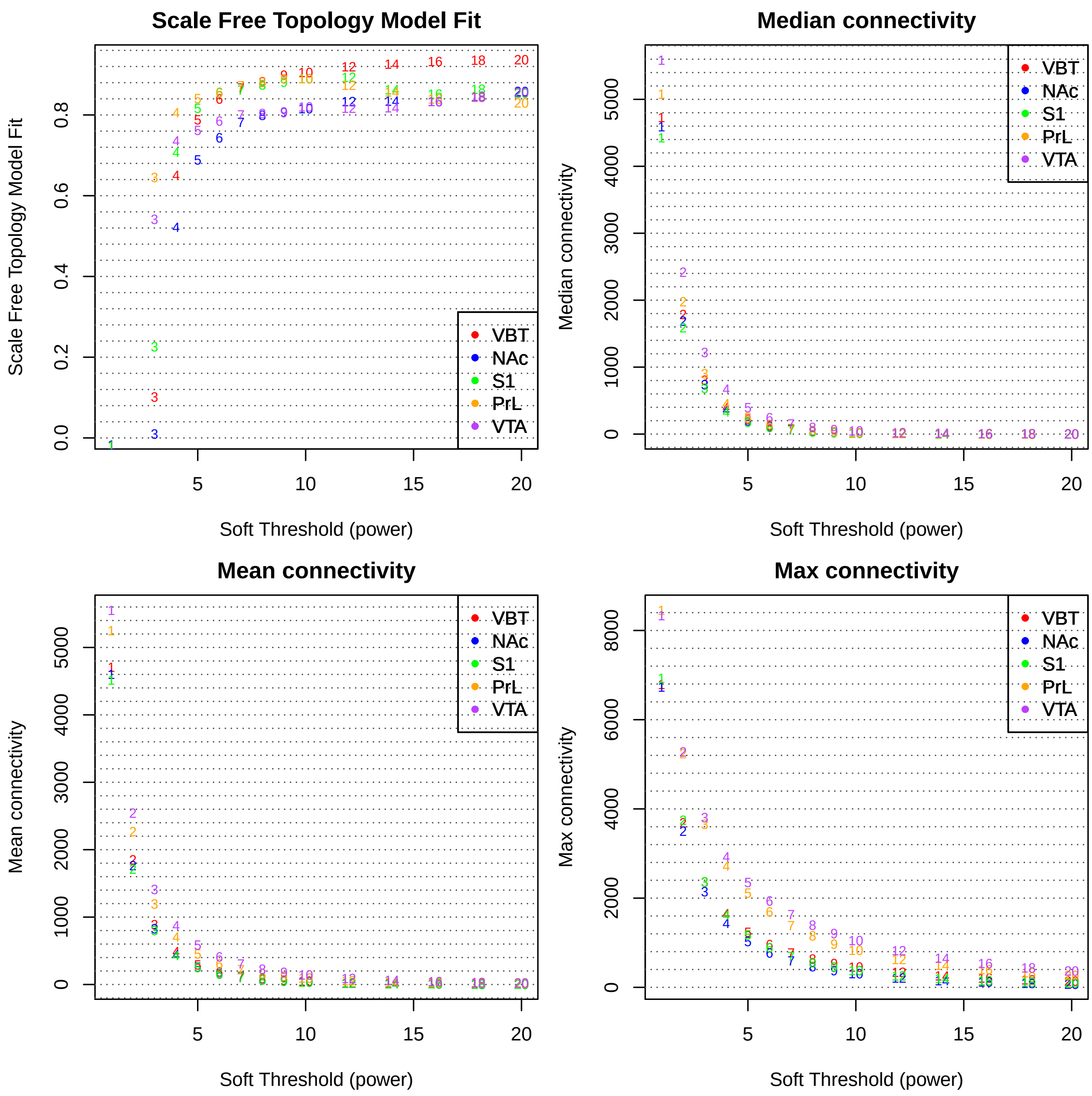


```
# dev.off();
```

##### Construct the WGCNA network for each Brain region separately

*Results for this part is loaded before*

```
powerthres = 9
netPrL = blockwiseModules(multiExpr[[1]]$data, corType = "bicor",
    maxPOutliers = 0.1, power = powerthres, networkType = "signed",
    , maxBlockSize = 19000, minModuleSize = 25, reassignThreshold = 0,
    mergeCutHeight = 0.25, numericLabels = TRUE, minMEtoStay = 0,
    pamRespectsDendro = FALSE, saveTOMs = TRUE, saveTOMFileBase = "TOM.PrL",
    loadTOM = T, verbose = 5)
```

```r
# save(netPrL,file=paste0(workdir,'/net_PrL.Rdata'))

netNAc = blockwiseModules(multiExpr[[2]]$data, corType = "bicor",
    maxPOutliers = 0.1, power = powerthres, networkType = "signed",
    , maxBlockSize = 19000, minModuleSize = 25, reassignThreshold = 0,
    mergeCutHeight = 0.25, numericLabels = TRUE, minMEtoStay = 0,
    pamRespectsDendro = FALSE, saveTOMs = TRUE, saveTOMFileBase = "TOM.NAc",
    loadTOM = T, verbose = 5)
# save(netNAc,file=paste0(workdir,'/net_NAc.Rdata'))

netVBT = blockwiseModules(multiExpr[[3]]$data, corType = "bicor",
    maxPOutliers = 0.1, power = powerthres, networkType = "signed",
    , maxBlockSize = 19000, minModuleSize = 25, reassignThreshold = 0,
    mergeCutHeight = 0.25, numericLabels = TRUE, minMEtoStay = 0,
    pamRespectsDendro = FALSE, saveTOMs = TRUE, saveTOMFileBase = "TOM.VBT",
    loadTOM = T, verbose = 5)
# save(netVBT,file=paste0(workdir,'/net_VBT.Rdata'))

netS1 = blockwiseModules(multiExpr[[5]]$data, corType = "bicor",
    maxPOutliers = 0.1, power = powerthres, networkType = "signed",
    , maxBlockSize = 19000, minModuleSize = 25, reassignThreshold = 0,
    mergeCutHeight = 0.25, numericLabels = TRUE, minMEtoStay = 0,
    pamRespectsDendro = FALSE, saveTOMs = TRUE, saveTOMFileBase = "TOM.S1",
    loadTOM = T, verbose = 5)
# save(netS1,file=paste0(workdir,'/net_S1.Rdata'))

netVTA = blockwiseModules(multiExpr[[4]]$data, corType = "bicor",
    maxPOutliers = 0.1, power = powerthres, networkType = "signed",
    , maxBlockSize = 19000, minModuleSize = 25, reassignThreshold = 0,
    mergeCutHeight = 0.25, numericLabels = TRUE, minMEtoStay = 0,
    pamRespectsDendro = FALSE, saveTOMs = TRUE, saveTOMFileBase = "TOM.VTA",
    loadTOM = T, verbose = 5)
# save(netVTA,file=paste0(workdir,'/net_VTA.Rdata'))
```

##### Module Size of the network

```r
modulesize = list()
modulesize[["PrL"]] = (table(netPrL$colors))
modulesize[["NAc"]] = (table(netNAc$colors))
modulesize[["VBT"]] = table(netVBT$colors)
modulesize[["VTA"]] = (table(netVTA$colors))
modulesize[["S1"]] = (table(netS1$colors))

par(mfrow = c(1, 5))
col = c("springgreen3", "orange", "skyblue", "red2",
    "hotpink")
j = 0
for (i in names(modulesize)) {
    j = j + 1
    bp = barplot(modulesize[[i]][-c(1)], cex.axis = 1,
        cex.names = 0.7, horiz = T, las = 1, col = col[j],
        border = NA, ylab = "Module ID", xlab = "Module size",
        main = i)
```

```r
    text(200, bp - 0.7, modulesize[[i]][-c(1)], pos = 3,
        cex = 0.7)
}
```

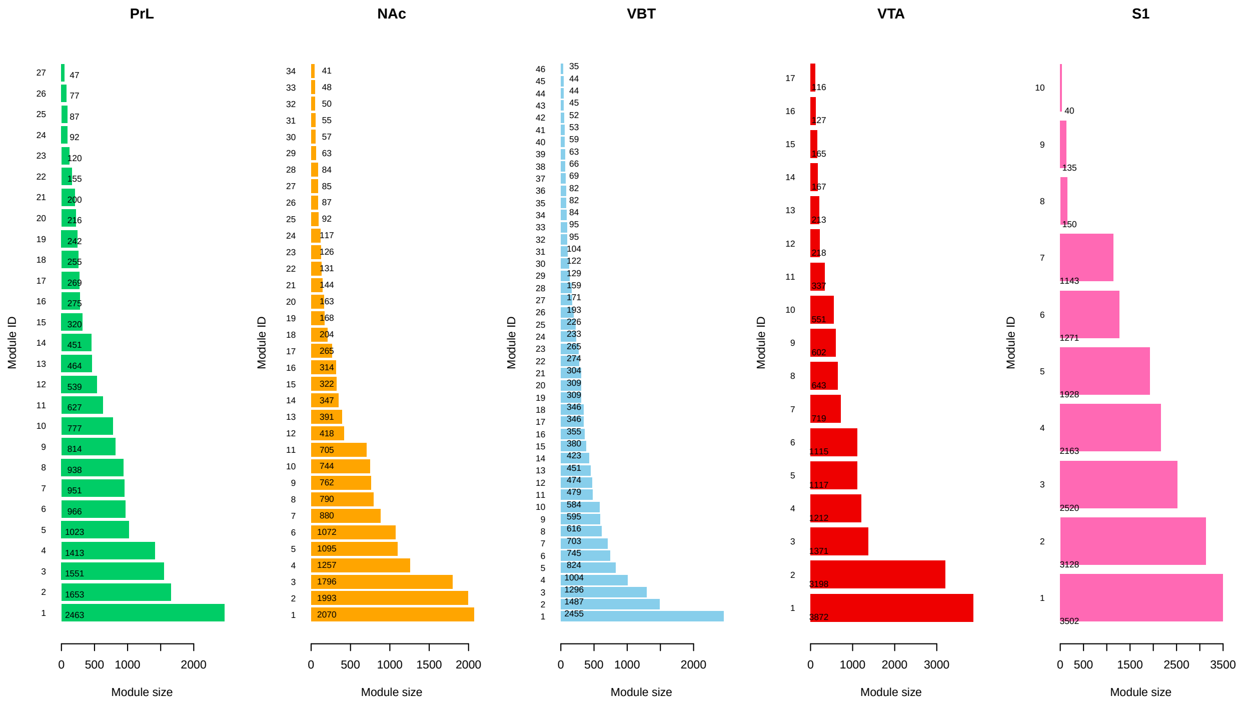


#### Network dendogram figure

```r
plotDendroAndColors(netPrL$dendrograms[[1]], labels2colors(netPrL$colors)[netPrL$blockGenes[[1]]],
    "Module colors", dendroLabels = FALSE, hang = 0.03,
    addGuide = TRUE, guideHang = 0.05, main = "PrL : gene dendrogram and module colors")
```

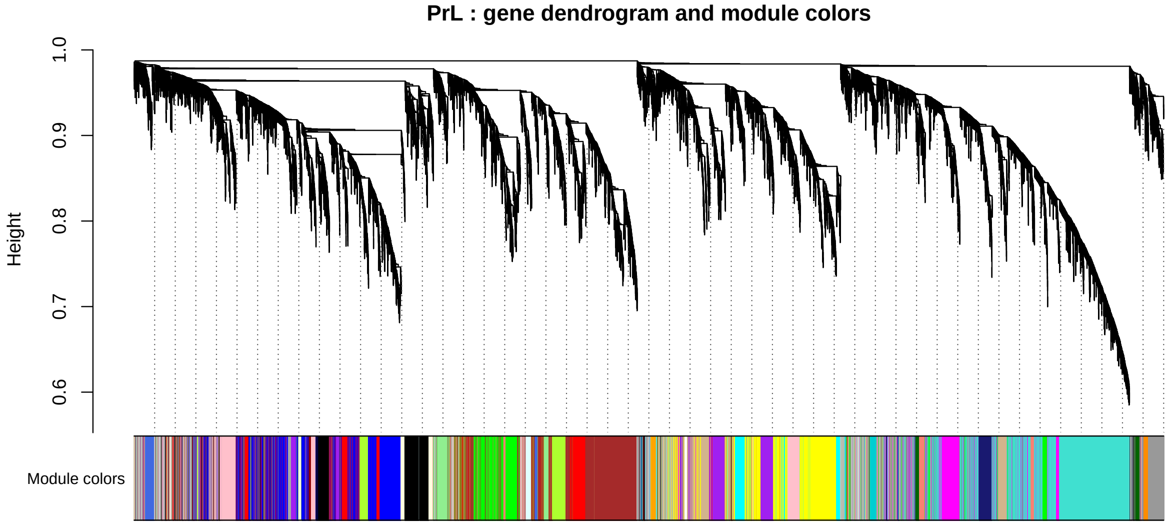

```r
plotDendroAndColors(netNAc$dendrograms[[1]], labels2colors(netNAc$colors)[netNAc$blockGenes[[1]]],
    "Module colors", dendroLabels = FALSE, hang = 0.03,
    addGuide = TRUE, guideHang = 0.05, main = "NAc : gene dendrogram and module colors")
```

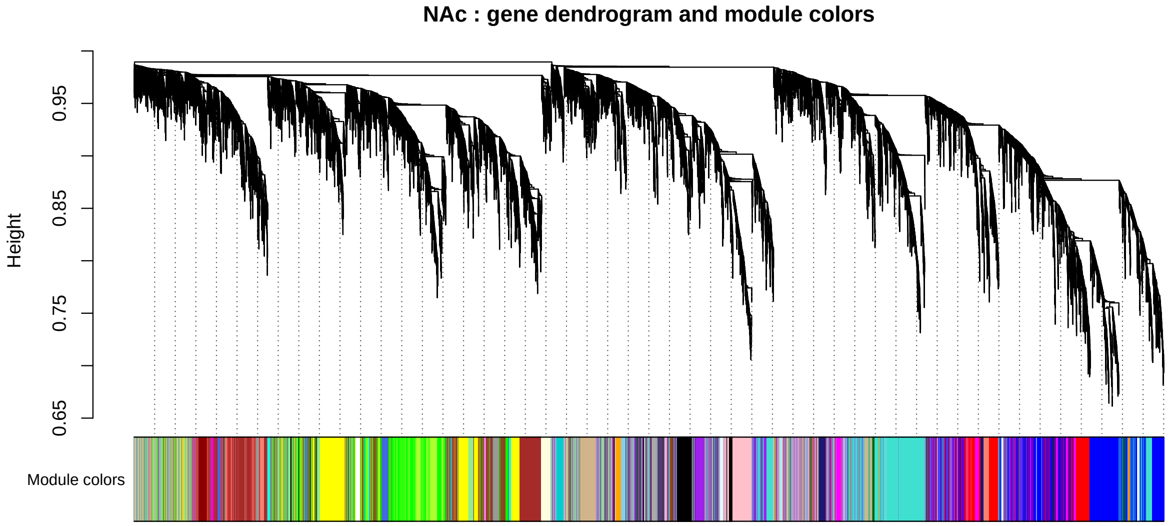


```r
plotDendroAndColors(netVBT$dendrograms[[1]], labels2colors(netVBT$colors)[netVBT$blockGenes[[1]]],
    "Module colors", dendroLabels = FALSE, hang = 0.03,
    addGuide = TRUE, guideHang = 0.05, main = "VBT : gene dendrogram and module colors")
```

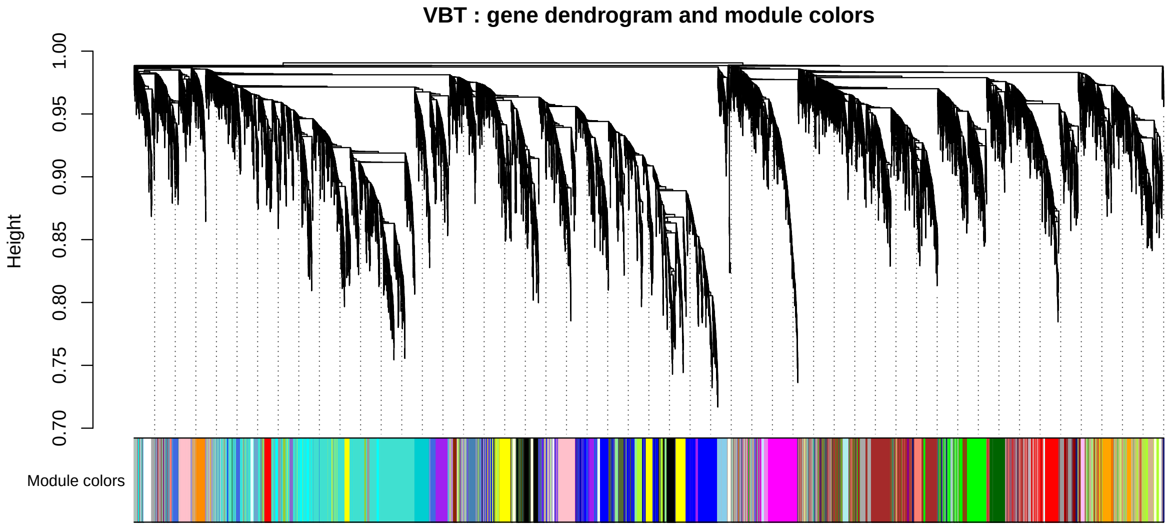


```r
plotDendroAndColors(netVTA$dendrograms[[1]], labels2colors(netVTA$colors)[netVTA$blockGenes[[1]]],
    "Module colors", dendroLabels = FALSE, hang = 0.03,
    addGuide = TRUE, guideHang = 0.05, main = "VTA : gene dendrogram and module colors")
```

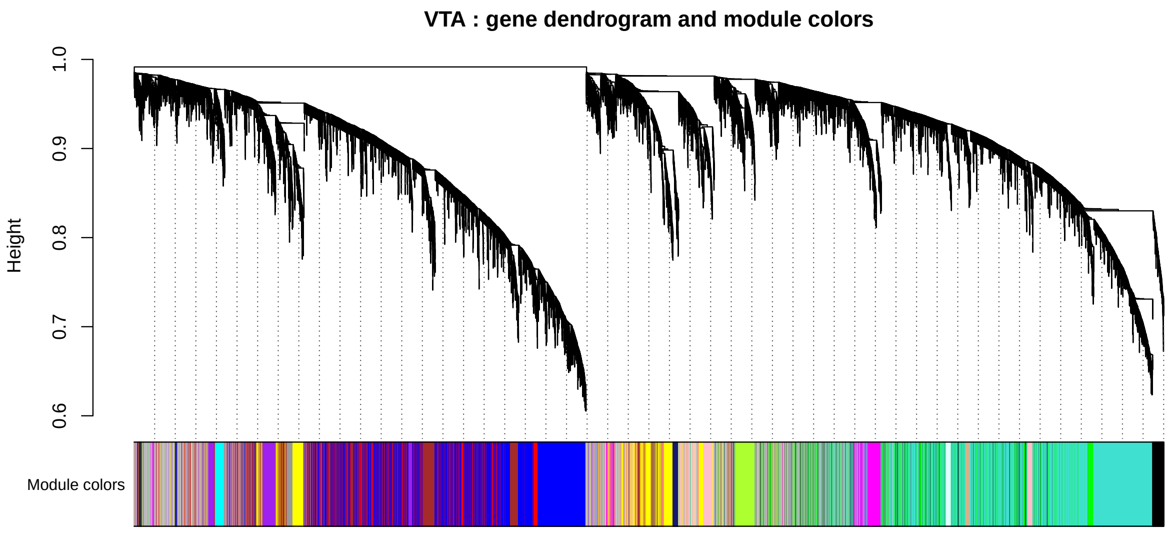


```r
plotDendroAndColors(netS1$dendrograms[[1]], labels2colors(netS1$colors)[netS1$blockGenes[[1]]],
    "Module colors", dendroLabels = FALSE, hang = 0.03,
    addGuide = TRUE, guideHang = 0.05, main = "S1 :gene dendrogram and module colors")
```

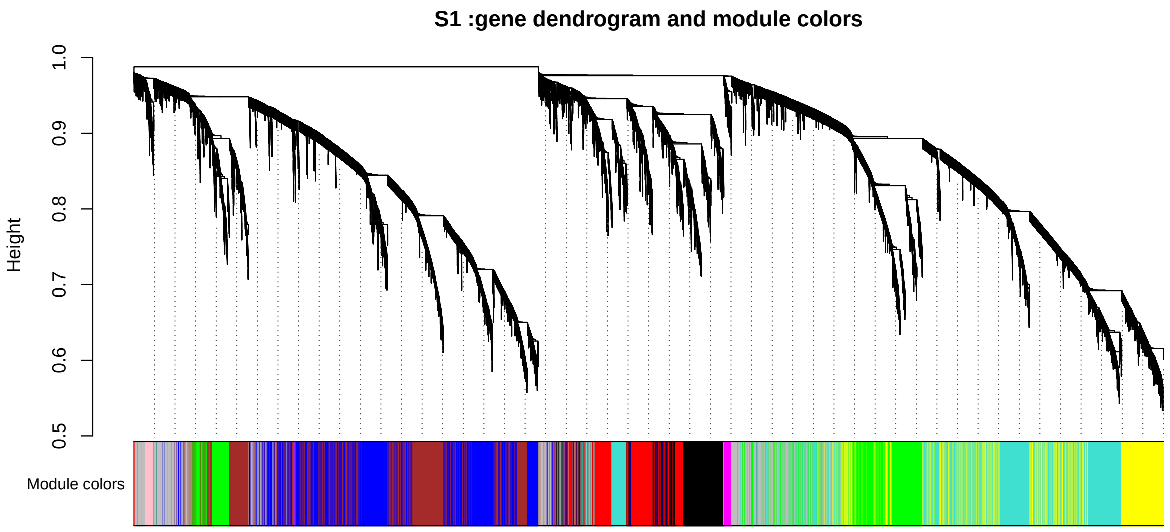


##### Module-Module similarities : gene overlap

```r
netcol = list()
netcol[["PrL"]] = netPrL$colors
netcol[["NAc"]] = netNAc$colors
netcol[["VBT"]] = netVBT$colors
netcol[["VTA"]] = netVTA$colors
netcol[["S1"]] = netS1$colors
j = 0
for (i in celltype) {
    j = j + 1
```

```
    MEs.AvE = moduleEigengenes(multiExpr[[j]]$data,
        netcol[[i]])$averageExpr
    MEs.Eig = moduleEigengenes(multiExpr[[j]]$data,
        netcol[[i]])$eigengenes
    MET.AvE = orderMEs(MEs.AvE)
    MET.Eig = orderMEs(MEs.Eig)

    plotEigengeneNetworks(MET.AvE, paste0(i, "-Average Expr adjacency heatmap"),
        marDendro = c(0, 4, 1, 2), marHeatmap = c(3,
            4, 2, 2), cex.lab = 1.2, xLabelsAngle = 90)
    print(plotEigengeneNetworks(MET.Eig, paste0(i,
        "-Eigengene adjacency heatmap"), marDendro = c(0,
        4, 1, 4), marHeatmap = c(3, 4, 2, 2), cex.lab = 1.2,
        xLabelsAngle = 90))
}
```

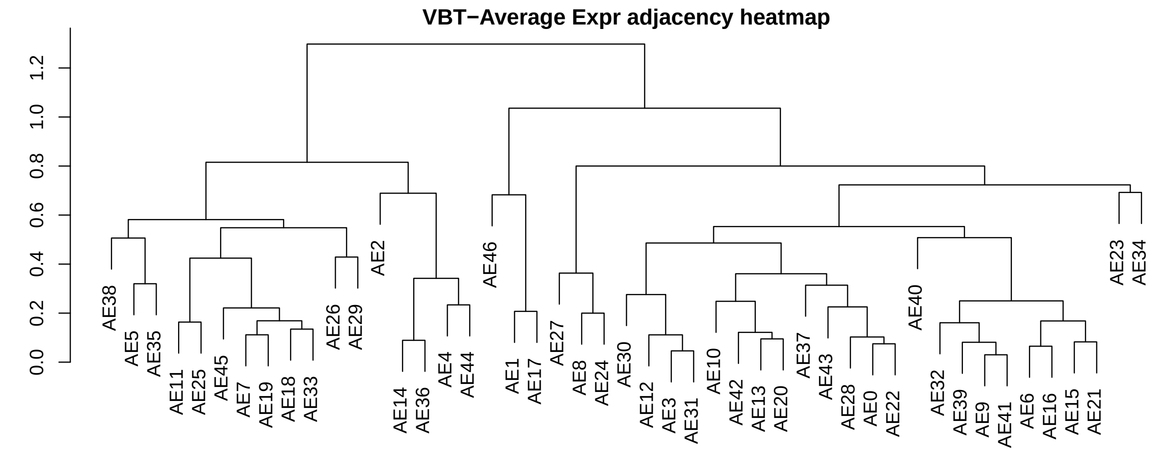


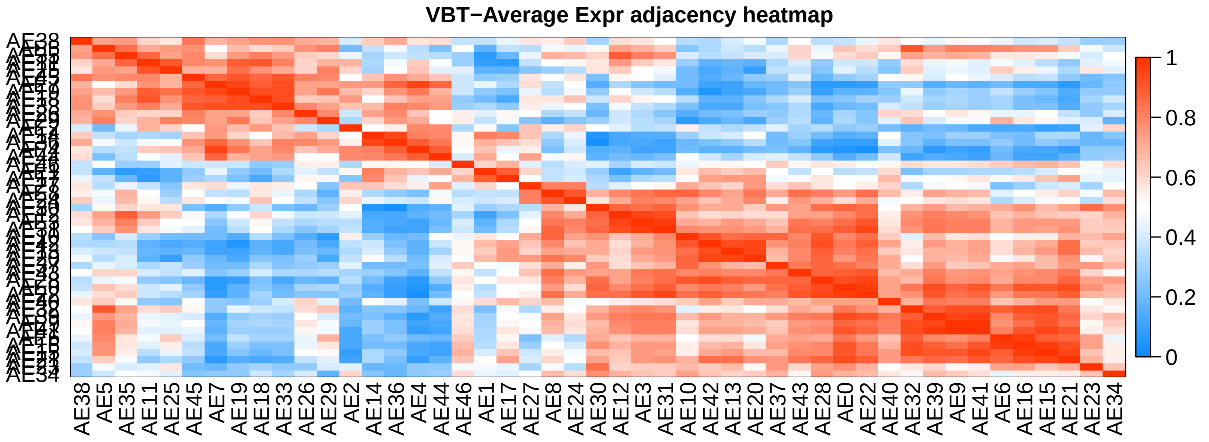

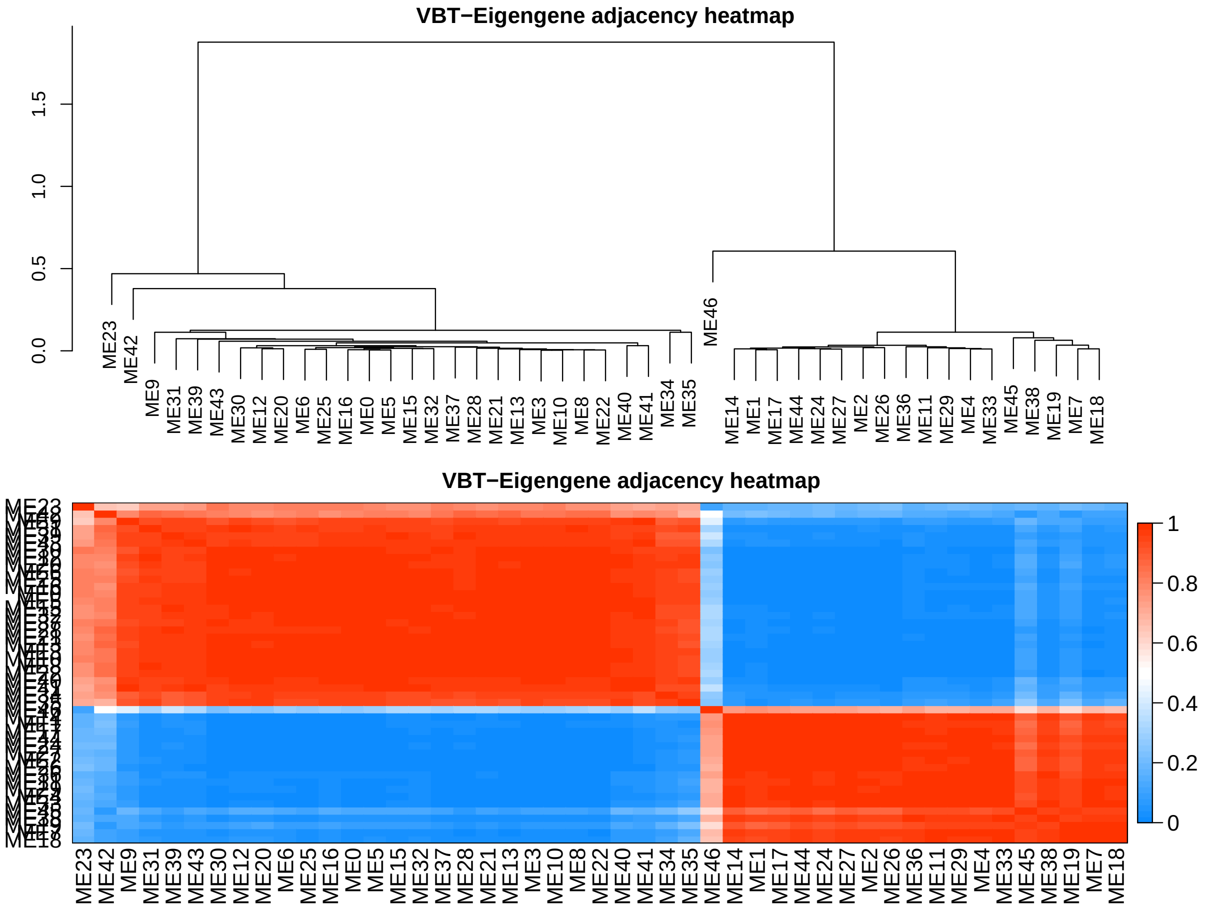
VBT−Eigengene adjacency heatmap
0.0
0.5
1.0
1.5
ME23 ME42 ME9 ME31 ME39 ME43 ME30 ME12 ME20 ME6 ME25 ME16 ME0 ME5 ME15 ME32 ME37 ME28 ME21 ME13 ME3 ME10 ME8 ME22 ME40 ME41 ME34 ME35 ME46 ME14 ME1 ME17 ME44 ME24 ME27 ME2 ME26 ME36 ME11 ME29 ME4 ME33 ME45 ME38 ME19 ME7 ME18
VBT−Eigengene adjacency heatmap
ME23 ME42 ME9 ME31 ME39 ME43 ME30 ME12 ME20 ME6 ME25 ME16 ME0 ME5 ME15 ME32 ME37 ME28 ME21 ME13 ME3 ME10 ME8 ME22 ME40 ME41 ME34 ME35 ME46 ME14 ME1 ME17 ME44 ME24 ME27 ME2 ME26 ME36 ME11 ME29 ME4 ME33 ME45 ME38 ME19 ME7 ME18
1
0.8
0.6
0.4
0.2
0

```
## NULL
```

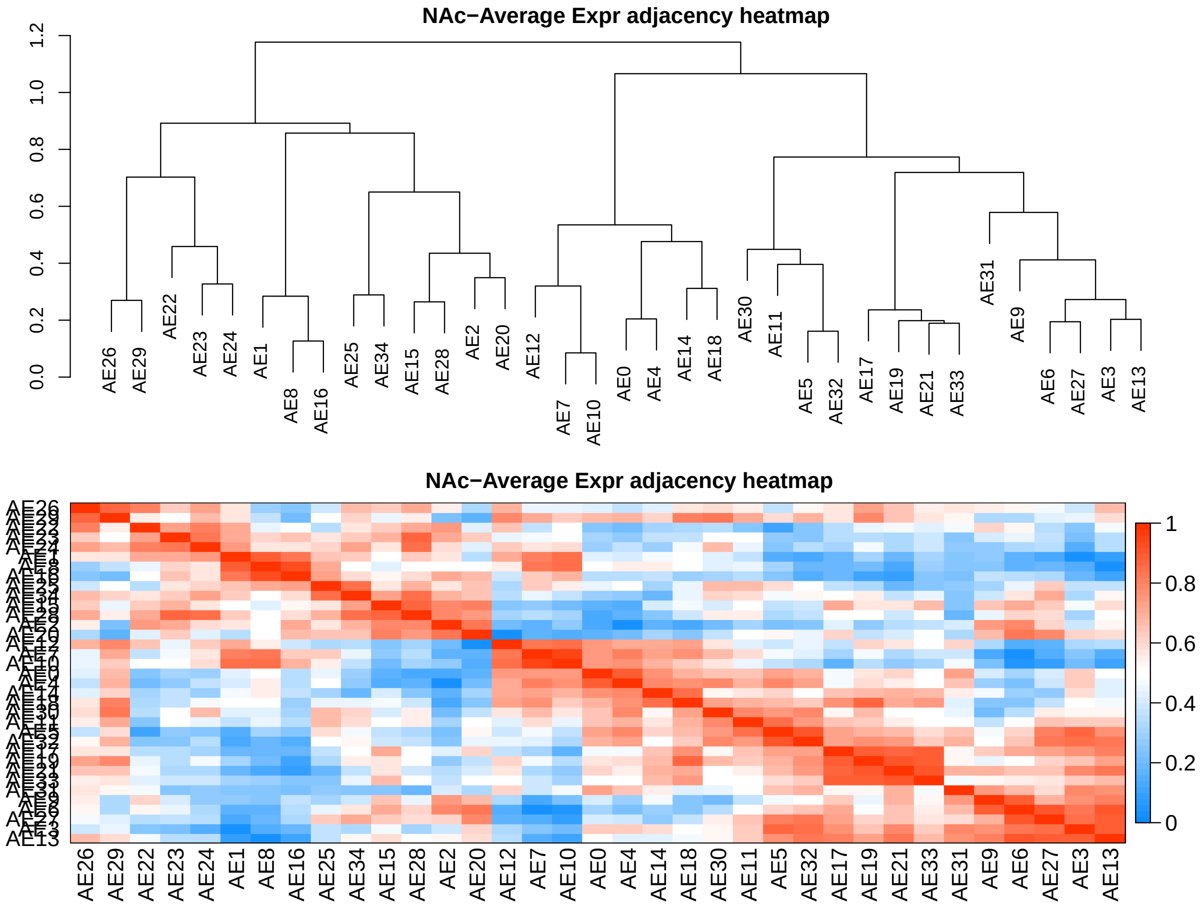
NAc−Average Expr adjacency heatmap
0.0
0.2
0.4
0.6
0.8
1.0
1.2
AE26 AE29 AE22 AE23 AE24 AE1 AE8 AE16 AE25 AE34 AE15 AE28 AE2 AE20 AE12 AE7 AE10 AE0 AE4 AE14 AE18 AE30 AE11 AE5 AE32 AE17 AE19 AE21 AE33 AE31 AE9 AE6 AE27 AE3 AE13
NAc−Average Expr adjacency heatmap
AE26 AE29 AE22 AE23 AE24 AE1 AE8 AE16 AE25 AE34 AE15 AE28 AE2 AE20 AE12 AE7 AE10 AE0 AE4 AE14 AE18 AE30 AE11 AE5 AE32 AE17 AE19 AE21 AE33 AE31 AE9 AE6 AE27 AE3 AE13
1
0.8
0.6
0.4
0.2
0

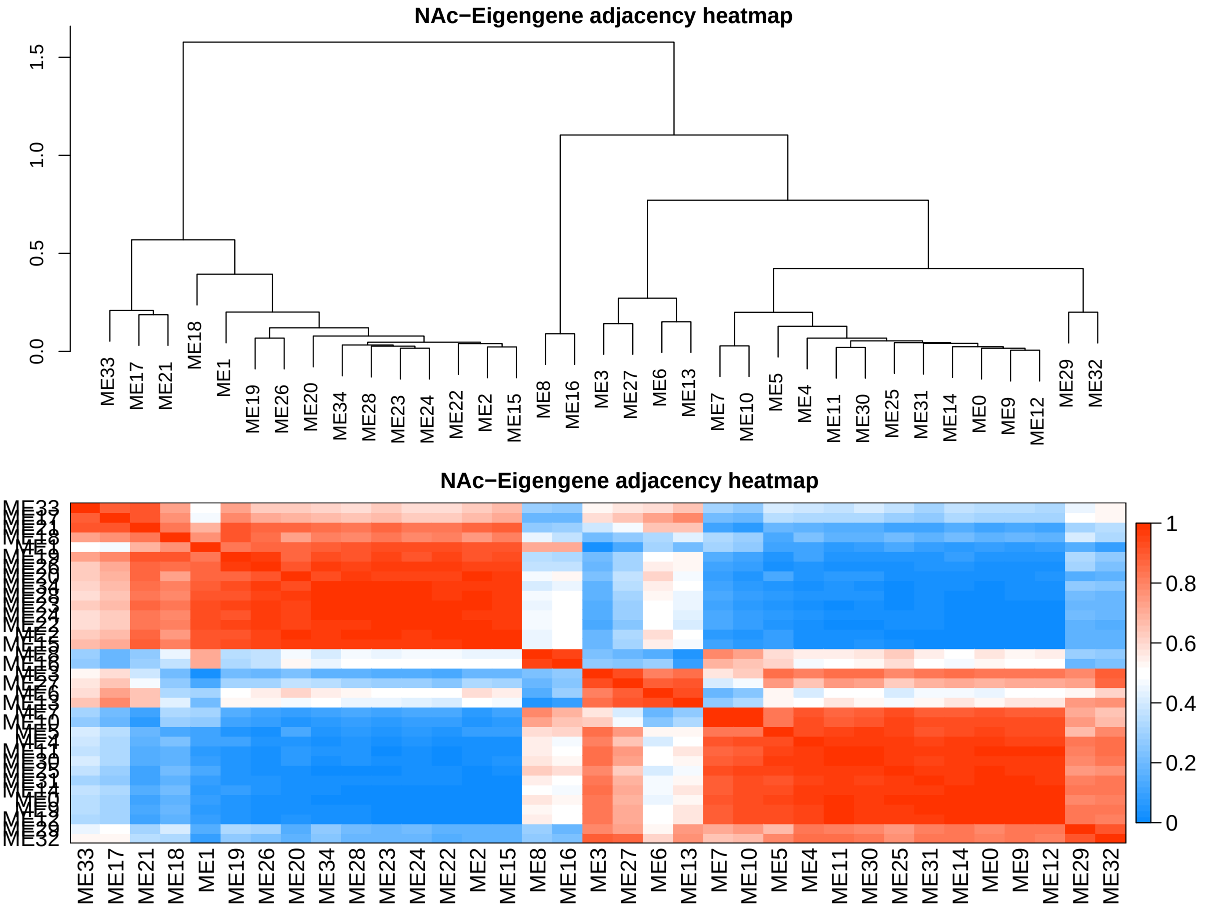
NAc−Eigengene adjacency heatmap
1.5
1.0
0.5
0.0
ME33 ME17 ME21 ME18 ME1 ME19 ME26 ME20 ME34 ME28 ME23 ME24 ME22 ME2 ME15 ME8 ME16 ME3 ME27 ME6 ME13 ME7 ME10 ME5 ME4 ME11 ME30 ME25 ME31 ME14 ME0 ME9 ME12 ME29 ME32
NAc−Eigengene adjacency heatmap
ME33 ME17 ME21 ME18 ME1 ME19 ME26 ME20 ME34 ME28 ME23 ME24 ME22 ME2 ME15 ME8 ME16 ME3 ME27 ME6 ME13 ME7 ME10 ME5 ME4 ME11 ME30 ME25 ME31 ME14 ME0 ME9 ME12 ME29 ME32
1
0.8
0.6
0.4
0.2
0

```
## NULL
```

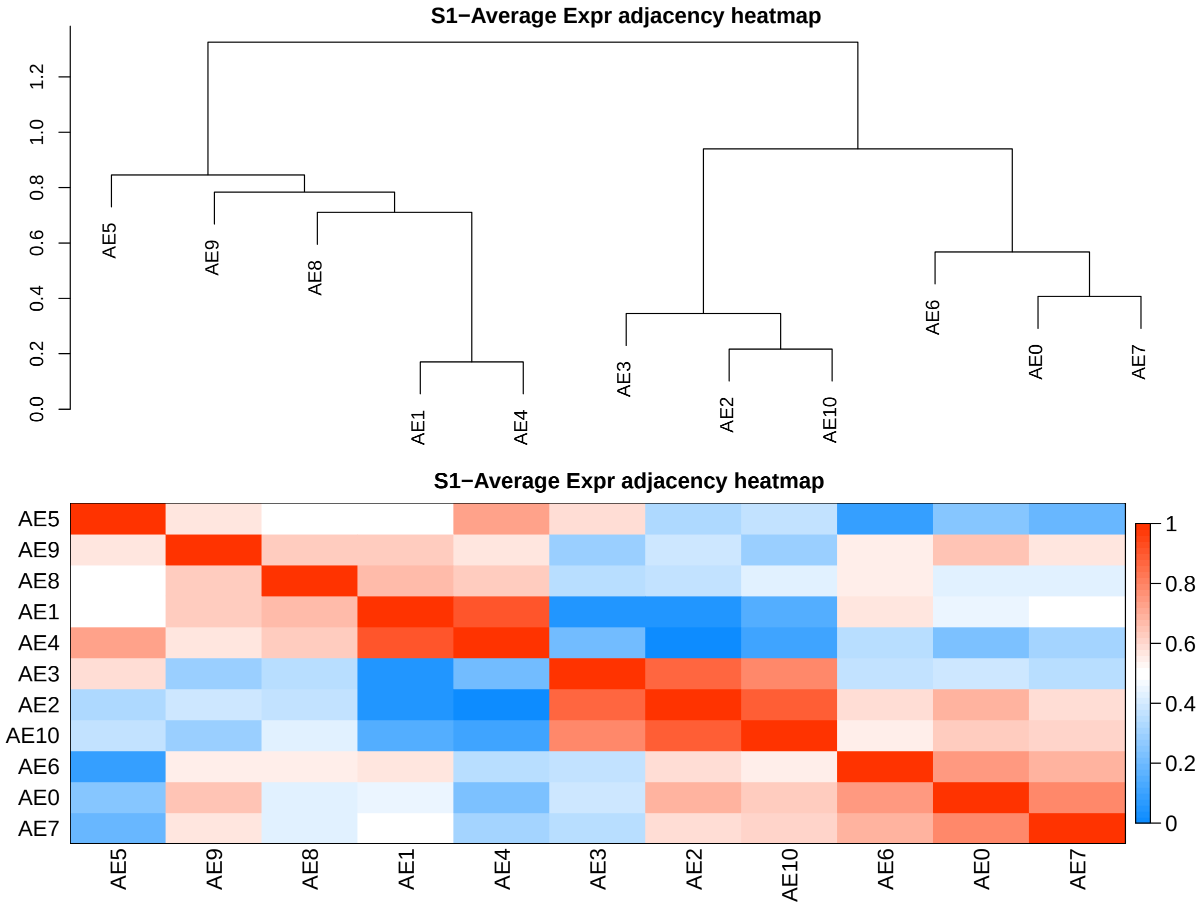
S1−Average Expr adjacency heatmap
AE5
AE9
AE8
AE1
AE4
AE3
AE2
AE10
AE6
AE0
AE7
0.0
0.2
0.4
0.6
0.8
1.0
1.2
S1−Average Expr adjacency heatmap
1
0.8
0.6
0.4
0.2
0

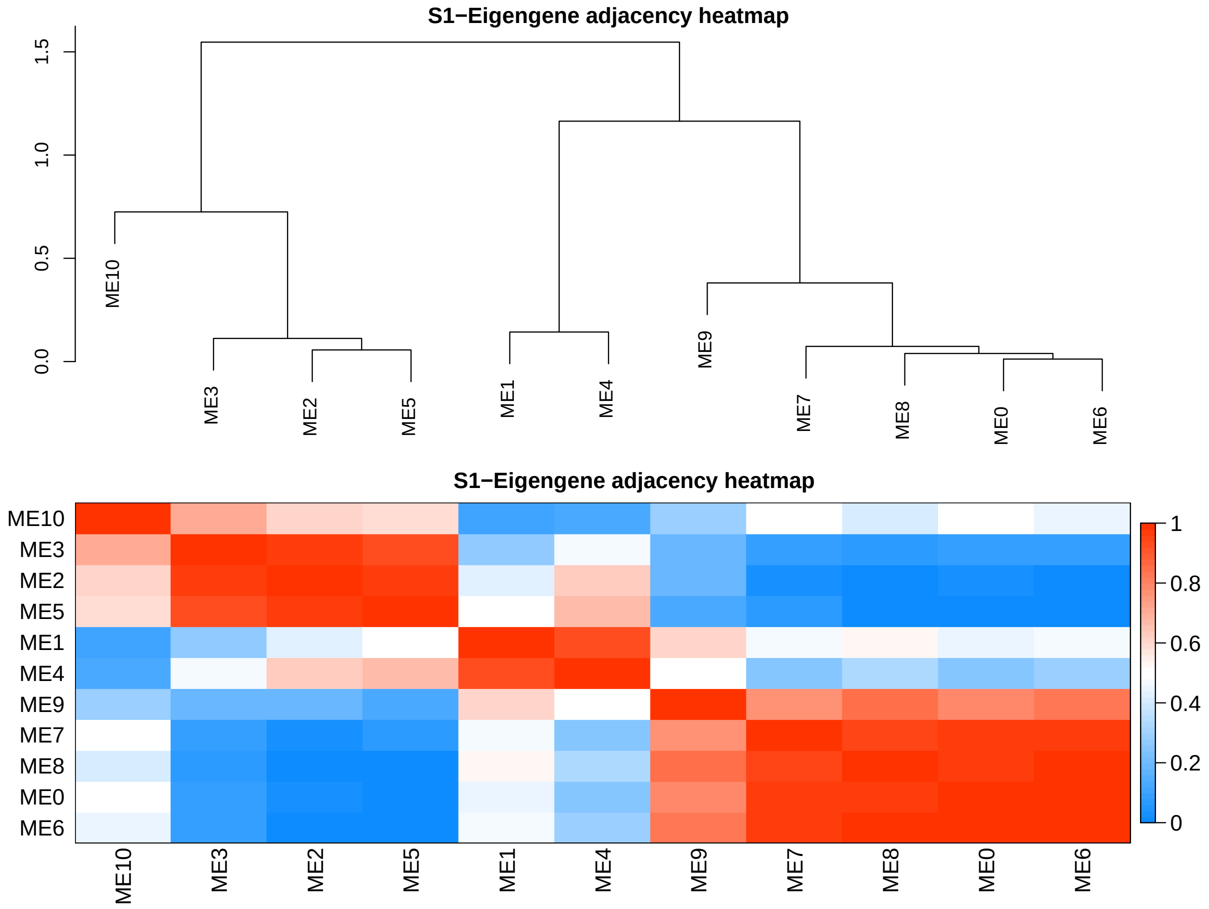
S1−Eigengene adjacency heatmap
0.0
0.5
1.0
1.5
ME10
ME3
ME2
ME5
ME1
ME4
ME9
ME7
ME8
ME0
ME6
S1−Eigengene adjacency heatmap
1
0.8
0.6
0.4
0.2
0

```
## NULL
```

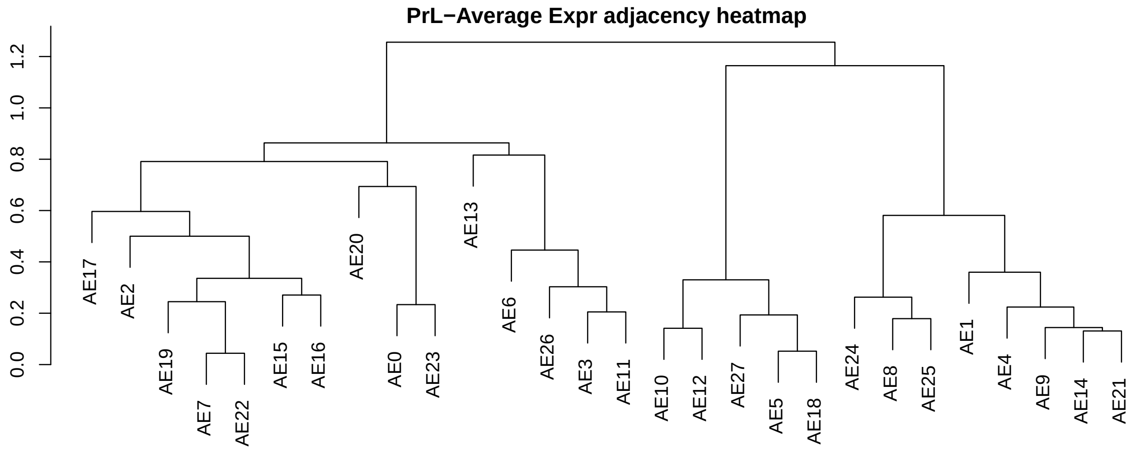
PrL−Average Expr adjacency heatmap
0.0
0.2
0.4
0.6
0.8
1.0
1.2
AE17
AE2
AE19
AE7
AE22
AE15
AE16
AE20
AE0
AE23
AE13
AE6
AE26
AE3
AE11
AE10
AE12
AE27
AE5
AE18
AE24
AE8
AE25
AE1
AE4
AE9
AE14
AE21

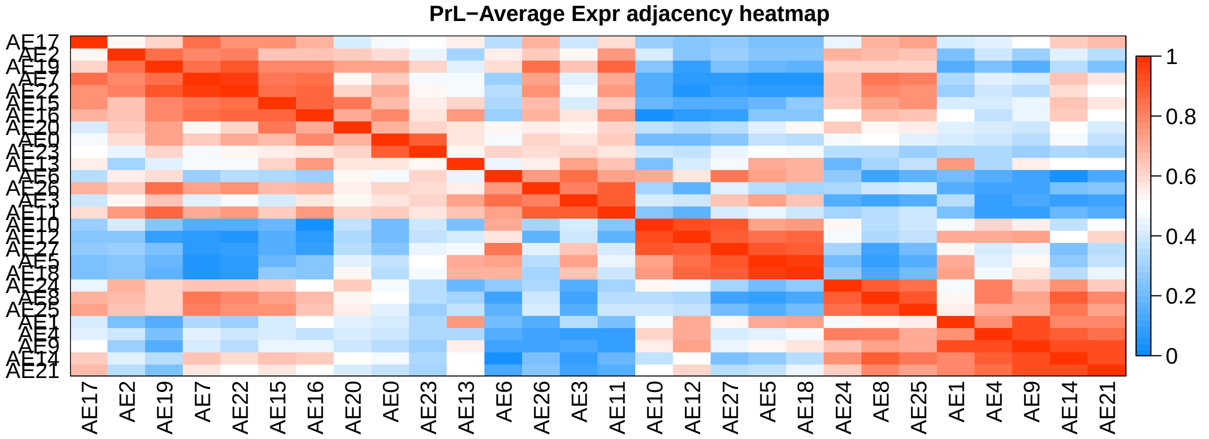
PrL−Average Expr adjacency heatmap
AE17
AE2
AE19
AE7
AE22
AE15
AE16
AE20
AE0
AE23
AE13
AE6
AE26
AE3
AE11
AE10
AE12
AE27
AE5
AE18
AE24
AE8
AE25
AE1
AE4
AE9
AE14
AE21
1
0.8
0.6
0.4
0.2
0

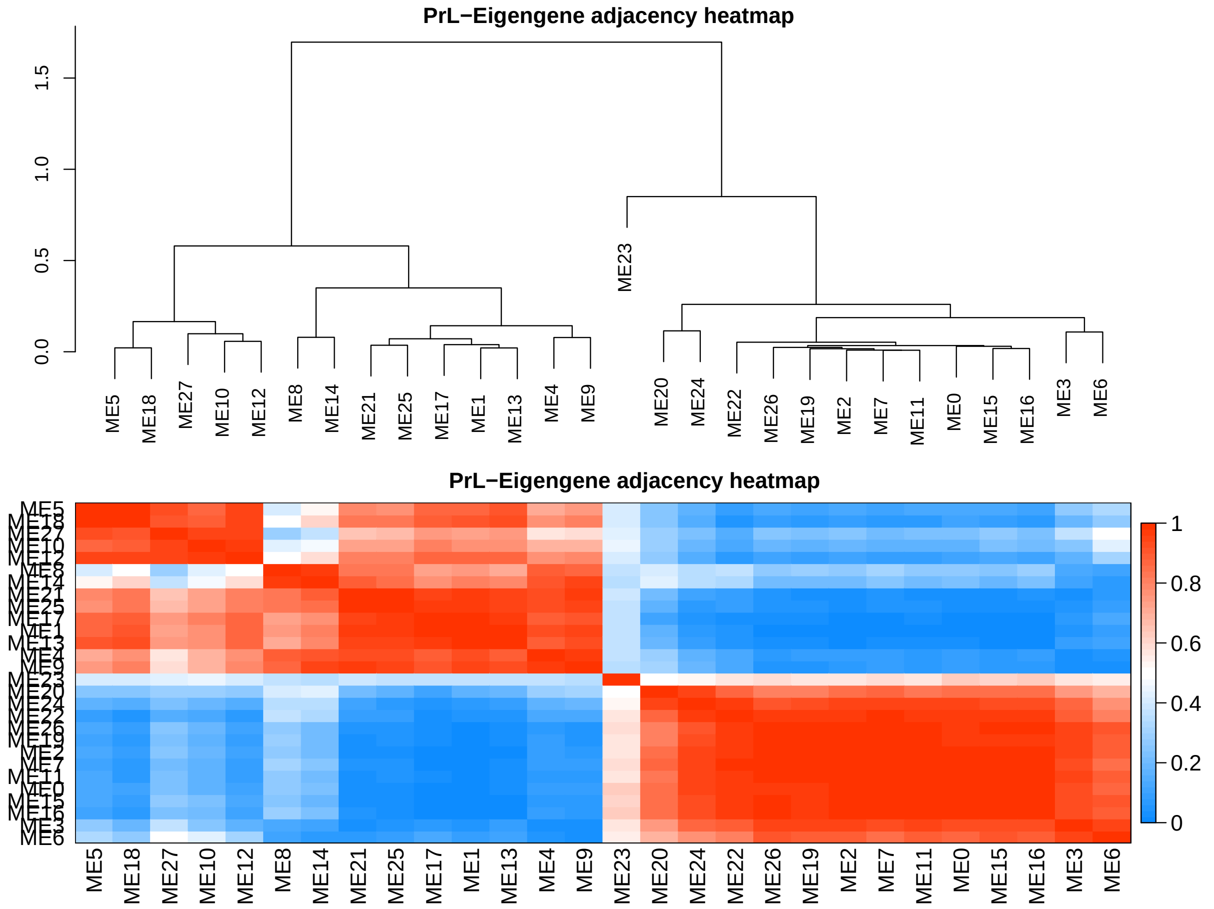
PrL−Eigengene adjacency heatmap
PrL−Eigengene adjacency heatmap
1
0.8
0.6
0.4
0.2
0

```
## NULL
```

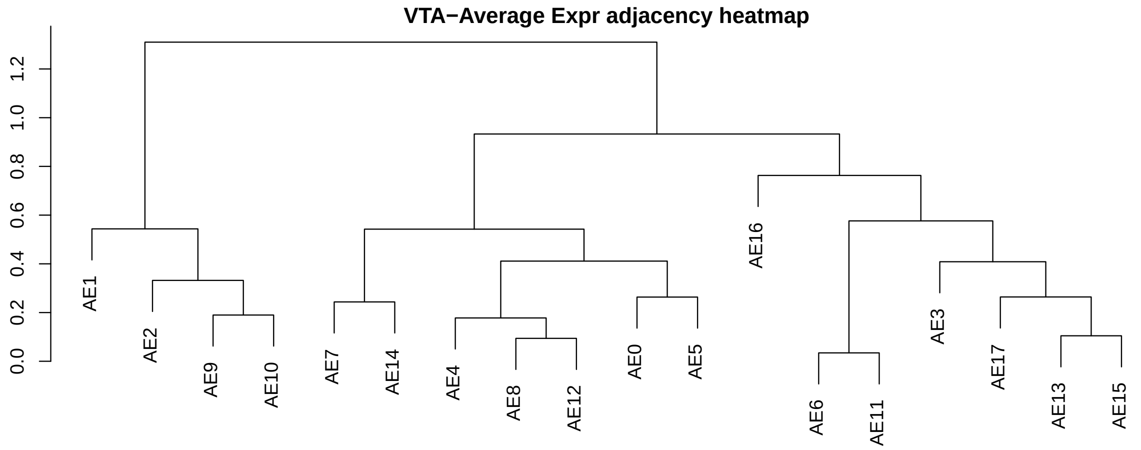
VTA−Average Expr adjacency heatmap
0.0
0.2
0.4
0.6
0.8
1.0
1.2
AE1
AE2
AE9
AE10
AE7
AE14
AE4
AE8
AE12
AE0
AE5
AE16
AE6
AE11
AE3
AE17
AE13
AE15

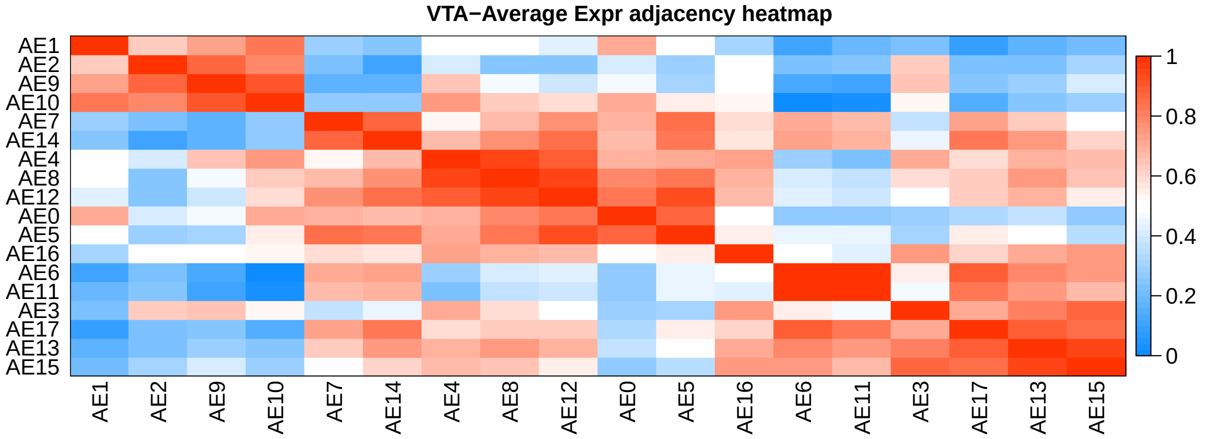
VTA−Average Expr adjacency heatmap
AE1
AE2
AE9
AE10
AE7
AE14
AE4
AE8
AE12
AE0
AE5
AE16
AE6
AE11
AE3
AE17
AE13
AE15
1
0.8
0.6
0.4
0.2
0

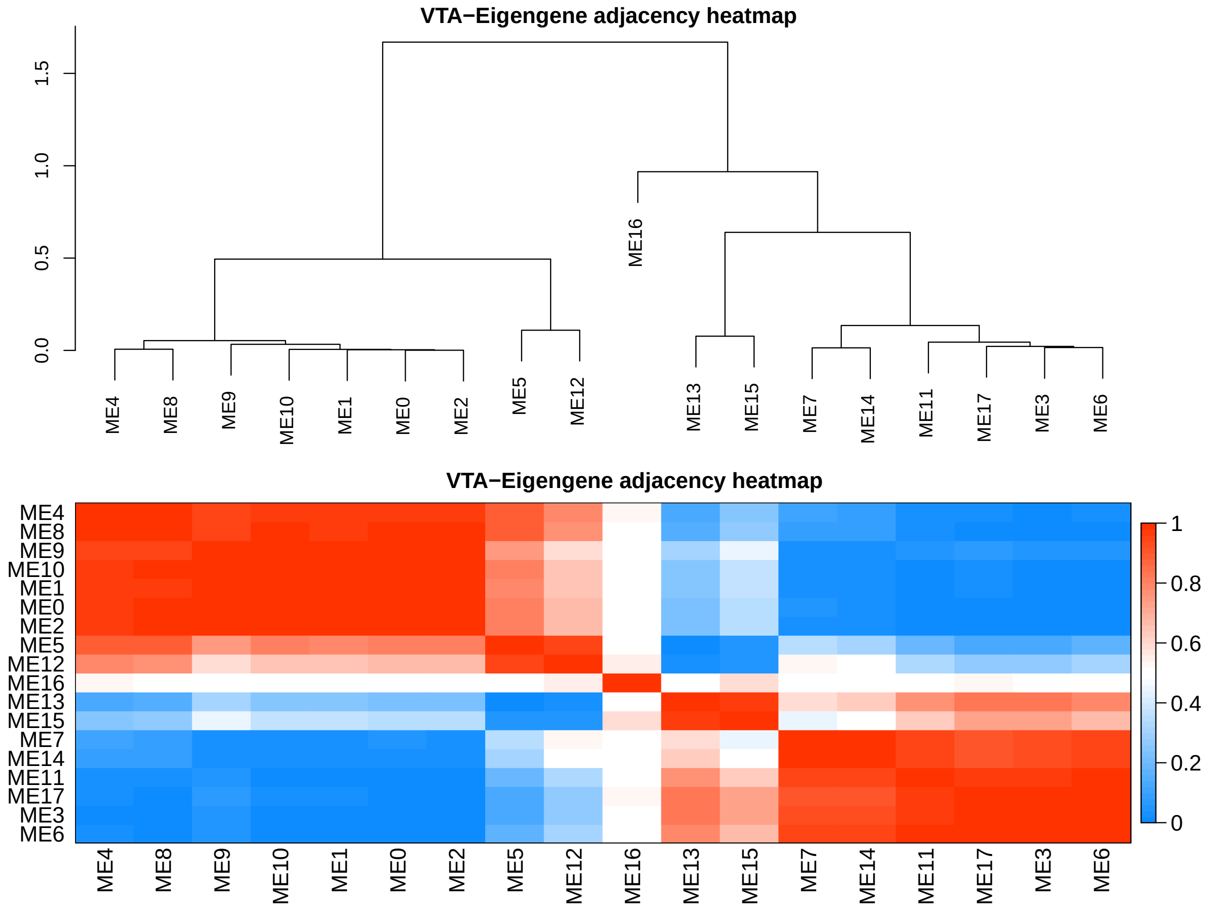


```
## NULL
```

#### Relate Modules to traits : limma

```
limmatraits = list()
MEslist = list()

for (i in celltype) {
    MEs = moduleEigengenes(datExpr[grep(i, rownames(datExpr)),
        ], netcol[[i]])
    MEs = MEs$averageExpr
    MEslist[[i]] = list(dataAE = MEs, nmod = ncol(MEs))
    meta1 = meta[which(meta$celltype == i), ]
    design2 = model.matrix(~0 + meta1$Group)
    colnames(design2) = gsub(paste0("meta1\\$Group",
        i, "_"), "", colnames(design2))

    contr = makeContrasts(M = (MF - MW), F = (FF -
        FW), C = (MF + FF)/2 - (MW + FW)/2, levels = design2)

    fitME = lmFit(t(MEs), design = design2)
    fitME <- eBayes(fitME)
```

```
    fitME2 = contrasts.fit(fitME, contr)
    fitME2 = eBayes(fitME2)
    res2 = topTable(fitME2, coef = 1, number = Inf)
    res2 = merge(res2, topTable(fitME2, coef = 2, number = Inf),
        by = 0)
    res2 = merge(res2, topTable(fitME2, coef = 3, number = Inf),
        by.x = 1, by.y = 0)

    n = ncol(contr)
    limma.ME.pvals = res2[, c(1:n * 6 - 1)]
    limma.ME.logFC = res2[, c(1:n * 6 - 4)]
    colnames(limma.ME.pvals) = colnames(limma.ME.logFC) = colnames(contr)
    rownames(limma.ME.pvals) = rownames(limma.ME.logFC) = res2[,
        1]
    colnames(limma.ME.pvals) = paste("P.Value", colnames(limma.ME.pvals),
        sep = "_")
    colnames(limma.ME.logFC) = paste("logFC", colnames(limma.ME.logFC),
        sep = "_")

    limmatraits[[i]] = list(data = res2, pval = limma.ME.pvals,
        logFC = limma.ME.logFC)
}

siglimmatraits = lapply(1:length(limmatraits), function(i) {
    keep = rowSums(limmatraits[[names(limmatraits)[i]]]$pval <
        0.05) > 0
    x = limmatraits[[names(limmatraits)[i]]]$pval[keep,
        ]
    y = limmatraits[[names(limmatraits)[i]]]$logFC[keep,
        ]
    return(list(pval = x, logFC = y))
})
names(siglimmatraits) = names(limmatraits)
```

##### Modules with significant DE (Fentanyl ~ Water)

Significant case with pvalue<=0.05 are marked red. The model in each module is build using the average expression of the eigen genes. LogFC=log(F/W) >0 => F/W>1 and log(F/W) <0 => F/W <1

```
for (i in celltype) {
    dt = cbind(signif(siglimmatraits[[i]]$pval, 3),
        signif(siglimmatraits[[i]]$logFC, 3))
    print(dt %>% mutate(Module = gsub("AE", "M", rownames(dt)),
        P.Value_M = cell_spec(P.Value_M, "html", color = ifelse(P.Value_M <=
            0.05, "red", "black")), P.Value_F = cell_spec(P.Value_F,
            "html", color = ifelse(P.Value_F <= 0.05,
                "red", "black")), P.Value_C = cell_spec(P.Value_C,
            "html", color = ifelse(P.Value_C <= 0.05,
                "red", "black"))) %>% dplyr::select(Module,
        P.Value_M, P.Value_F, P.Value_C, logFC_M, logFC_F,
        logFC_C) %>% knitr::kable(caption = paste0("Significant cases for ",
        i)) %>% kable_styling(c("striped", "condensed"),
```

```
        full_width = F, position = "left") %>% scroll_box(width = "60%",
        height = "200px"))
    cat("\n")
}
```

Table 1: Significant cases for VBT

| Module | P.Value_M | P.Value_F | P.Value_C | logFC_M | logFC_F | logFC_C |
|---|---|---|---|---|---|---|
| M1 | 0.0622 | 0.132 | 0.0206 | 0.2700 | 0.2150 | 0.243 |
| M11 | 0.1 | 0.0241 | 0.00782 | -0.2710 | -0.3820 | -0.327 |
| M19 | 0.0486 | 0.309 | 0.0376 | -0.4700 | -0.2340 | -0.352 |
| M25 | 0.0485 | 0.0668 | 0.00953 | -0.2950 | -0.2720 | -0.284 |
| M34 | 0.0433 | 0.638 | 0.251 | 0.3950 | -0.0878 | 0.153 |
| M35 | 0.192 | 0.031 | 0.0171 | -0.2840 | -0.4860 | -0.385 |
| M44 | 0.878 | 0.0472 | 0.125 | 0.0254 | 0.3430 | 0.184 |

Table 2: Significant cases for NAc

| Module | P.Value_M | P.Value_F | P.Value_C | logFC_M | logFC_F | logFC_C |
|---|---|---|---|---|---|---|
| M1 | 0.047 | 0.534 | 0.309 | 0.401 | -0.120 | 0.1400 |
| M13 | 0.0242 | 0.291 | 0.354 | -0.684 | 0.306 | -0.1890 |
| M16 | 0.00022 | 0.1 | 0.0693 | 0.802 | -0.312 | 0.2450 |
| M17 | 0.0391 | 0.284 | 0.446 | -0.426 | 0.213 | -0.1060 |
| M19 | 0.0372 | 0.317 | 0.408 | -0.357 | 0.165 | -0.0961 |
| M21 | 0.0297 | 0.273 | 0.405 | -0.408 | 0.197 | -0.1050 |
| M8 | 0.000511 | 0.456 | 0.0286 | 0.829 | -0.154 | 0.3370 |

Table 3: Significant cases for S1

| Module | P.Value_M | P.Value_F | P.Value_C | logFC_M | logFC_F | logFC_C |
|---|---|---|---|---|---|---|
| M1 | 0.776 | 0.0306 | 0.178 | -0.0525 | 0.4040 | 0.176 |
| M10 | 0.562 | 0.00195 | 0.0092 | -0.2100 | -1.2200 | -0.716 |
| M2 | 0.779 | 0.00433 | 0.0248 | -0.0507 | -0.5500 | -0.301 |
| M4 | 0.336 | 0.00383 | 0.00779 | 0.1910 | 0.6080 | 0.400 |
| M5 | 0.0941 | 0.223 | 0.0453 | 0.2350 | 0.1580 | 0.196 |
| M7 | 0.00692 | 0.164 | 0.00485 | -0.3730 | -0.1670 | -0.270 |
| M8 | 0.0115 | 0.844 | 0.0423 | 0.2230 | 0.0149 | 0.119 |

Table 4: Significant cases for PrL

| Module | P.Value_M | P.Value_F | P.Value_C | logFC_M | logFC_F | logFC_C |
|---|---|---|---|---|---|---|
| M10 | 0.115 | 0.15 | 0.0379 | 0.341 | 0.309 | 0.325 |
| M11 | 0.0306 | 0.0782 | 0.00786 | -0.307 | -0.245 | -0.276 |
| M12 | 0.07 | 0.026 | 0.00633 | 0.463 | 0.582 | 0.522 |
| M16 | 0.123 | 0.103 | 0.0296 | -0.268 | -0.285 | -0.277 |
| M19 | 0.0431 | 0.0321 | 0.00503 | -0.377 | -0.403 | -0.390 |
| M22 | 0.366 | 0.0354 | 0.036 | -0.168 | -0.411 | -0.289 |

| Module | P.Value_M | P.Value_F | P.Value_C | logFC_M | logFC_F | logFC_C |
|---|---|---|---|---|---|---|
| M26 | 0.153 | 0.067 | 0.0253 | -0.267 | -0.348 | -0.308 |
| M9 | 0.166 | 0.103 | 0.0379 | 0.331 | 0.393 | 0.362 |

Table 5: Significant cases for VTA

| Module | P.Value_M | P.Value_F | P.Value_C | logFC_M | logFC_F | logFC_C |
|---|---|---|---|---|---|---|
| M10 | 9.97e-05 | 0.0477 | 8.19e-05 | -1.3400 | -0.534 | -0.935 |
| M11 | 0.00351 | 0.361 | 0.00639 | 0.9810 | 0.257 | 0.619 |
| M17 | 0.0227 | 0.441 | 0.0302 | 0.5220 | 0.155 | 0.339 |
| M6 | 0.00115 | 0.265 | 0.00206 | 0.9610 | 0.269 | 0.615 |
| M7 | 0.339 | 0.0209 | 0.0248 | 0.0833 | 0.203 | 0.143 |
| M9 | 0.0111 | 0.0708 | 0.00343 | -0.4360 | -0.278 | -0.357 |

##### Boxplot for significant Modules

```
for (i in celltype) {
    mod.x = rownames(siglimmatraits[[i]]$pval)
    par(mfrow = c(1, 4))
    for (m in mod.x) {
        boxplot(MEslist[[i]]$dataAE[, m] ~ meta[meta$celltype ==
            i, ]$Group, col = c("firebrick4", "lightblue"),
            ylab = "Avg expr of eigen gene", xlab = "",
            boxwex = 0.3, main = paste0(i, " : ", gsub("AE",
                "M", m)))
        boxplot(MEslist[[i]]$dataAE[, m] ~ meta[meta$celltype ==
            i, ]$trt, col = c("firebrick4", "lightblue"),
            ylab = "Avg expr of eigen gene", xlab = "",
            boxwex = 0.3, main = paste0(i, " : ", gsub("AE",
                "M", m)))
    }
}
```

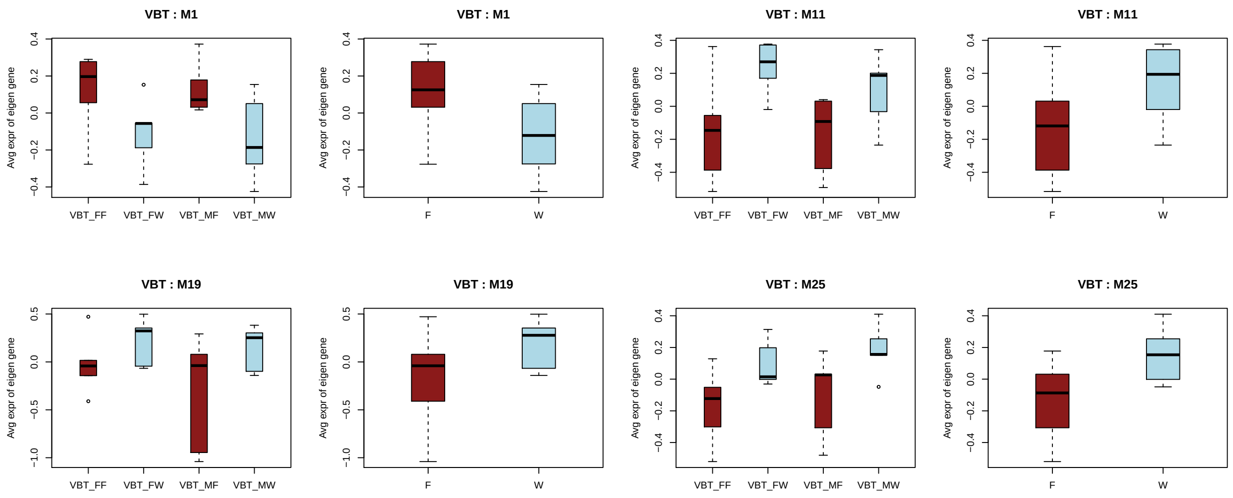

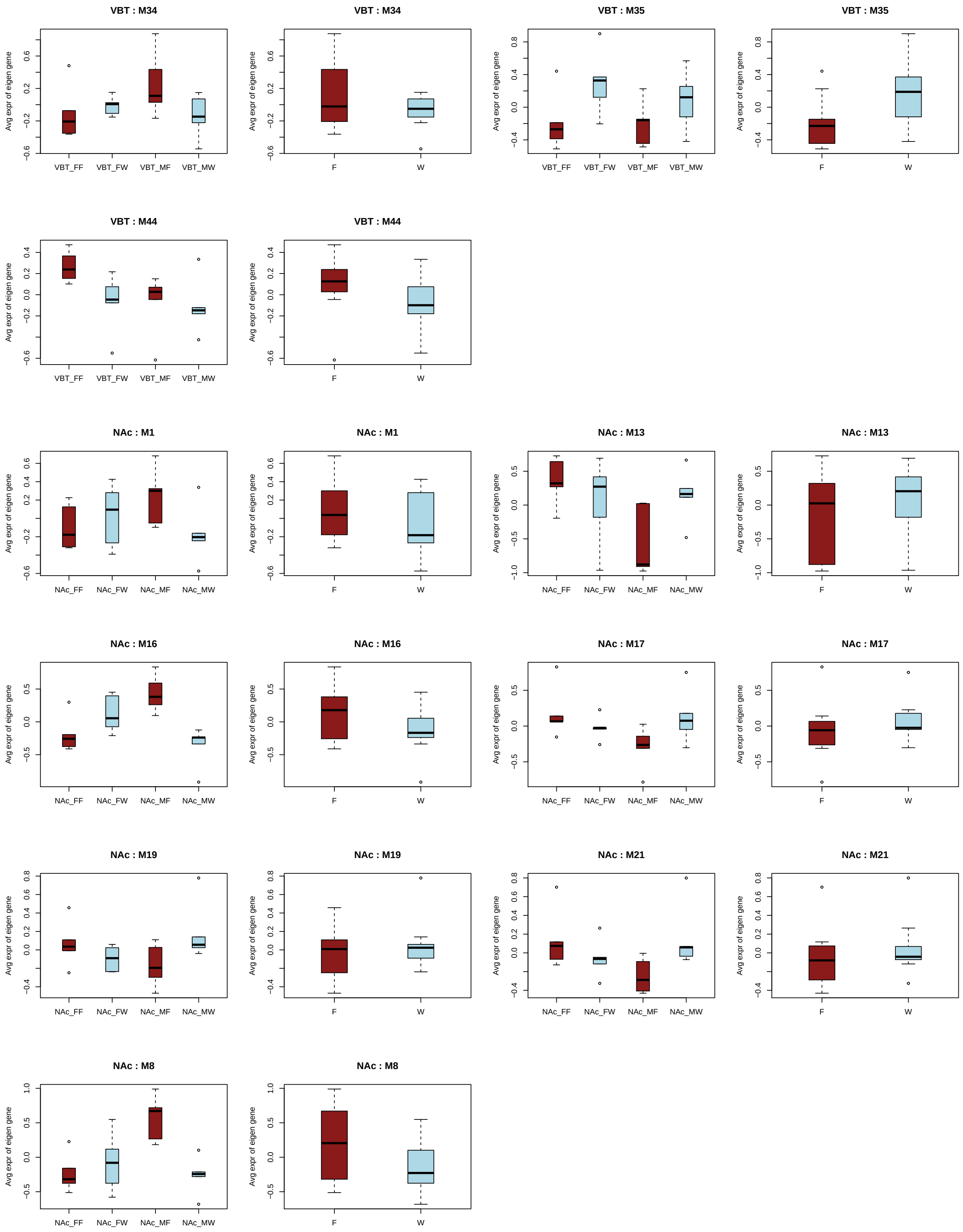
VBT : M34
VBT : M34
VBT : M35
VBT : M35
VBT : M44
VBT : M44
NAc : M1
NAc : M1
NAc : M13
NAc : M13
NAc : M16
NAc : M16
NAc : M17
NAc : M17
NAc : M19
NAc : M19
NAc : M21
NAc : M21
NAc : M8
NAc : M8
Avg expr of eigen gene
VBT_FF
VBT_FW
VBT_MF
VBT_MW
NAc_FF
NAc_FW
NAc_MF
NAc_MW
F
W

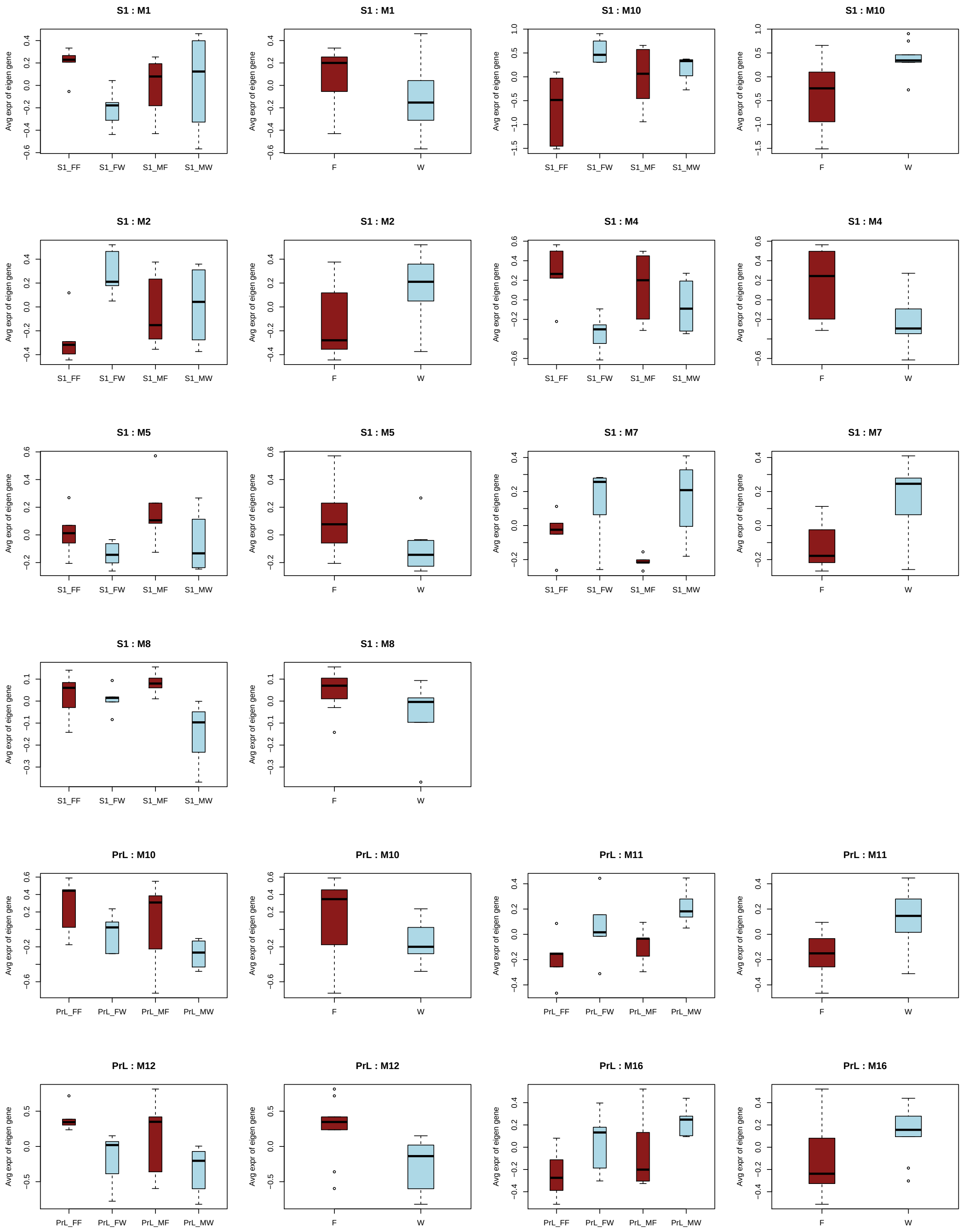
S1 : M1
S1 : M1
S1 : M10
S1 : M10
S1 : M2
S1 : M2
S1 : M4
S1 : M4
S1 : M5
S1 : M5
S1 : M7
S1 : M7
S1 : M8
S1 : M8
PrL : M10
PrL : M10
PrL : M11
PrL : M11
PrL : M12
PrL : M12
PrL : M16
PrL : M16
Avg expr of eigen gene
S1_FF
S1_FW
S1_MF
S1_MW
PrL_FF
PrL_FW
PrL_MF
PrL_MW
F
W

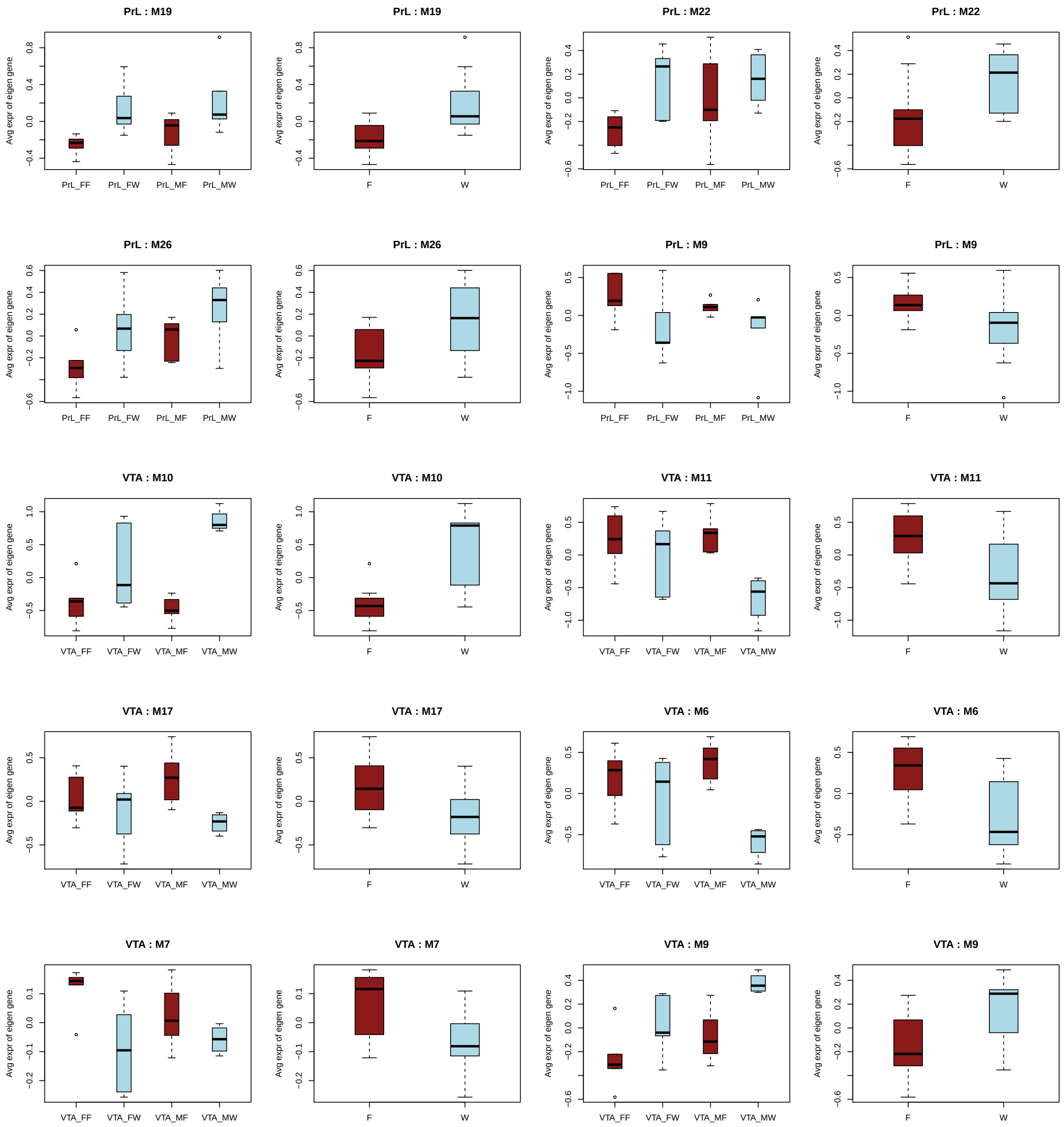


#### Relate Modules to traits : ROAST

Model is build in each module usign the gene expression value.

```
mroast_module <- function(logcpm.x, meta.x, contrast,
    modlist.x) {
    i = which(meta.x$Group %in% contrast)
    meta1 = meta.x$Group[i]
    design = model.matrix(~meta1)
    if (meta1[1] == contrast[2]) {
```

```
        design[, 2] = (!(design[, 2])) * 1
    }
    logcpm1 = logcpm.x[, i]
    x = mroast(logcpm1, modlist.x, design, contrast = 2,
        nrot = 1000, adjust.method = "BH", set.statistic = "msq")
    return(x)
}

modlist = list()
for (j in celltype) {
    x = unique(netcol[[j]])
    x = x[x > 0]
    modlist[[j]] = lapply(1:length(x), function(i) {
        which(rownames(logcpm) %in% names(netcol[[j]][netcol[[j]] ==
            i]))
    })
    names(modlist[[j]]) = paste0("M", seq(length(modlist[[j]])))
}

# design = model.matrix( ~ 0 + meta$Group )
# colnames(design)
# =unlist(strsplit(colnames(design),'meta\\$Group'))[seq(2,40,2)]

roasttrait = list()
for (i in celltype) {
    x = cbind(sex = "Female", mroast_module(logcpm,
        meta, c(paste0(i, "_FF"), paste0(i, "_FW")),
        modlist[[i]]))
    y = cbind(sex = "Male", mroast_module(logcpm, meta,
        c(paste0(i, "_MF"), paste0(i, "_MW")), modlist[[i]]))
    roasttrait[[i]] = rbind(cbind(Module = rownames(x),
        x), cbind(Module = rownames(y), y))
}

sigroasttrait = list()
sigroasttrait = lapply(1:length(roasttrait), function(i) {
    roasttrait[[names(roasttrait)[i]]][which(roasttrait[[names(roasttrait)[i]]]$PValue <=
        0.05), ]
})
names(sigroasttrait) = names(roasttrait)
```

##### PLOT : ROAST results

*Plot the significant cases: Modules for which the genes are significntly differentially expressed (Fentanyl ~ water) for male and female separately.*

```
par(mfrow = c(2, 2))
for (i in celltype) {
    if (nrow(sigroasttrait[[i]]) > 0) {
        x = diag(as.matrix(sigroasttrait[[i]][, paste0("Prop",
            sigroasttrait[[i]][, "Direction"])]))
        name = sigroasttrait[[i]][, "Module"]
        col = labels2colors(sigroasttrait[[i]][, "Direction"],
```

```
            colorSeq = c("slategray1", "lightsalmon"))
        bp = barplot(x, names.arg = name, col = col,
            ylim = c(0, 1), main = i, xlab = "Modules",
            ylab = "Proportion of genes", las = 2)
        text(bp, -0.03, labels = sigroasttrait[[i]]$sex,
            xpd = TRUE, cex = 0.8)
        text(bp, x + 0.04, labels = signif(sigroasttrait[[i]]$PValue,
            2), col = "blue")
        legend("top", fill = c("skyblue", "indianred1"),
            c("Down", "Up"))
    }
}
```

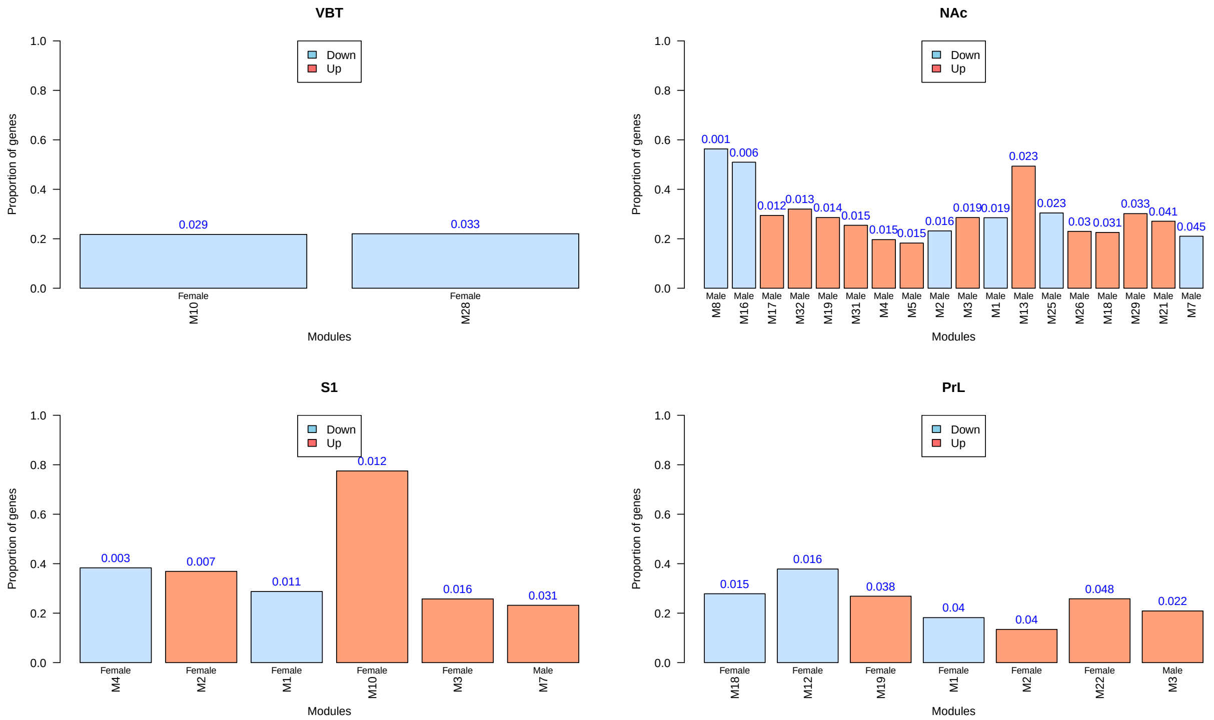

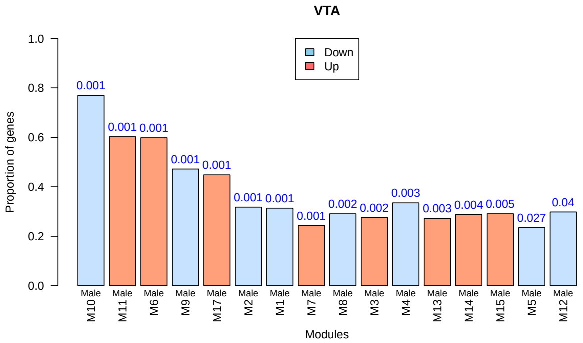


#### Boxplot for significant Modules from ROAST

```
for (i in celltype) {
    par(mfrow = c(1, 4))
    if (nrow(sigroasttrait[[i]]) > 0) {
        name = sigroasttrait[[i]][, "Module"]
        for (k in name) {
            ii = modlist[[i]][[k]]
            jj = which(meta$celltype == i)
            logcpm1 = logcpm[ii, jj]
            boxplot(colMeans(logcpm1) ~ meta[jj, ]$Group,
                col = c("sienna3", "paleturquoise"),
                ylab = "Avg expr of all genes in module",
                xlab = "", boxwex = 0.3, main = paste0(i,
                  " : ", k))
        }
    }
}
```

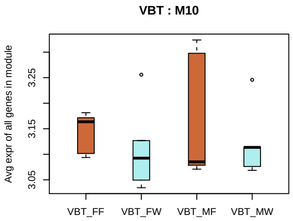


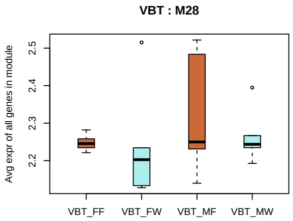

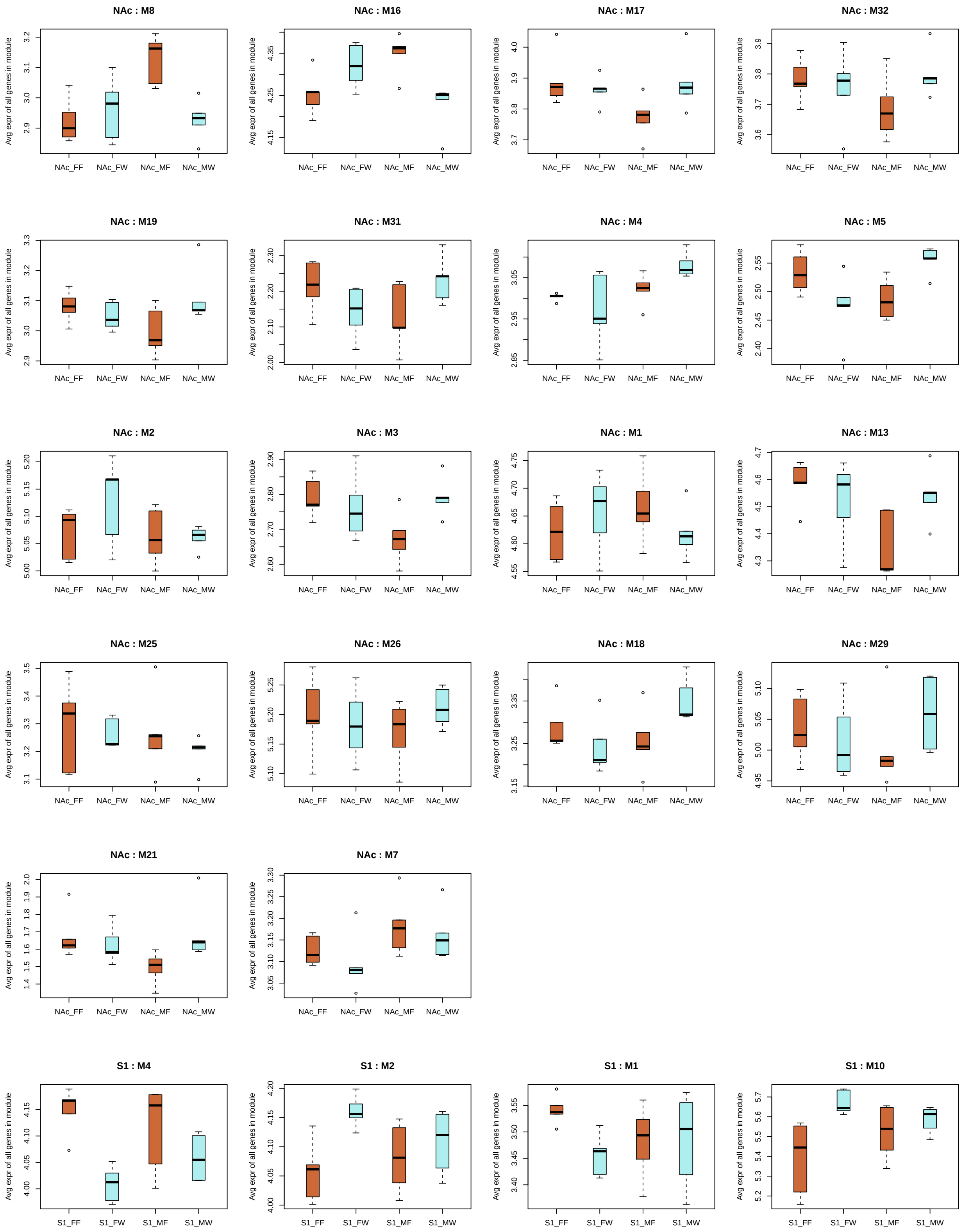
NAc : M8
NAc : M16
NAc : M17
NAc : M32
NAc : M19
NAc : M31
NAc : M4
NAc : M5
NAc : M2
NAc : M3
NAc : M1
NAc : M13
NAc : M25
NAc : M26
NAc : M18
NAc : M29
NAc : M21
NAc : M7
S1 : M4
S1 : M2
S1 : M1
S1 : M10
Avg expr of all genes in module
NAc_FF
NAc_FW
NAc_MF
NAc_MW
S1_FF
S1_FW
S1_MF
S1_MW

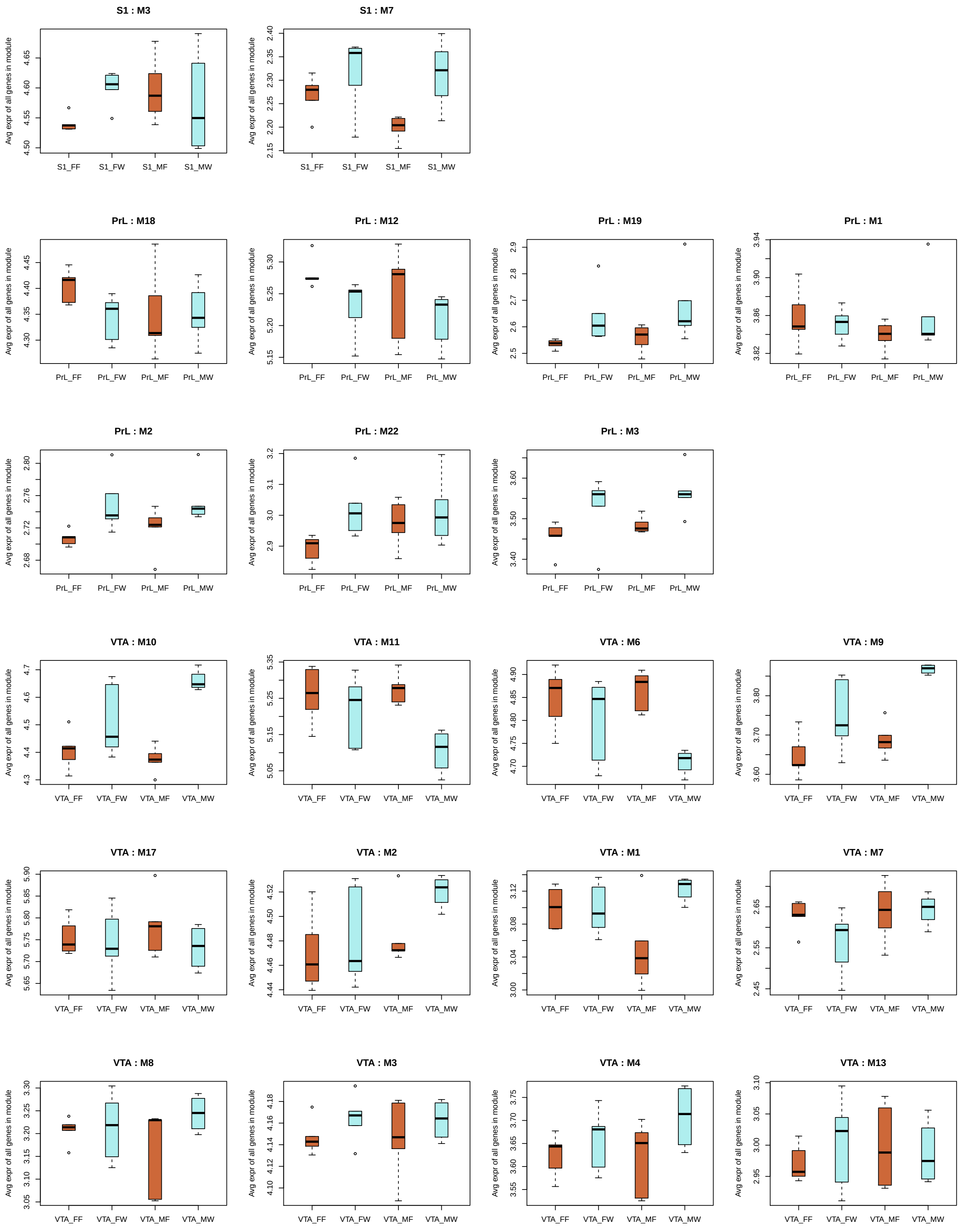
S1 : M3
S1 : M7
PrL : M18
PrL : M12
PrL : M19
PrL : M1
PrL : M2
PrL : M22
PrL : M3
VTA : M10
VTA : M11
VTA : M6
VTA : M9
VTA : M17
VTA : M2
VTA : M1
VTA : M7
VTA : M8
VTA : M3
VTA : M4
VTA : M13
Avg expr of all genes in module
S1_FF
S1_FW
S1_MF
S1_MW
PrL_FF
PrL_FW
PrL_MF
PrL_MW
VTA_FF
VTA_FW
VTA_MF
VTA_MW

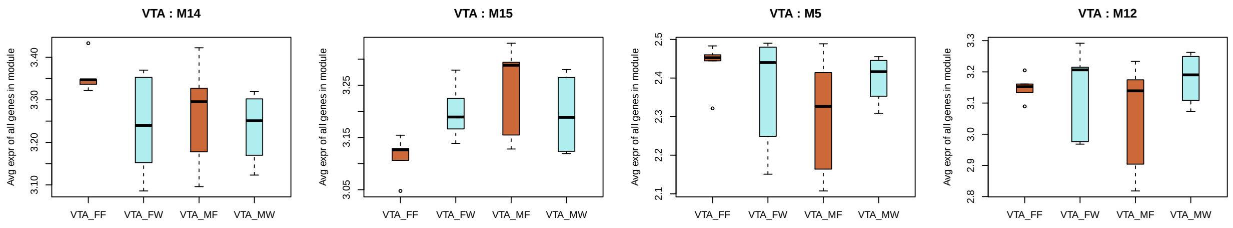


#### Modules genes list and traits

```
library(biomaRt)
geneModuleMembership = list()
for (i in 1:length(celltype)) {
    MEs = moduleEigengenes(multiExpr[[i]]$data, netcol[[celltype[i]]])
    MEs = MEs$averageExpr
    nSamples = nrow(multiExpr[[i]]$data)
    ModuleMembership = as.data.frame(bicor(multiExpr[[i]]$data,
        MEs, use = "p"))
    MMPvalue = as.data.frame(corPvalueStudent(as.matrix(ModuleMembership),
        nSamples))
    names(ModuleMembership) = gsub("AE", paste0(celltype[i],
        ".MM"), names(ModuleMembership))
    names(MMPvalue) = gsub("AE", "p.MM", names(ModuleMembership))
    geneModuleMembership[[celltype[i]]] = list(membership = as.matrix(ModuleMembership),
        pvals = as.matrix(MMPvalue))
}

netGenes = data.frame(gene_id = colnames(datExpr),
    module.PrL = netcol[["PrL"]], module.NAc = netcol[["NAc"]],
    module.VBT = netcol[["VBT"]], module.VTA = netcol[["VTA"]],
    module.S1 = netcol[["S1"]])


mart = useMart("ensembl")
mart = useDataset("mmusculus_gene_ensembl", mart)
ann = getBM(mart = mart, attributes = c("ensembl_gene_id",
    "mgi_symbol", "chromosome_name", "gene_biotype",
    "description"), filters = "ensembl_gene_id", values = colnames(datExpr))
ann = ann[duplicated(ann$ensembl_gene_id) == F, ]

outp = merge(ann, netGenes, by = 1)
for (i in names(geneModuleMembership)) {
    outp = merge(outp, geneModuleMembership[[i]][[1]],
        by.x = 1, by.y = 0)
}

write.table(outp, paste0(workdir, "/gene_summary.txt"),
    sep = "\t")

# Topgenes
top10gene = list()
```

```
for (k in 1:length(celltype)) {
    n = max(netcol[[celltype[k]]])
    top10 = rep(NA, n)
    for (i in 1:n) {
        tmp = outp[outp[, paste0("module.", celltype[k])] ==
            i, ]
        tmp = tmp[order(tmp[, paste0(celltype[k], ".MM",
            i)], decreasing = T)[1:10], "mgi_symbol"]
        tmp = setdiff(tmp, "")
        top10[i] = paste(tmp, collapse = ",")
    }
    top10gene[[celltype[k]]] = data.frame(module = paste0(celltype[k],
        ".AE", 1:n), top10)
}
moduleannolist = list()
for (k in 1:length(celltype)) {
    modanno = cbind(limmatraits[[celltype[k]]]$pval,
        limmatraits[[celltype[k]]]$logFC)
    rownames(modanno) = paste(celltype[k], rownames(modanno),
        sep = ".")
    modanno = merge(modanno, top10gene[[celltype[k]]],
        by.x = 0, by.y = 1)
    nGenes = as.data.frame(table(netcol[[celltype[k]]]))
    nGenes[, 1] = paste(celltype[k], ".AE", nGenes[,
        1], sep = "")
    names(nGenes) = c("module", "nGenes")
    modanno = merge(modanno, nGenes, by = 1)
    modanno = modanno[order(as.numeric(gsub(paste0(celltype[k],
        ".AE"), "", modanno[, 1]))), c(1, 9, 8, 5,
        2, 6, 3, 7, 4)]
    colnames(modanno)[1] = "Module"

    dtroast = roasttrait[[celltype[k]]]
    dtroast$Module = gsub("M", paste0(celltype[k],
        ".AE"), dtroast$Module)

    modanno = merge(modanno, dtroast[dtroast$sex ==
        "Female", ], by.x = 1, by.y = 1)
    modanno = merge(modanno, dtroast[dtroast$sex ==
        "Male", ], by.x = 1, by.y = 1)

    moduleannolist[[celltype[k]]] = modanno
}

wb <- createWorkbook(type = "xlsx")
for (i in 1:length(celltype)) {
    sheet <- createSheet(wb, sheetName = celltype[i])
    xlsx.addHeader(wb, sheet, value = "Module summary")
    xlsx.addLineBreak(sheet, 1)
    xlsx.addHeader(wb, sheet, value = celltype[i])
    xlsx.addLineBreak(sheet, 1)
    xlsx.addTable(wb, sheet, moduleannolist[[celltype[i]]],
        startCol = 2)
```

```
}
saveWorkbook(wb, paste0(workdir, "/Module_summary.xlsx"))
```
