## Supplementary figure and tables for "Transcriptomic profiling of reward and sensory brain areas in perinatal fentanyl exposed juvenile mice"

**Fig. S1**

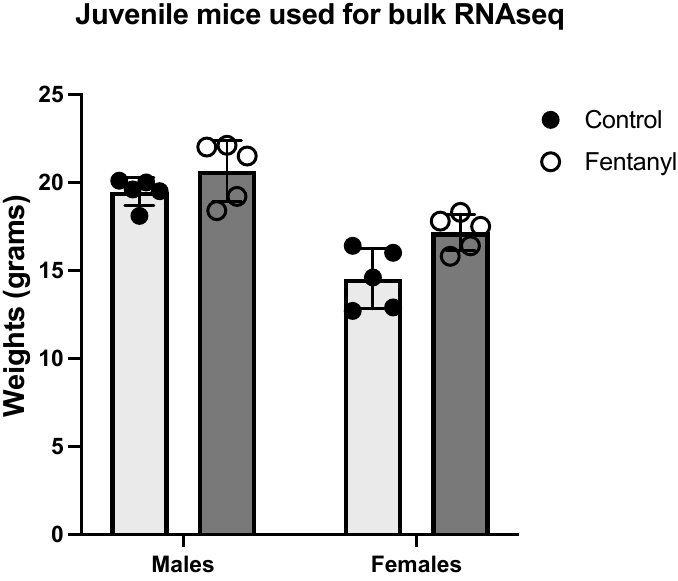

**Supplementary Figure 1. Weights in grams of P35 mice used for RNAsequencing in this study.**

**Fig. S2**

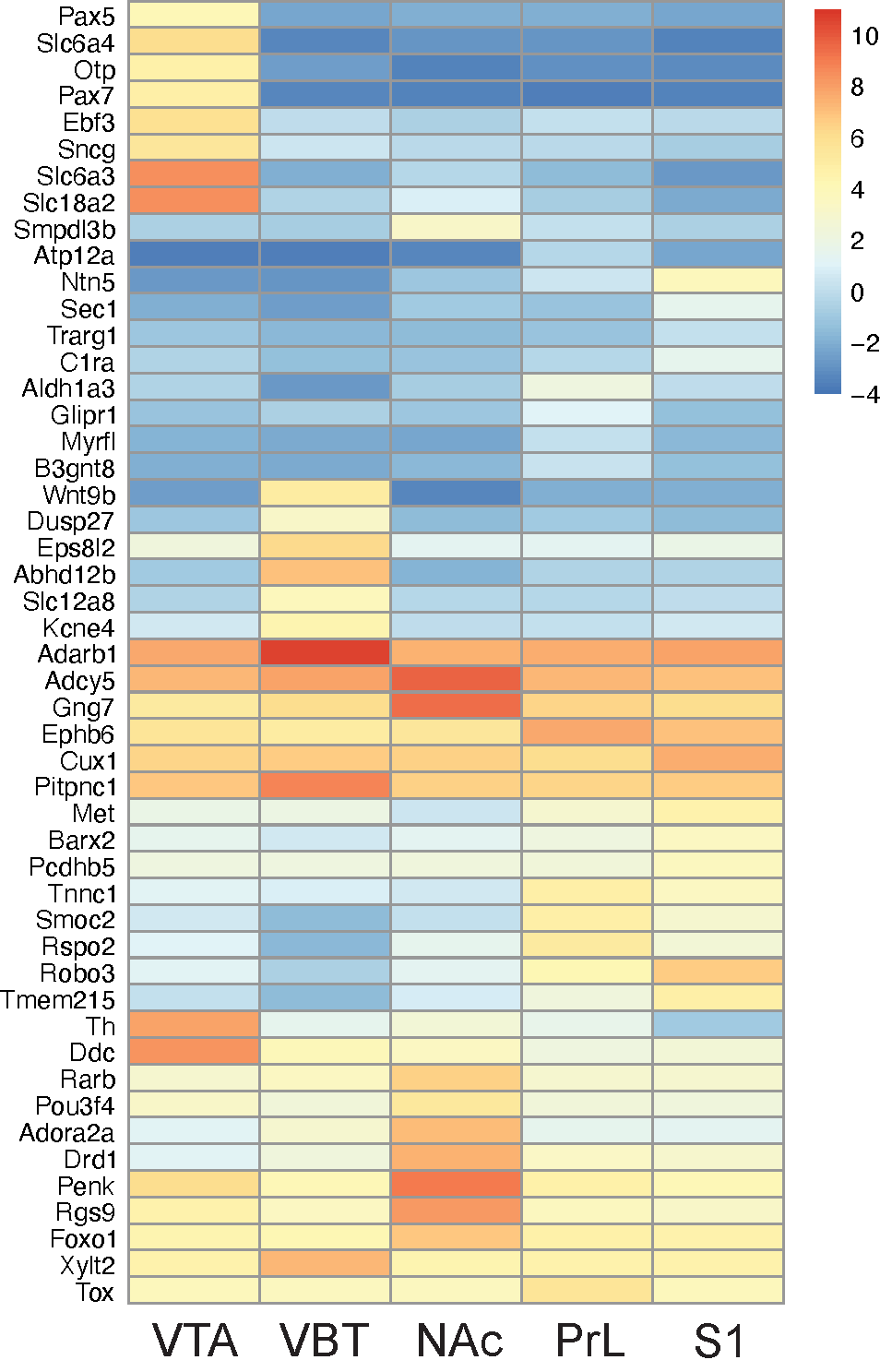

**Supplementary Figure 2. Expression pattern of top DEGs in control brains showing region markers across brain areas.**

**Fig. S3**

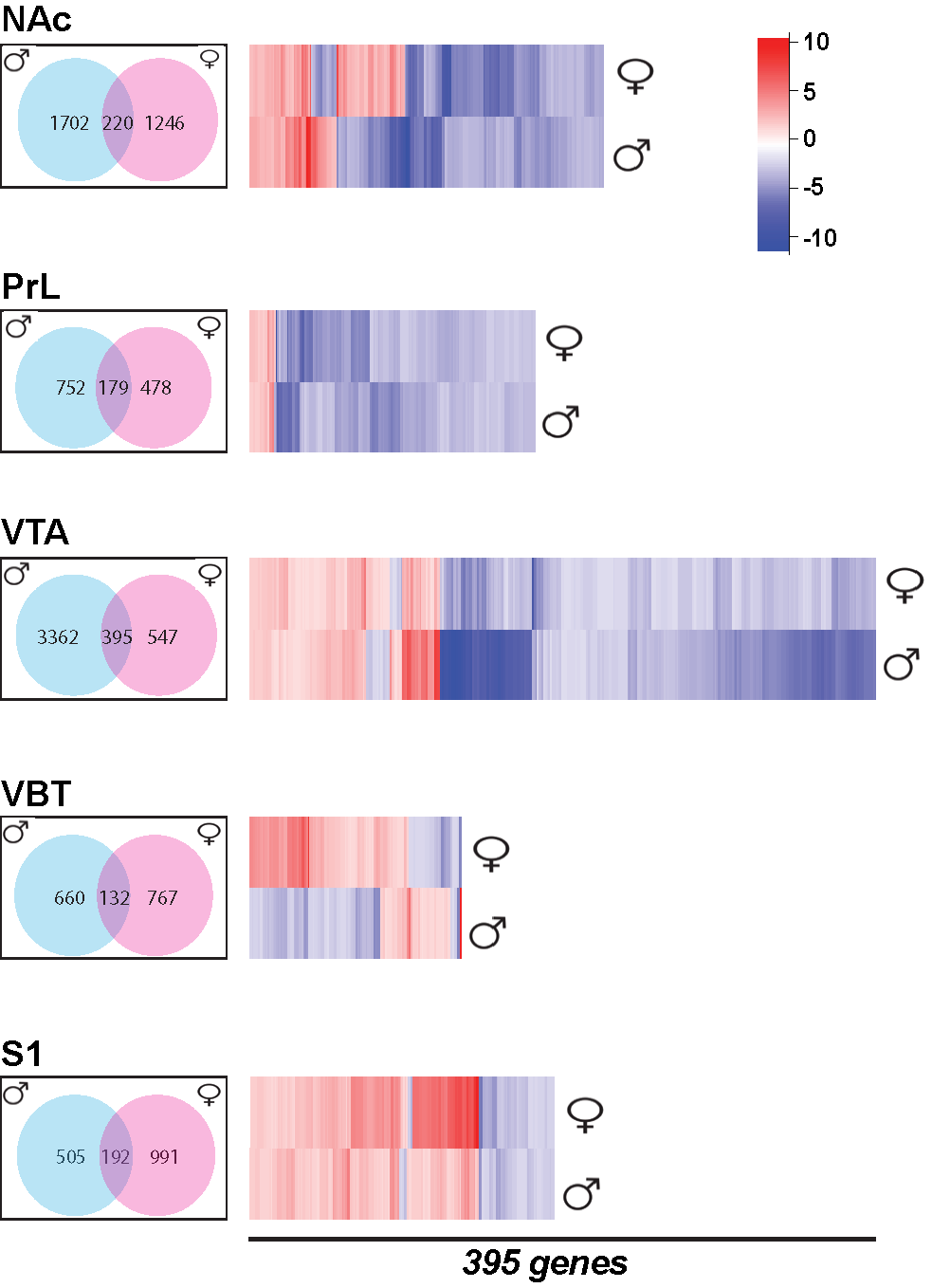

**Supplementary Figure 3.** Venn diagram of trending gene transcripts between males (blue) and females (pink) across brain areas using an uncorrected p < 0.05 and Log Fold Change of +/- 0.3. Heatmaps show expression pattern of common gene transcripts between sexes. The prelimbic (PrL), ventral tegmental area (VTA) and primary somatosensory area (S1) show similar expression pattern of common gene transcripts between sexes however, in the nucleus accumbens (NAc) and ventrobasal thalamus (VBT) many common upregulated female gene transcripts were downregulated in males. This was particularly obvious in the VBT.

**Fig. S4**

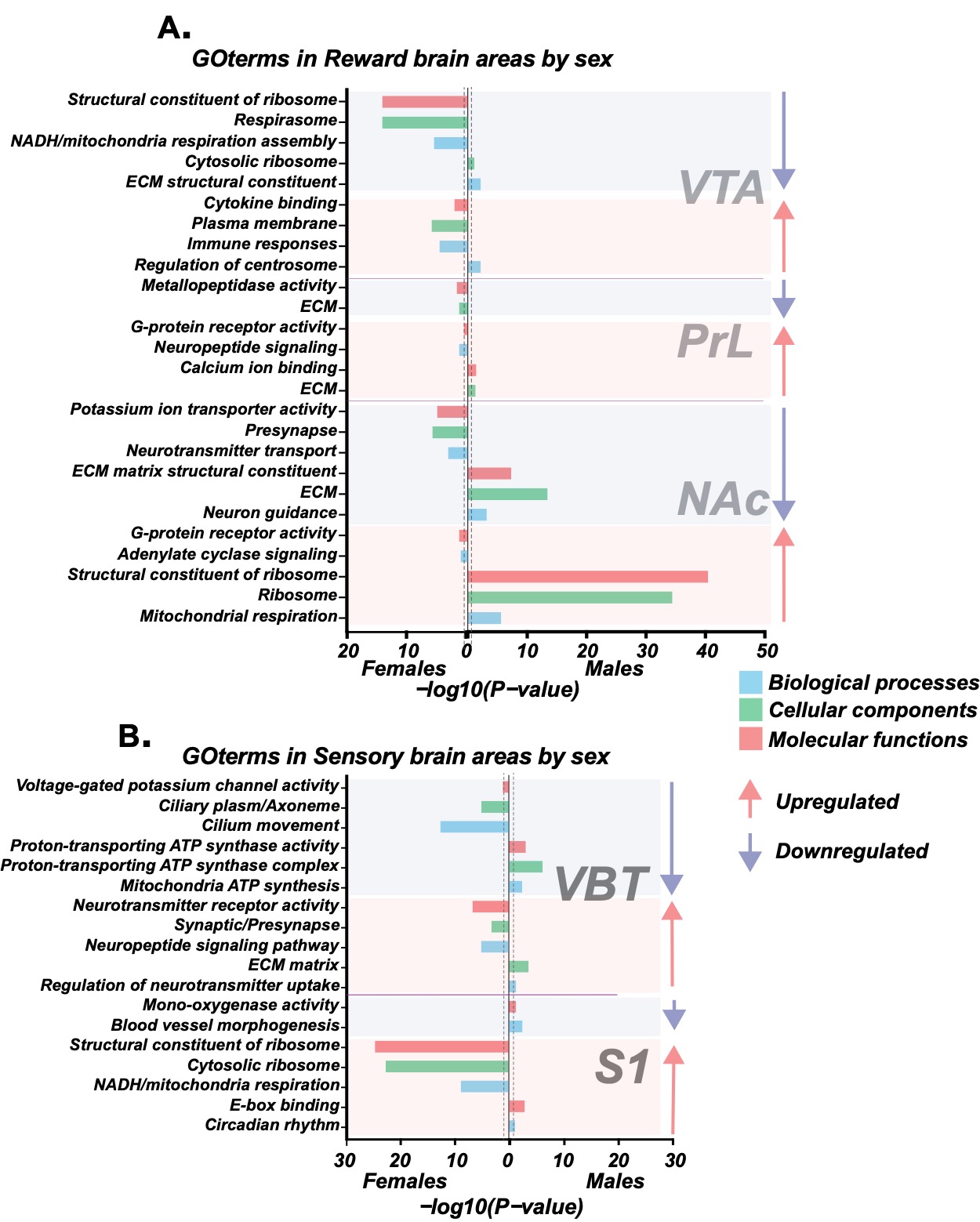

**Supplementary Figure 4. Gene enrichment of reward and sensory brain areas in perinatal fentanyl exposed juvenile mice using uncorrected p<0.05.** (A) and (B) Gene Ontology Enrichment for trending genes across reward and sensory brain areas in a sex-wise manner. Y-axis indicate top enrichment terms and X-axis indicate significance level of enrichment.

**Table S1A.**

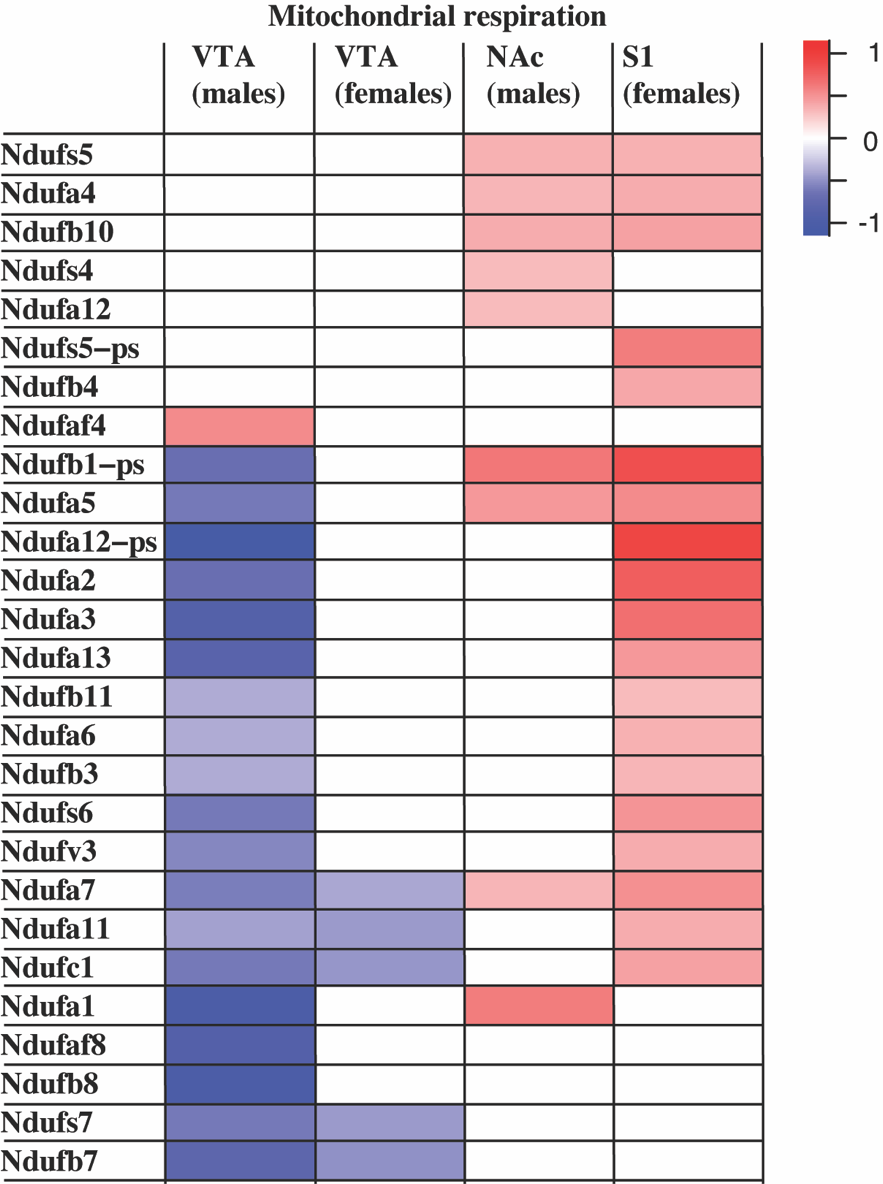

**Table S1A**. Expression profile mitochondrial complex 1 NADH:ubiquinone oxidoreductase subunits in perinatal fentanyl exposure by sex across brain regions. Red values indicate upregulation while blue values indicate downregulation. Significance level was set at FDR < 0.05; LogFC, log fold change.

**Table S1B.**

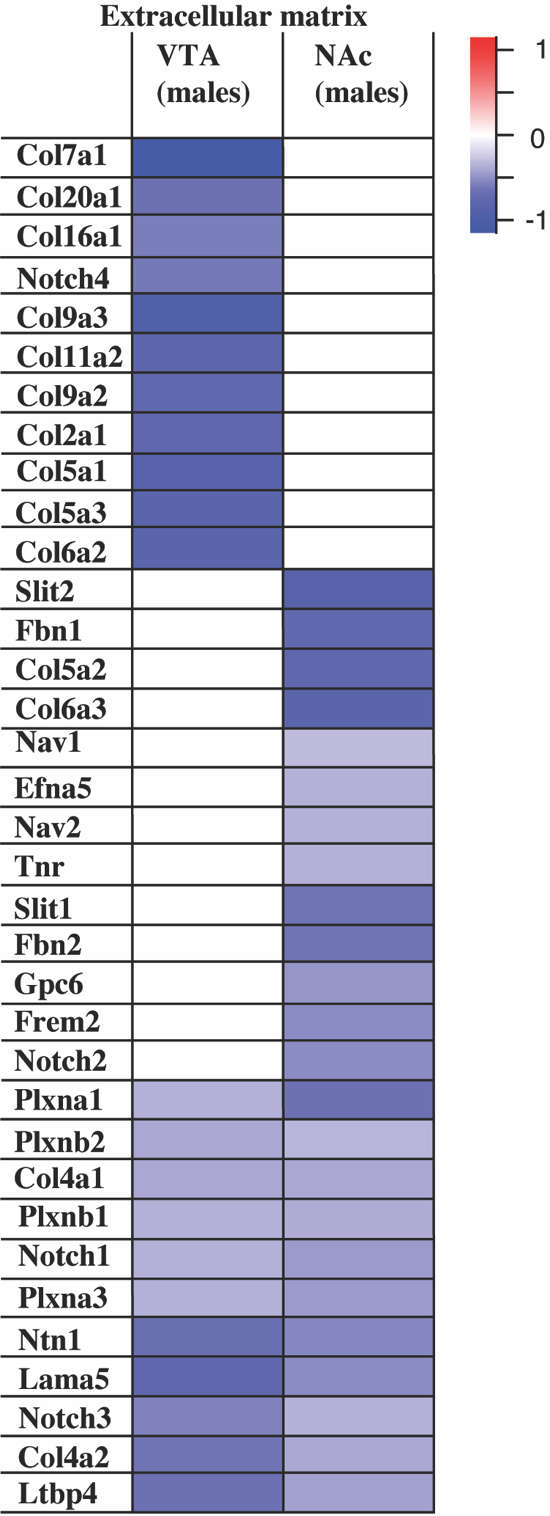

**Table S1B.** Expression profile of ECM signaling molecules and neuronal guidance genes downregulated specifically in the NAc and VTA of male juvenile mice exposed to perinatal fentanyl. Significance level was set at FDR < 0.05.; LogFC, log fold change.

**Table S1C.**

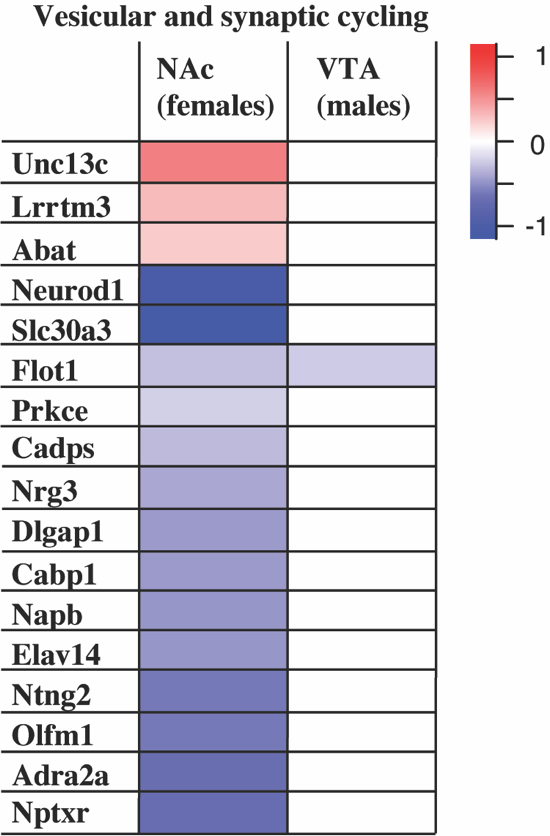

**Table S1C.** Expression profile of some vesicular and synaptic signaling genes specifically dysregulated in NAc of female juvenile mice exposed to perinatal fentanyl. Red values indicate upregulation while blue values indicate downregulation. Significance level was set at FDR < 0.05. LogFC, log fold change.
